# Leveraging biobank-scale rare and common variant analyses to identify *ASPHD1* as the main driver of reproductive traits in the 16p11.2 locus

**DOI:** 10.1101/716415

**Authors:** Katrin Männik, Thomas Arbogast, Maarja Lepamets, Kaido Lepik, Anna Pellaz, Herta Ademi, Zachary A Kupchinsky, Jacob Ellegood, Catia Attanasio, Andrea Messina, Samuel Rotman, Sandra Martin-Brevet, Estelle Dubruc, Jacqueline Chrast, Jason P Lerch, Lily R Qiu, Triin Laisk, The 16p11.2 European Consortium, The Simons VIP Consortium, The eQTLGen Consortium, R Mark Henkelman, Sébastien Jacquemont, Yann Herault, Cecilia M Lindgren, Hedi Peterson, Jean Christophe Stehle, Nicholas Katsanis, Zoltan Kutalik, Serge Nef, Bogdan Draganski, Erica E Davis, Reedik Mägi, Alexandre Reymond

## Abstract

Whereas genome-wide association studies (GWAS) allowed identifying thousands of associations between variants and traits, their success rate in pinpointing causal genes has been disproportionately low. Here, we integrate biobank-scale phenotype data from carriers of a rare copy-number variant (CNV), Mendelian randomization and animal modeling to identify causative genes in a GWAS locus for age at menarche (AaM). We show that the dosage of the 16p11.2 BP4-BP5 interval is correlated positively with AaM in the UK and Estonian biobanks and 16p11.2 clinical cohorts, with a directionally consistent trend for pubertal onset in males. These correlations parallel an increase in reproductive tract disorders in both sexes. In support of these observations, 16p11.2 mouse models display perturbed pubertal onset and structurally altered reproductive organs that track with CNV dose. Further, we report a negative correlation between the 16p11.2 dosage and relative hypothalamic volume in both humans and mice, intimating a perturbation in the gonadotropin-releasing hormone (GnRH) axis. Two independent lines of evidence identified candidate causal genes for AaM; Mendelian randomization and agnostic dosage modulation of each 16p11.2 gene in zebrafish *gnrh3:egfp* models. *ASPHD1*, expressed predominantly in brain and pituitary gland, emerged as a major phenotype driver; and it is subject to modulation by *KCTD13* to exacerbate GnRH neuron phenotype. Together, our data highlight the power of an interdisciplinary approach to elucidate disease etiologies underlying complex traits.

## Introduction

Recent studies suggest that, in addition to common variants, rare variants contribute substantially to population variance of complex traits. In particular, individually rare but collectively common copy-number variants (CNVs) have been shown to account for traits such as education attainment, body mass index (BMI) or height [1–4]. A hallmark example of a rare variant of strong effect on complex traits in humans is the 16p11.2 600 kb BP4-BP5 CNV interval [5]. The deletion (OMIM #611913) and reciprocal duplication (OMIM #614671) of this interval are mediated by positively-selected *Homo sapiens*-specific low-copy repeats [6, 7]. The 16p11.2 CNVs are among the most frequent known genetic causes of mental disorders such as autism spectrum disorders, schizophrenia and cognitive impairment [5, 8–12]. They also impact satiety response and BMI in a dosage-dependent manner [13–15]. Correspondingly, 16p11.2 gene dosage affects head circumference, and specifically, the size of brain structures associated with reward, language and social cognition [9, 13, 15–17]. Genome-wide association studies (GWAS) have identified reproducible associations between common SNPs of the 16p11.2 interval and BMI [18], schizophrenia [19, 20] and age at menarche (AaM) [21–23]. However, altered pubertal timing has not been reported as part of the 16p11.2 CNV phenotype, possibly because of ascertainment strategies biased towards pediatric neurodevelopmental cases in CNV studies.

Whereas GWAS have identified thousands of correlations between common genetic variants and complex traits, the success rate in pinpointing the causal genes has been disproportionately low [24, 25]. One recently proposed solution to this challenge involves intersection of expression quantitative trait loci (eQTL) data with GWAS risk loci [26, 27], a strategy called Mendelian Randomization (MR). This approach aims to resolve gene-trait associations and offer the ability to estimate the strength of the causal effects in GWAS loci [28–30].

Here, we coalesce rare CNV associations in unbiased adult populations and clinical cohorts; phenotyping of 16p11.2 mouse models; MR and single gene *in vivo* zebrafish assays to characterize hitherto unappreciated genetic associations for reproductive traits. Our agnostic approaches identified likely causative genes for impairment of both timing and developmental processes involved in sexual development.

## Materials and methods

Materials and methods are detailed in **Supplementary Methods**

### Study cohorts

The 16p11.2 CNV carriers are defined as individuals who carry the recurrent proximal 16p11.2 BP4-BP5 600 kb hemizygous deletion or heterozygous duplication in the UK biobank (UKBB), the Estonian Genome Center, University of Tartu biobank (EGCUT), the Pan-European and the Simons VIP 16p11.2 cohorts. See **Supplementary Table S1** for summary characteristics and **Supplementary Methods** for cohort descriptions, details on genotyping and association analyses.

### 16p11.2 mouse models

We assessed female and male reproductive parameters including estrous cyclicity, sperm count, anogenital distance, morphology of urogenital tract organs and hypothalamic volume in previously engineered 16p11.2^Del/+^ and 16p11.2^Dup/+^ mouse models [31, 32].

### Brain structural magnetic resonance imaging (MRI)

Human structural MRI data were acquired, processed and analyzed as described [16, 17, 33]. The mass-univariate statistical analysis of whole-brain volume maps was performed in 146 postpubertal individuals as described [34]. Mouse MRI was performed as detailed in [35].

### Disease enrichment analysis

We assessed enrichments by g:Profiler [36] and *MetaCore*^™^ in lists of genes differentially expressed in 16p11.2 CNV carriers and models [37, 38].

### Mendelian randomization analysis

We used univariate [28] and multivariate [39–41] MR to assess causal effects of 16p11.2 genes on regulation of AaM.

### GnRH neuron phenotyping in zebrafish

We performed overexpression and CRISPR/Cas9 genome editing of 16p11.2 genes in zebrafish embryos as described [42, 43]. *In vivo* imaging of neuronal patterning in *gnrh3:egfp* transgenic reporter larvae [44] is described in **Supplementary Methods**.

### Global gene expression similarity

We analyzed expression similarity of identified candidate genes across publicly available human and mouse gene expression datasets using Multi Experiment Matrix (MEM) [45] and funcExplorer [46] tools. Downstream enrichment analyses were conducted using g:Profiler toolset [47].

## Results

Similar to their clinically ascertained counterparts [13], we demonstrated that adult population biobank carriers of the 16p11.2 CNVs present mirror effects on BMI. We observed that genomic dosage of the 16p11.2 interval is associated significantly with BMI in the first release of the UKBB cohort (first UKBB; ANOVA p=1.65×10^−17^). We replicated these results in both the independent second release of the UKBB (second UKBB; ANOVA p=5.37×10^−36^) and the geographically distinct EGCUT (ANOVA p=6.4×10^−08^) cohorts (**Supplementary Results, Table S2** and **Figure S1-S2**). Next, we investigated AaM, an understudied trait in 16p11.2 CNV carriers that was associated previously with common SNPs in the 16p11.2 interval [21–23]. We identified a significant association between 16p11.2 dosage and AaM (corrected ANOVA p=1.93×10^−05^, first UKBB). Compared to controls (n=60,466; μ_AAM_=12.9 years) AaM was significantly earlier in deletion carriers (n=11; μ_AAM_=11.4 years, Δ=-1.5 years; corrected p=0.00098, Wilcoxon test) and delayed in duplication carriers (n=21; μ_AAM_=14.4 years, Δ=+1.5 years; corrected p=0.002, Wilcoxon test) (**Figure 1A**). These differences were independent of ethnicity (**Supplementary Table S3** and **Figure S1-S2**) and remained unchanged after correcting for BMI, year of birth, and population structure. We replicated this association in the second UKBB (corrected p=6.5×10^−04^, ANOVA), EGCUT (corrected p=2.4×10^−05^, ANOVA), and confirmed the mirror effect between deletion and duplication in two independent 16p11.2 clinical cohorts (Simons VIP Collection p=7.7×10^−05^; Pan-European cohort p=0.0034) (**Figure 1BC, Supplementary Results** and **Figure S1**). We identified a similar, but non-significant trend in males possibly due to a less distinctive onset of puberty in this sex. Compared to controls in the first UKBB, the second UKBB and the Simons VIP cohorts, male deletion carriers trended toward “younger than average age” at the onset of pubertal traits (e.g. all ancestries combined, “first facial hair” in first UKBB: OR=2.73, p=0.08; in second UKBB: OR=2.69, p=0.025) and duplication carriers, oppositely, toward “older than average age” (in first UKBB: OR=2.78, p=0.09; in second UKBB: OR=1.58, p=0.32) (**Figure 1DE, Supplementary Results** and **Table S4**). The associations between 16p11.2 CNV and pubertal onset are accompanied by increased diagnoses of the female reproductive tract disorders in the EGCUT cohort, e.g. “absent, scanty and rare menstruation” (N91 in WHO ICD-10; OR=4.4, p=0.013, Fisher’s Exact Test) or “noninflammatory disorders of ovary, fallopian tube and broad ligament” (N83; OR=5.2, p=0.003, Fisher’s Exact Test). Altogether 50% and 64% of deletion and duplication EGCUT females were diagnosed with irregular/absent menstruation, disorders of hormonal or ovarian dysfunction, respectively (**Figure 1F, Supplementary Results** and **Table S5**). We analogously observed that pediatric male duplication carriers in the Simons VIP cohort had an increase in undescended testicles (13.2%, n=5) and hypospadias (5.3%, n=2) compared to published rates [48, 49]. Upon inclusion of other genital problems, 11 out of 38 (29%) boys with 16p11.2 duplication were affected.

**Figure 1.**
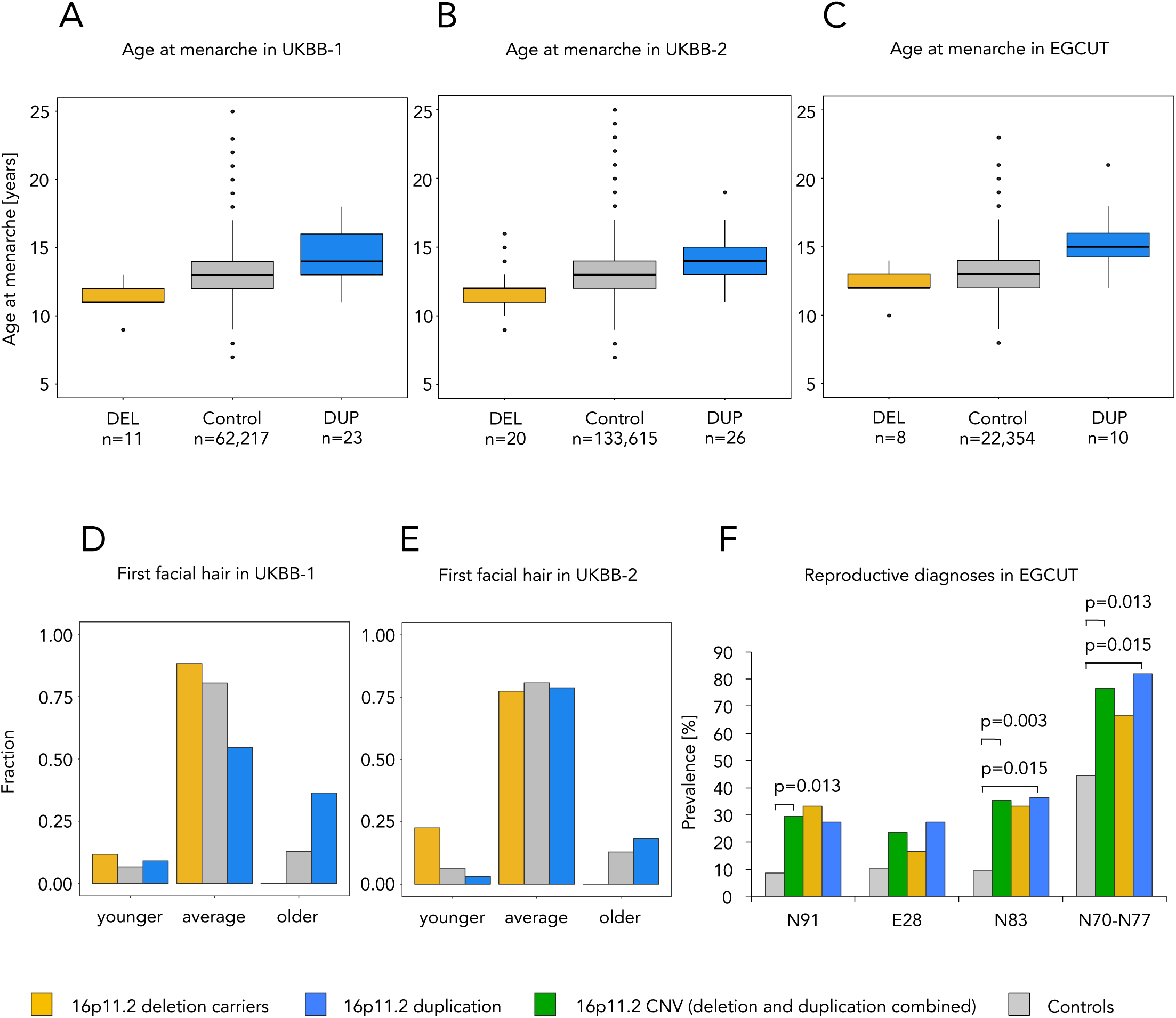
Reproductive traits of 16p11.2 CNV carriers from unselected adult population cohorts. Yellow, blue and green depict carriers of 16p11.2 deletion, duplication and CNV (deletion and duplication combined), respectively, while controls are shown in grey. Mirror association with age at menarche in individuals of European ancestry (**A**) the first UKBB cohort (UKBB-1), (**B**) the second UKBB cohort (UKBB-2), and (**C**) the EGCUT cohort. Directionally consistent non-significant trend towards altered timing of male pubertal traits, e.g. self-reporting of “first facial hair” at a “younger”, “average” or “older” age than contemporaries in (**D**) the first UKBB, and in (**E**) the second UKBB cohorts. (**F**) Frequency of selected reproductive diagnoses in female 16p11.2 CNV carriers in the EGCUT cohort. Diagnosed diseases are presented according to the ICD-10 codes, N91 “absent, scanty and rare menstruation”, E28 “ovarian dysfunction”, N83 “noninflammatory disorders of ovary, fallopian tube and broad ligament”, N70-77 “inflammatory diseases of female pelvic organs”. AaM: Age at menarche; DEL: 16p11.2 deletion; DUP: 16p11.2 duplication

Next, we asked if these genetic associations were paralleled by molecular alterations. We analyzed published transcriptome profiles and found that genes correlated with 16p11.2 CNV dosage in lymphoblastoid cell lines (LCLs) derived from 16p11.2 CNV carriers and cerebral cortices of syntenic mouse models [37, 38] were enriched for genes associated with urogenital dysfunctions (LCL: 8 of 25 top-ranked terms, FDR <1.43×10^−06^; cortex: 9 of 25, FDR <1.75×10^−12^) (**Supplementary Results** and **Table S6**). These results prompted us to assess the effect of 16p11.2 dosage on reproductive traits in mouse models. Consistent with previous reports [7, 31] the 16p11.2^Del/+^ and 16p11.2^Dup/+^ animals showed a mirror effect on body weight that is opposite to that of human 16p11.2 CNV carriers. By recording timing of first ovulation in females, we noted a reciprocal effect on sexual maturation that, similar to weight [31, 32], was inverted compared to humans. The 16p11.2^Dup/+^ and 16p11.2^Del/+^ mice reached, respectively, first ovulation two days earlier (p=0.0021, Breslow-Wilcoxon test) and five days later (p=4.8×10^−06^, Breslow-Wilcoxon test) than their wild-type littermates (**Figure 2AB**). These results were complemented by perturbation of time spent in individual stages of the cycle in 16p11.2^Del/+^ females (estrus: p=0.034, diestrus: p=0.029; Student’s t-test; **Supplementary Results**) and gross morphological changes of the reproductive organs in both 16p11.2^Dup/+^ females and males. 16p11.2^Dup/+^ females had an increased uterine weight compared to estrous cycle-harmonized control females (p=0.04, Student’s t-test, normalized for body weight; **Figure 2C** and **Supplementary Results**). Additionally, 16p11.2^Dup/+^ males presented a significant reduction of the ano-genital distance compared to controls (p=0.004, Student’s t-test, normalized for body weight; **Figure 2D** and **Supplementary Results**), a commonly used endpoint for hormonally regulated disorders of sexual differentiation in rodents and humans, and an indicator of abnormal reproductive tract masculinization [50]. The histological architecture of both 16p11.2^Del/+^ and 16p11.2^Dup/+^ mice post pubertal testis was normal with no major changes in seminiferous tubules organization although testes from 3 out of 5 16p11.2^Dup/+^ mice display some seminiferous tubules with spermatogenic defects. These include tubules with germ cell degeneration and one case of tubular atrophy with vacuolization that however did not affect dramatically sperm count compared to controls (**Figure 2E** and **Supplementary Results)**.

**Figure 2.**
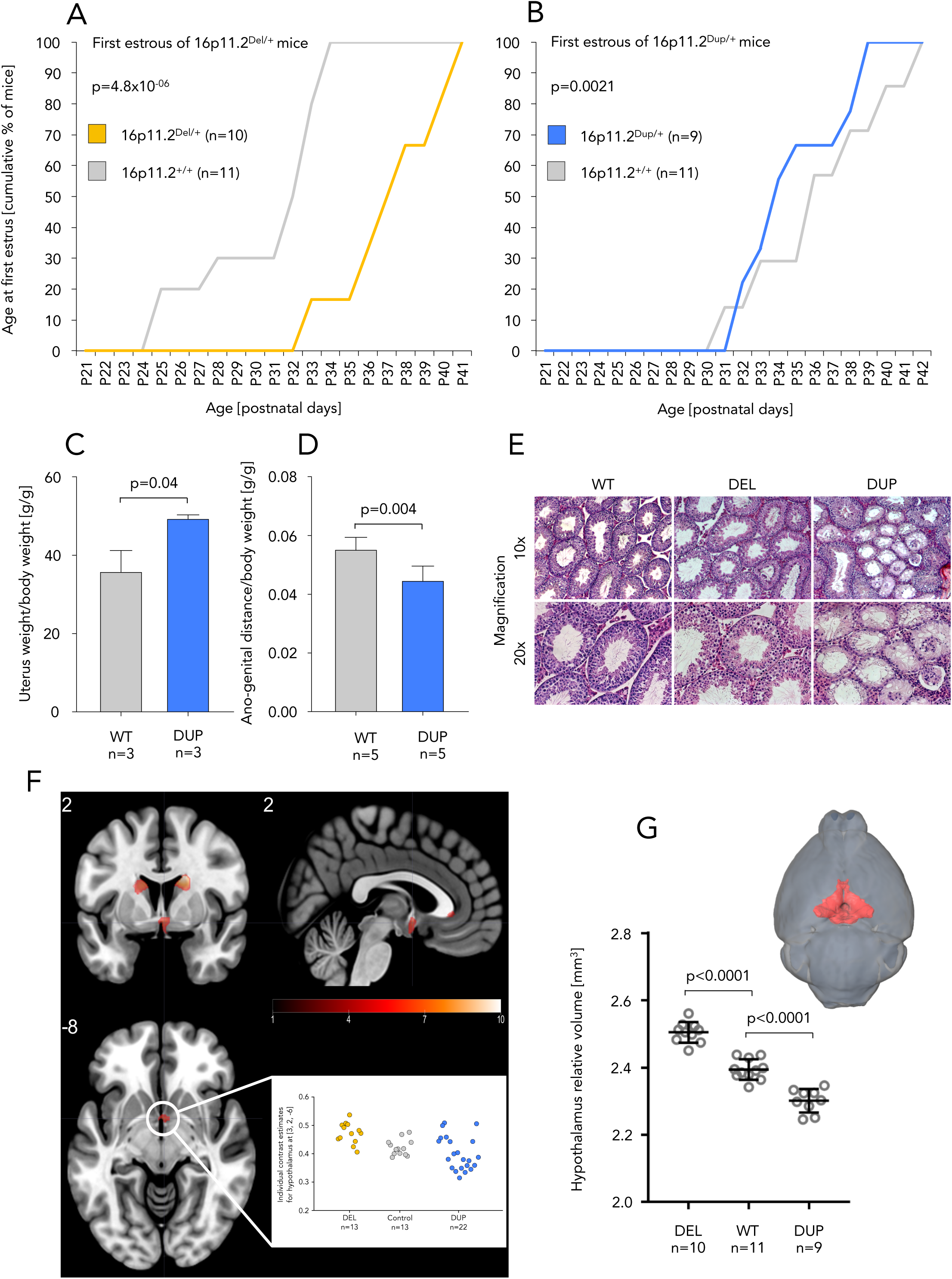
Reproductive traits of 16p11.2 mouse models and hypothalamic volume in 16p11.2 mice and human patients. Color code as in **Figure 1**. Altered age at first estrous of (**A**) 16p11.2^Del/+^ and (**B**) 16p11.2^Dup/+^ female mice compared to wild-type littermates. (**C**) Significantly increased size of uterus in 16p11.2^Dup/+^ female and (**D**) shortened ano-genital distance in 16p11.2^Dup/+^ male mice. (**E**) The architecture of seminiferous tubules shows regions with abnormal histology in 3 out of 5 analyzed 16p11.2^Dup/+^ males, specifically tubules with germ cell degeneration and tubular atrophy with vacuolization. Significant negative correlation between 16p11.2 dosage and the adjusted hypothalamic volume in human and mouse female individuals. (**F**) Differential brain structure changes associated with age of menarche in 16p11.2 CNV carriers. Color-coded representation of statistical parametric maps (SPMs) of interaction analysis between “group” (DEL vs DUP vs CTR) and AaM. These SPMs are displayed on an average human brain in standardized space using a heat scale (left panel). For visualization purposes SPMs are thresholded at p<0.001 (uncorrected for multiple comparisons). Group differences of hypothalamus volume (pFWE<0.05), contrast estimates and 90% confidence interval are plotted for local maxima within the hypothalamus. (**G**) Relative volume of the hypothalamus normalized by total brain volume in 16p11.2 mice. The data are represented as the mean ± standard deviation; Tukey’s test was applied following a significant one-way ANOVA. A MRI acquisition of a mouse brain with the hypothalamus highlighted in red is shown in the top-right inset. AaM: Age at menarche; DEL: 16p11.2 deletion; DUP: 16p11.2 duplication; CTR: non-CNV carrier controls; WT: wild-type animals

Sexual development in mammals is regulated by hormonal interplay between hypothalamus, pituitary gland and gonads. We used human MRI data [16, 17, 33] to assess possible 16p11.2 dosage-dependent brain anatomy changes. Given our strong *a priori* hypothesis of an impaired neuroendocrine axis, we restricted the search volume to a 5 mm sphere centered on the hypothalamic region. We found in female CNV carriers, but not in male, a significant negative correlation (p_FWE_<0.05) between the 16p11.2 CNV dosage and the hypothalamic volume adjusted for individuals’ total intracranial volume as proxy for head size (**Figure 2F** and **Supplementary Table S7**). Consistent with human data, we replicated previously reported abnormal hypothalamic morphology in 16p11.2^Del/+^ mouse models [32, 51]. We observed a significant negative correlation between 16p11.2 dosage and hypothalamus volume relative to the total brain volume in female mice. The deletion and duplication animals have, respectively, larger and smaller relative hypothalamic regions than wild type animals (both p<0.0001, one-way ANOVA, post-hoc Tukey; **Figure 2G** and **Supplementary Table S8**). These results suggest that volumetric perturbations of the hypothalamus, a structure comprised of GnRH secreting neurons essential for initiating puberty and activating the hypothalamus–pituitary–gonadal axis, could contribute to the 16p11.2-linked alterations of sexual development.

To determine which genes in the 16p11.2 interval (**Figure 3A**) are causally relevant for AaM, we performed two-sample uni-[28] and multivariate [39, 40] MR using GWAS summary statistics for AaM [22] and an eQTL meta-analysis of 14,115 whole-blood samples [52]. We obtained support for causality for four genes, two of which were identified by both analyses, *INO80E* (uni: b=0.098, SE=0.015, p=1.3×10^−10^; multi: b=0.071, SE=0.018, p=9.3×10^−5^) and *KCTD13* (uni: b=-0.154, SE=0.034, p=4.5×10^−6^; multi: b=-0.074, SE=0.023, p=9.7×10^−4^) (**Figure 3BC; Supplementary Table S9** and **Figure S3**). However, less than half (12 out of 28) of the unique 16p11.2 interval genes could be tested, while the rest had insufficient valid instruments. Independently, we modulated the dosage of all 16p11.2 genes agnostically in a *Tg(gnrh3:egfp)* transgenic zebrafish model (**Supplementary Figure S4**) [44, 53] and monitored GnRH3 neuron patterning *in vivo* using an automated imaging platform [54, 55]. The processes governing GnRH axis establishment in zebrafish are highly conserved with mammals. Importantly, *gnrh3* neurons in zebrafish perform hypophysiotropic roles in control of neurogenesis, neuronal migration and reproduction [56]. To model the 16p11.2 duplication, we evaluated the consequences of overexpressing each human mRNA on relative GnRH3 reporter neuron area in the dorsal aspect (n=6,700 larvae; **Figure 4A**). We measured the area of GFP (green fluorescent protein)-positive cells at 5 days post fertilization (dpf) and found that overexpression of only a single transcript, *ASPHD1* (that was not testable by MR), was sufficient to reduce GFP signal significantly in comparison with controls (19% reduction; p<0.0001, one-way ANOVA, post-hoc Tukey) (**Figure 4BC; Supplementary Results** and **Table S10**). To model *ASPHD1* hemizygosity, we depleted endogenous *asphd1* zebrafish transcripts by CRISPR/Cas9 genome editing and quantified the *gnrh3:egfp* neuron area of F0 (founder) mutant larval batches at 5 dpf. We found that reduced expression of *asphd1* also triggered a significant decrease of GFP signal in F0 versus controls (13% reduction, p=0.0031, one-way ANOVA, post-hoc Tukey; **Figure 4BD; Supplementary Results** and **Table S10**). We have shown previously that genes within the 16p11.2 interval can interact to induce additive or epistatic phenotypes [35, 42, 43, 57]. To investigate whether other genes in the 16p11.2 region could also exacerbate or mitigate the *ASPHD1* effect on the developing reproductive axis, we halved the dose of *ASPHD1* and co-injected *KCTD13, INO80E, MAPK3* or *YPEL3*, the genes that showed causal evidence in MR. We found that the larvae co-injected with *ASPHD1* and *KCTD13* showed a significantly aggravated phenotype (compared to controls: 24% reduction, p<0.0001; compared to *ASPHD1* mono-injected batches: 14% reduction, p=0.003, one-way ANOVA, post-hoc Tukey) suggesting genetic interaction between these two genes (**Figure 4E; Supplementary Results** and **Table S11**).

**Figure 3.**
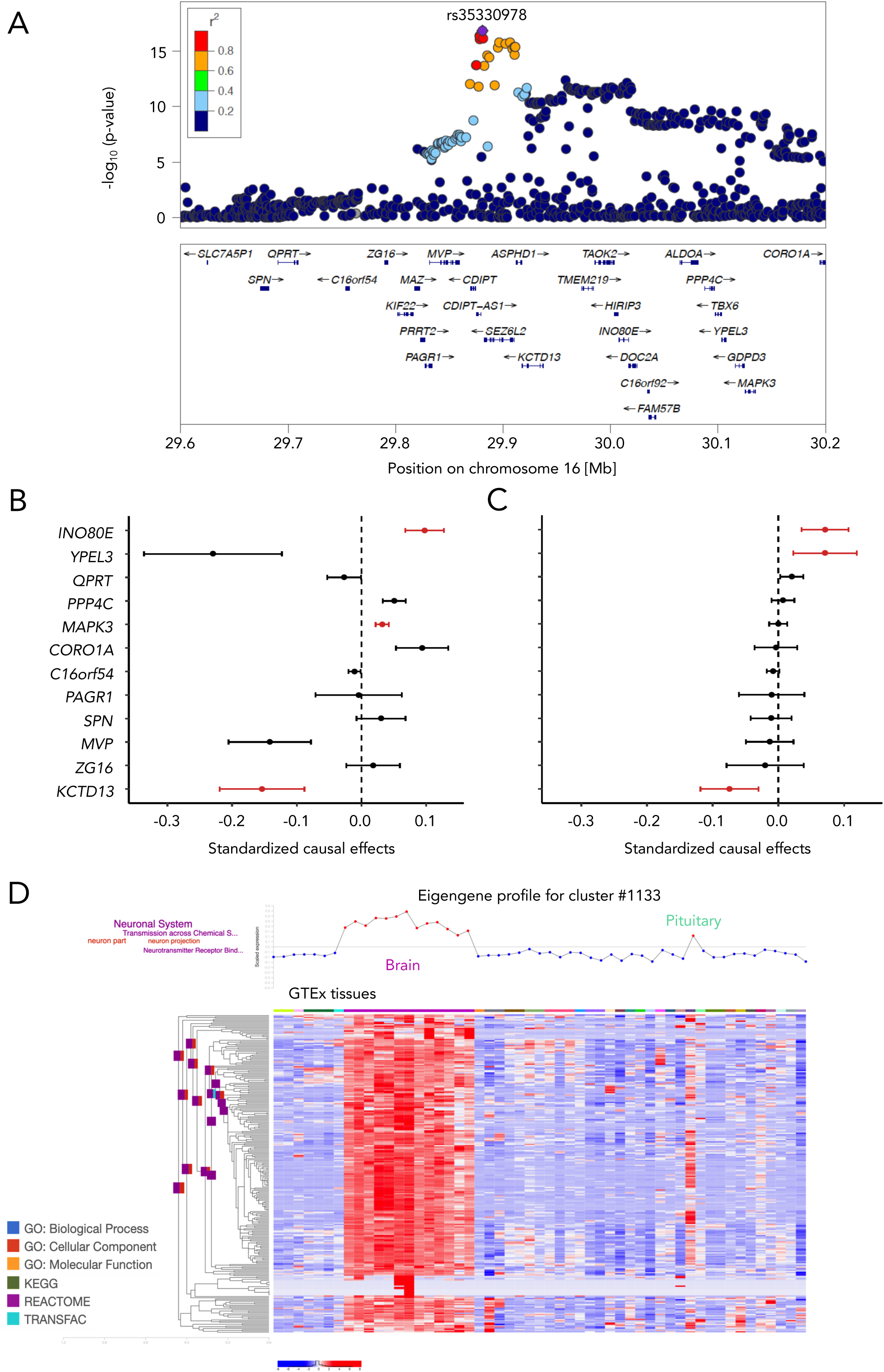
AaM candidate genes in the 16p11.2 interval. (**A**) GWAS peak for age at menarche relative to the CNV genes in the 16p11.2 interval calculated with LocusZoom using data from [22]. (**B**) Univariate and (**C**) multivariate Mendelian randomization analyses showing standardized causal effect estimates for AaM with 95% confidence intervals. Results in red pass the Bonferroni-corrected significant threshold (P < 0.05/12) and in case of univariate analysis, also the HEIDI test (P_HEIDI_ > 0.009). Effects of genes *INO80E* and *KCTD13* on AaM are consistent in both analyses. (**D**) Expression data of 38 GTEx human tissues, clustered using automatic enrichment analysis by funcExplorer, revealed that *ASPHD1* and *CELF4* belong to the same cluster. The cluster #1133 consists of 258 co-expressed genes that show activation mostly in brain regions and pituitary. This is shown by the eigengene profile characterizing the full cluster, and heatmap of these 258 genes across 38 tissues. The cluster is enriched with Gene Ontology and Reactome terms related to neuronal system, neuron projection and neurotransmission (left).

**Figure 4.**
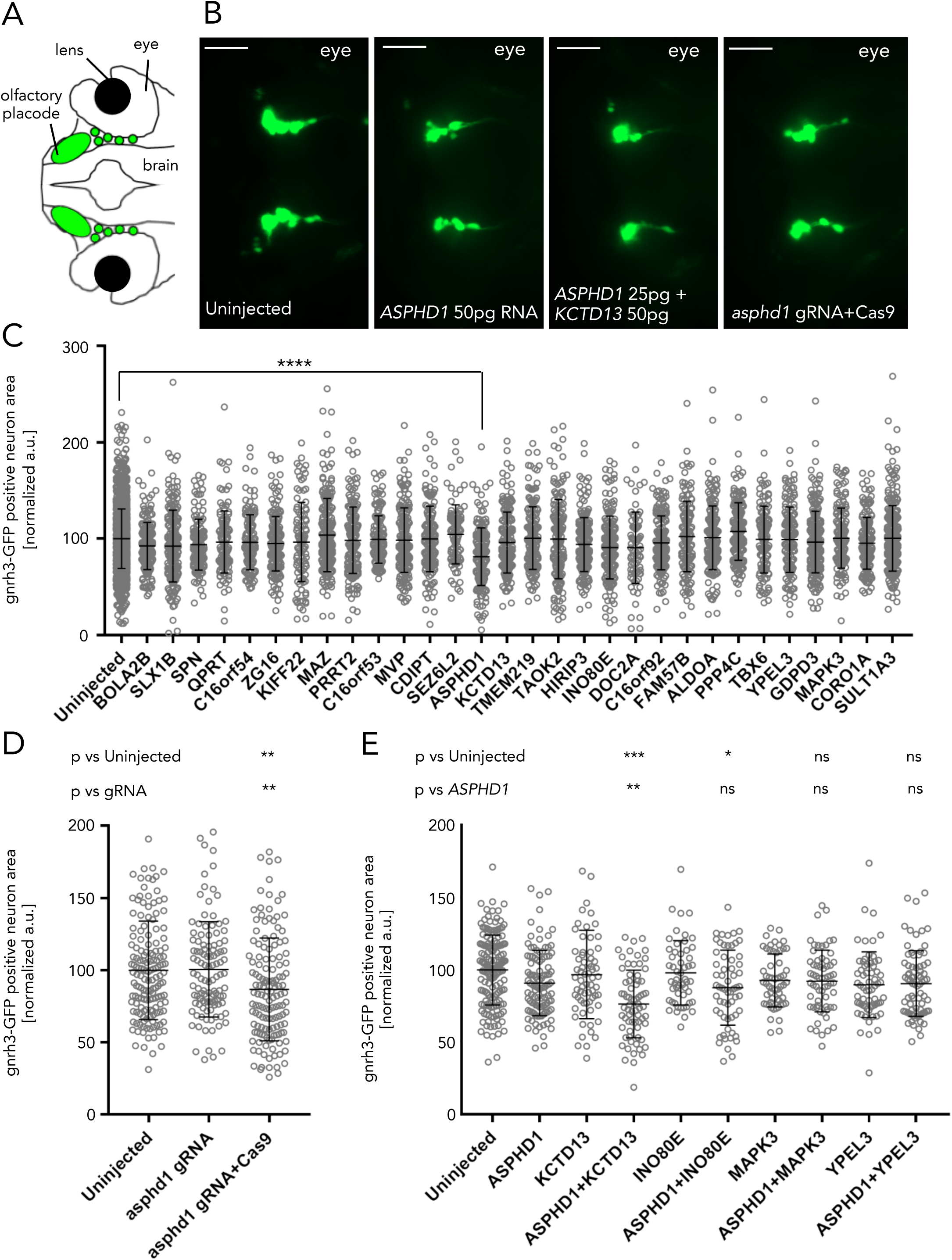
Agnostic gene dosage modulation in a zebrafish model of GnRH axis development. (**A**) Schematic dorsal representation of a *gnrh3:egfp* transgenic larva at 5 dpf, with *gnrh3* expressing neuron highlighted in green. (**B**) From left to right representative dorsal views of GFP signal in Tg(*gnrh3:egfp*) larvae uninjected, injected with *ASPHD1, ASPHD1* and *KCTD13*, asphd1 guide RNA (gRNA) and Cas9 at 5 dpf; scale bar 50 µm. Quantitative assessment of GFP signal in *gnrh3:egfp* larvae injected with human mRNAs coding for genes mapping to the 16p11.2 BP4-BP5 interval. (**C**) *ASPHD1* mRNA induced a significant reduction in GFP signal compared to controls. Dosage: 12.5 pg for *KIF22* and *PPP4C*; 50 pg for all other genes. See **Supplementary Table S10** for larvae numbers. (**D**) F0 mutant larvae injected with *asphd1* gRNA display reduction in GFP signal. Dosage: 100 pg *asphd1* gRNA and 200 pg Cas9 protein. Larvae numbers: Uninjected (n=153), *asphd1* gRNA (n=111), *asphd1* gRNA+Cas9 (n=138). (**E**) Co-injection of *ASPHD1* mRNA with the transcripts prioritized by Mendelian randomization (*KCTD13, INO80E, MAPK3*, and *YPEL3*) identified epistasis between *ASPHD1* and *KCTD13*. Dosage: 25 pg for *ASPHD1*; 50 pg for all other genes. See **Supplementary Table S11** for larvae numbers. The data are represented as the mean ± standard deviation; ns, not significant; *p < 0.05, **p < 0.01, ****p < 0.0001. Tukey’s test was applied following a significant one-way ANOVA.

*ASPHD1* encodes aspartate beta-hydroxylase domain containing 1, a protein of relatively unknown function. We used global gene expression similarity analysis in human and mouse to gain insight into its biological role. We found that *ASPHD1* is co-expressed with genes related to synaptic vesicle function, in particular, with *SEZ6L2*, its distal physical neighbor on 16p11.2; and with *CELF4* a transcript encompassing SNPs associated with neuroticism and depression [58] (**Figure S5**; the results are available at **http://bit.ly/asphd1hsapiens** and **http://bit.ly/asphd1mmusculus**). The three genes share specific expression patterns in multiple brain regions and pituitary (GTEx data set of 38 human tissues), with *ASPHD1* and *CELF4* belonging to the same cluster of 258 co-regulated genes enriched for neuronal processes (“neuron projection” GO:0043005, p=2.16×10^−09^; “transmission across chemical synapses” REAC:112315, p=1.04×10^−10^; “neurotransmitter receptor binding and downstream transmission in the postsynaptic cell” REAC:112314, p=3.7×10^−09^) (**Figure 3D** and **Figure S6**; g:profiler results are available at **https://biit.cs.ut.ee/gplink/l/AOYgmspNQu**).

## Discussion

Accumulating evidence suggests that common and rare variation act additively to influence complex traits [59, 60]. Here, we report a model to leverage information from both rare and common variants to understand the biological basis of reproduction, a multifactorial phenotype.

Despite numerous studies on the 16p11.2 CNVs in clinical cohorts, involvement in the reproductive axis was mostly overlooked with the exception of a reported enrichment of 16p11.2 BP4-BP5 deletions in Müllerian aplasia patients [61, 62]. Here, we show that the 16p11.2 dosage in humans and mice is associated significantly with pubertal onset, reproductive traits and hypothalamus volume. Thus, we add to the repertoire of previously reported links between these CNVs and mental disorders, BMI, head circumference and brain size [8, 9, 12, 13, 15, 16, 31, 32]. The hitherto underappreciation of sexual development phenotypes may have resulted from an ascertainment bias, wherein phenotypes presenting with early onset and the greatest medical management challenges were prioritized. Notably, our findings are consistent with a convergent theme of endocrine, metabolic and behavioral phenotypes in rare CNV carriers and common variant associations in the 16p11.2 locus [18, 22, 63].

Akin to GWAS loci, CNVs present a defined interval associated to phenotype, but often offer no distinct causative genes [64]. Whereas MR has become a popular tool for post-GWAS prioritization of causally relevant genes, authors of these initial studies have refrained from functional validation of their findings *de facto* acknowledging that insufficient eQTL data from the target tissues might be an insurmountable limitation [28, 30]. Here, we exploited the benefits of large-scale MR and overcame its tissue-specific limitation by agnostically assessing the effect of single gene dosage alteration on patterning of GnRH-expressing neurons in the context of zebrafish development. We demonstrate the power of combining *in silico* and *in vivo* methods by identifying *ASPHD1* and *KCTD13* as the driver and modifier of the 16p11.2 reproductive axis, respectively.

*KCTD13* was shown previously to act on neurogenesis through RhoA signaling; reduced KCTD13 levels result in concomitantly reduced synaptic transmission [42, 65]. Although the function of *ASPHD1* is unknown, its expression is restricted to the brain and pituitary gland (https://gtexportal.org) lending support to the above-cited tissue-specific shortcomings of *in silico* causality predictions. Further, the spatially constrained expression pattern of *ASPHD1* (**Figure 3D**) strengthens the tentative link to the neuroendocrine axis that governs pubertal timing. We show that *ASPHD1* is co-expressed with genes involved in neurotransmission and synaptic vesicle location and function. In particular, *CELF4* is involved in neurodevelopmental regulation [66] and is associated with educational attainment [67], neuroticism and mood related alterations [68–70]. Notably, the potential contribution of *ASPHD1* to reproductive function via a putative role in synaptic vesicle integrity is consistent with the biological roles of some Mendelian GnRH deficiency disorder genes. For example, recessive mutations in *DMXL2*, encoding the synaptic vesicle protein rabconnectin-3α, cause delayed puberty due to a reduction of GnRH neurons in the hypothalamus and attenuated kisspeptin responsiveness [71, 72]. Further studies will be required to elucidate the molecular roles and interaction partners of *ASPHD1.*

Two other genes residing at 16p11.2, *TBX6* and *MAZ*, were associated previously with congenital defects of kidney and urinary tract [73, 74], while GWAS have suggested *TBX6* [23], *MAPK3* [21] and *INO80E* [22] as potential candidates for AaM. Furthermore, the 14-gene cluster (distal to proximal: *SPN, QPRT, ZG16, KIF22, PRRT2, MAZ, MVP, SEZ6L2, ASPHD1, KCTD13, TMEM219, TAOK2, INO80E, DOC2A*) was shown to be under coordinated estrogen-mediated regulation [75]. Together, these data offer independent support of the genetic complexity underlying phenotypes in the 16p11.2 locus and reinforce a likely oligogenic etiology with primary drivers and multiple modifiers governing the variance in associated traits [42, 57, 76, 77].

In conclusion, our study illustrates how characterization of traits associated with rare variants in unselected adult populations can provide unbiased insight into disease etiology. We demonstrate further the power of an interdisciplinary approach to pinpoint candidate genes and underlying biological processes in GWAS loci for complex traits.

## Acknowledgements

We express our gratitude to EGCUT participants. We thank the EGCUT personnel for their assistance in recruiting, phenotyping, routing samples, genotyping and administrative responsibilities especially Lili Milani, Viljo Soo, Kairit Mikkel and Mari-Liis Tammesoo. We thank 16p11.2 Pan-European enrollees and their families for their contribution to this study. We are grateful to all of the families at the participating Simons Variation in Individuals Project (Simons VIP) sites, as well as the Simons VIP Consortium. We appreciate obtaining access to SNP genotype, phenotype and imaging data on SFARI Base. Approved researchers can obtain the Simons VIP population dataset described in this study by applying at https://base.sfari.org. The research in this paper has been carried out using the UK Biobank resource (application 17085). Data analyses of this work were carried out in part in the High Performance Computing Center of University of Tartu. The human brain imaging was carried out on the MRI platform of the Département des Neurosciences Cliniques, Centre Hospitalier Universitaire Vaudois, which is generously supported by the Roger De Spoelberch and Partridge Foundations. We thank Yonathan Zohar (University of Maryland) for providing the *gnrh3:egfp* transgenic zebrafish line.

This work was supported by grants from the Swiss National Science Foundation (31003A-143914 to ZK; 31003A_160203 and 31003A_182632 to AR; 32003B_159780 to BD; PP00P3_144902 to SJ), the Horizon2020 Twinning project ePerMed (692145 to AR); the Jacobs Foundation (to KM); the Jérôme Lejeune Foundation (to CA and AR); Estonian Research Council grants IUT20-60, IUT24-6, and PUTJD726 (to TL); European Union through the European Regional Development Fund Project No. 2014-2020.4.01.15-0012 GENTRANSMED and 2014-2020.4.01.16-0125; the Leenaards Foundation (to BD); and US NIH grants P50HD028138 (to NK and EED), R01MH106826 (to EED) and R01HD096326 (to NK). CA is recipient of a Pro-Women Scholarship from the Faculty of Biology and Medicine, University of Lausanne. SJ is a recipient of a Canada Research Chair in neurodevelopmental disorders, and a chair from the Jeanne et Jean Louis Levesque Foundation.

The funders had no role in study design, data collection and analysis, decision to publish, or preparation of the manuscript.

## Author contributions

KM, EED, RM and AR designed the study, supervised individual stages and contributed to data interpretation. KM, ML, KL, TL, CML and RM prepared and analyzed human cohorts data. AP, HA, CA, AM, SR, ED, JC, YH, JCS, SN, KM and AR provided the rodent models, contributed to the husbandry and phenotyping. SMB and BD performed the human MRI, and JE, JPL, LRQ and RMH the mouse MRI analysis. ML, KL and ZK performed the Mendelian randomization. TA, ZAK, NC and EED provided the zebrafish lines and performed experiments. HP and KM analyzed gene expression data. The 16p11.2 European and Simons VIP Consortia members provided phenotype information of European and Northern American patients, respectively. The eQTLGen Consortium members provided whole-blood eQTL meta-analysis data. KM and AR wrote the manuscript with contributions from TA, ML, KL, AP, HA, HP, SN, BD, EED and RM. All authors read and approved the final manuscript.

