## Supplementary Materials for "Leveraging biobank-scale rare and common variant analyses to identify *ASPHD1* as the main driver of reproductive traits in the 16p11.2 locus"

*Männik et al.*

#### European 16p11.2 Consortium - Author Information

François-Guillaume Debray[1], Sebastien Jacquemont[2,3], Joris Andrieux[4], Dominique Bonneau[5], Delphine Heron[6], Sandra Fert-Ferrer[7], Marion Gerard[8], Agnes Guichet[5], Olivier Guillin[9], Bertrand Isidor[10], Didier Lacombe[11], Cédric LeCaignec[10], James Lespinasse[7], Cyril Mignot[6], Ghislaine Plessis[8], Caroline Rooryck-Thambo[11], Mieke van Haelst[12], Catia Attanasio[13], Jacques S Beckmann[2], Manon Cevey-Macherel[2], Jacqueline Chrast[13], Bogdan Draganski[14], Giuliana Giannuzzi[13], Anne M Maillard[2], Sandra Martin-Brevet[14], Aurelie Pain[2], Anna Pellaz[13], Katrin Männik[13,15], Alexandre Reymond[13]

[1]Centre Maladies Métaboliques, Centre Hospitalier Universitaire de Liège, Liège, Belgium

[2]Lausanne University Hospital and University of Lausanne, Lausanne, Switzerland

[3]Centre Hospitalier Universitaire Sainte-Justine Research Center, Montréal, Canada

[4]Centre Hospitalier Régional Universitaire de Lille, Hôpital Jeanne de Flandre, Lille, France

[5]Service de génétique, Centre Hospitalier Universitaire d'Angers, Angers, France

[6]Department of Genetics, Pitié-Salpêtrière Hospital, Paris, France

[7]UF de Génétique Chromosomique, Centre Hospitalier Métropole Savoie, Site Chambéry, Chambéry, France

[8]Centre Hospitalier Universitaire de Caen, Hôpital Clémenceau, Caen, France

[9]Service Hospitalo-Universitaire, Centre Hospitalier du Rouvray, Sotteville-lès-Rouen, France

[10]Centre Hospitalier Universitaire de Nantes, Nantes, France

[11]Centre Hospitalier Universitaire de Bordeaux, Groupe Hospitalier Pellegrin, Bordeaux, France

[12]Department of Clinical Genetics, Amsterdam University Medical Center and University of Amsterdam, Amsterdam, The Netherlands

[13]Center for Integrative Genomics, University of Lausanne, Lausanne, Switzerland

[14]LREN, Department of Clinical Neuroscience, Lausanne University Hospital and University of Lausanne, Lausanne, Switzerland
[15]Estonian Genome Center, Institute of Genomics, University of Tartu, Tartu, Estonia

**eQTLGen Consortium – Author information**

Mawussé Agbessi[1], Habibul Ahsan[2], Isabel Alves[1], Anand Andiappan[3], Wibowo Arindrarto[4], Philip Awadalla[1], Alexis Battle[5,6], Frank Beutner[7], Marc Jan Bonder[8,9,10], Dorret Boomsma[11], Mark Christiansen[12], Annique Claringbould[8], Patrick Deelen[8,13], Tõnu Esko[14], Marie-Julie Favé[1], Lude Franke[8], Timothy Frayling[15], Sina A. Gharib[16,12], Gregory Gibson[17], Bastiaan T. Heijmans[4], Gibran Hemani[18], Rick Jansen[19], Mika Kähönen[20], Anette Kalnapenkis[14], Silva Kasela[14], Johannes Kettunen[21], Yungil Kim[6,22], Holger Kirsten[23], Peter Kovacs[24], Knut Krohn[25], Jaanika Kronberg-Guzman[14], Viktorija Kukushkina[14], Zoltan Kutalik[26], Bernett Lee[3], Terho Lehtimäki[27], Markus Loeffler[23], Urko M. Marigorta[17], Hailang Mei[4], Lili Milani[14], Grant W. Montgomery[28], Martina Müller-Nurasyid[29,30,31], Matthias Nauck[32], Michel Nivard[11], Brenda Penninx[19], Markus Perola[33], Natalia Pervjakova[14], Brandon L. Pierce[2], Joseph Powell[34], Holger Prokisch[35,36], Bruce M. Psaty[12,37,38], Olli T. Raitakari[39], Samuli Ripatti[40], Olaf Rotzschke[3], Sina Rüeger[26], Ashis Saha[6], Markus Scholz[23], Katharina Schramm[31,32], Ilkka Seppälä[27], Eline P. Slagboom[4], Coen D.A. Stehouwer[41], Michael Stumvoll[42], Patrick Sullivan[43], Peter A.C. ‘t Hoen[44], Alexander Teumer[45], Joachim Thiery[46], Lin Tong[2], Anke Tönjes[42], Jenny van Dongen[11], Maarten van Iterson[4], Joyce van Meurs[47], Jan H. Veldink[48], Joost Verlouw[47], Peter M. Visscher[28], Uwe Völker[49], Urmo Võsa[8,14], Harm-Jan Westra[8], Cisca Wijmenga[8], Hanieh Yaghootkar[15], Jian Yang[28,50], Biao Zeng[17], Futao Zhang[28]

Author list is ordered alphabetically.

[1]Computational Biology, Ontario Institute for Cancer Research, Toronto, Canada
[2]Department of Public Health Sciences, University of Chicago, Chicago, United States of America
[3]Singapore Immunology Network, Agency for Science, Technology and Research, Singapore, Singapore

[4]Department of Biomedical Data Sciences, Leiden University Medical Center, Leiden, The Netherlands
[5]Department of Biomedical Engineering, Johns Hopkins University, Baltimore, United States of America
[6]Department of Computer Science, Johns Hopkins University, Baltimore, United States of America
[7]Heart Center Leipzig, Universität Leipzig, Leipzig, Germany
[8]Department of Genetics, University Medical Centre Groningen, Groningen, The Netherlands
[9]European Molecular Biology Laboratory, European Bioinformatics Institute, Wellcome Genome Campus, Hinxton, United Kingdom
[10]Genome Biology Unit, European Molecular Biology Laboratory, Heidelberg, Germany
[11]Department of Biological Psychology, Vrije Universiteit Amsterdam, Amsterdam, The Netherlands
[12]Cardiovascular Health Research Unit, University of Washington, Seattle, United States of America
[13]Genomics Coordination Center, University Medical Centre Groningen, Groningen, The Netherlands
[14]Estonian Genome Center, Institute of Genomics, University of Tartu, Tartu 51010, Estonia
[15]Exeter Medical School, University of Exeter, Exeter, United Kingdom
[16]Department of Medicine, University of Washington, Seattle, United States of America
[17]School of Biological Sciences, Georgia Tech, Atlanta, United States of America
[18]MRC Integrative Epidemiology Unit, University of Bristol, Bristol, United Kingdom
[19]Department of Psychiatry and Amsterdam Neuroscience, Amsterdam UMC, Vrije Universiteit, Amsterdam, the Netherlands
[20]Department of Clinical Physiology, Tampere University Hospital and Faculty of Medicine and Health Technology, Tampere University, Tampere, Finland
[21]Centre for Life Course Health Research, University of Oulu, Oulu, Finland
[22]Genetics and Genomic Science Department, Icahn School of Medicine at Mount Sinai, New York, United States of America
[23]Institut für Medizinische Informatik, Statistik und Epidemiologie, LIFE – Leipzig Research Center for Civilization Diseases, Universität Leipzig, Leipzig, Germany

[24]IFB Adiposity Diseases, Universität Leipzig, Leipzig, Germany
[25]Interdisciplinary Center for Clinical Research, Faculty of Medicine, Universität Leipzig, Leipzig, Germany
[26]Institute of Social and Preventive Medicine, Lausanne University Hospital, Lausanne, Switzerland
[27]Department of Clinical Chemistry, Fimlab Laboratories and Finnish Cardiovascular Research Center-Tampere, Faculty of Medicine and Health Technology, Tampere University, Tampere, Finland
[28]Institute for Molecular Bioscience, University of Queensland, Brisbane, Australia
[29]Institute of Genetic Epidemiology, Helmholtz Zentrum München - German Research Center for Environmental Health, Neuherberg, Germany
[30]Department of Medicine I, University Hospital Munich, Ludwig Maximilian's University, München, Germany
[31]DZHK (German Centre for Cardiovascular Research), partner site Munich Heart Alliance, Munich, Germany
[32]Institute of Clinical Chemistry and Laboratory Medicine, University Medicine Greifswald, Greifswald, Germany
[33]National Institute for Health and Welfare, University of Helsinki, Helsinki, Finland
[34]Garvan Institute of Medical Research, Garvan-Weizmann Centre for Cellular Genomics, Sydney, Australia
[35]Institute of Human Genetics, Helmholtz Zentrum München, Neuherberg, Germany
[36]Institute of Human Genetics, Technical University Munich, Munich, Germany
[37]Departments of Epidemiology, Medicine, and Health Services, University of Washington, Seattle, United States of America
[38]Kaiser Permanente Washington Health Research Institute, Seattle, WA, United States of America
[39]Centre for Population Health Research, Department of Clinical Physiology and Nuclear Medicine, Turku University Hospital and University of Turku, Turku, Finland
[40]Statistical and Translational Genetics, University of Helsinki, Helsinki, Finland
[41]Department of Internal Medicine, Maastricht University Medical Centre, Maastricht, The Netherlands
[42]Department of Medicine, Universität Leipzig, Leipzig, Germany

[43]Department of Medical Epidemiology and Biostatistics, Karolinska Institutet, Stockholm, Sweden
[44]Center for Molecular and Biomolecular Informatics, Radboud Institute for Molecular Life Sciences, Radboud University Medical Center Nijmegen, Nijmegen, The Netherlands
[45]Institute for Community Medicine, University Medicine Greifswald, Greifswald, Germany
[46]Institute for Laboratory Medicine, LIFE – Leipzig Research Center for Civilization Diseases, Universität Leipzig, Leipzig, Germany
[47]Department of Internal Medicine, Erasmus Medical Centre, Rotterdam, The Netherlands
[48]Department of Neurology, University Medical Center Utrecht, Utrecht, The Netherlands
[49]Interfaculty Institute for Genetics and Functional Genomics, University Medicine Greifswald, Greifswald, Germany
[50]Institute for Advanced Research, Wenzhou Medical University, Wenzhou, Zhejiang 325027, China

**Supplementary Materials and Methods**

*Study cohorts*

Only carriers of recurrent BP4-BP5 CNVs in the 16p11.2 chromosome interval were included as cases in this study. Summary characteristics for each cohort are provided in **Supplementary Table S1**. The Ethics Review Committee on Human Research of the University of Tartu, Estonia and The Committee of Human Research Ethics of the Canton of Vaud, Switzerland approved the project. We obtained written informed consent from biobank or clinical cohort participants, respectively, to perform this study.

*UK Biobank population cohort*

The UK Biobank (UKBB) cohort is a volunteer-based general population biobank of the United Kingdom that aims to provide a platform for studying a range of complex diseases with high relevance to public health. The UKBB follows health data of 500,000 people across the country. The participants aged 40-69 years at the moment of recruitment (2006-2010) were recruited through the National Health Service patient registries. For each participant detailed daily life, socioeconomic, and health-related data were gathered via electronic questionnaires, followed by objective physical evaluation and collection of biological samples. The UKBB governing Research Ethics Committee has approved a generic Research Tissue Bank approval to the UKBB, which covers research using this resource. The Quality Management System of UKBB meets the requirements of the ISO 9001:2015 and ISO27001:2013 standards. A more detailed description of the UKBB is provided in [1] and at http://www.ukbiobank.ac.uk/about-biobank-uk. SNP-genotyping was performed using the Affymetrix Axiom (Thermo Fisher Scientific Inc., Waltham, MA; USA) platform. Data from participants genotyped in the first (n=152,581) and the second release cohort (n=336,195) were subjected to CNV analysis. The two cohorts were analyzed independently.

Health reports from the initial assessment visit (2006-2010) were used to acquire phenotype data. Body mass index (BMI) value at recruitment, recalled age at menarche (AaM) and available characteristics of reproductive health were used. The current study was conducted within UKBB application #17085 "Dissemination of shared genetics across phenotypes associated with reproductive health and related endophenotypes".

*EGCUT population cohort*

The Estonian Genome Center of the University of Tartu (EGCUT) cohort is a general population biobank containing approximately 5% of the Estonian adult population, and is reflective of the age, gender and geographical distribution of the Estonian adult population. At recruitment, an MD-level investigator performed a standardized objective examination of each participant and filled out a questionnaire that encompasses health- and lifestyle-related questions. Additionally, diagnoses of medical conditions present in the medical history of each participant were provided using the WHO international classification of diseases (WHO ICD-10) format. These data are updated continuously through periodic linking to national electronic health registries. EGCUT is conducted according to the Estonian Human Genes Research Act and managed in conformity with the standard ISO 9001:2008. More detailed description of the EGCUT cohort is provided in [2, 3]. SNP-genotyping was performed using the Illumina Infinium platform (Illumina Inc., San Diego, CA; USA). Participant genotype data obtained from the HumanOmniExpress (n=7,509), the Human CNV370 (n=2,146), the Human CoreExome (n=5,068) and the Global Screening Array (n=32,493) BeadChips platforms were subjected to CNV analysis. Phenotype data were obtained from the baseline recruitment questionnaire and from querying the Estonian Health Insurance Fund database (www.haigekassa.ee/en) in July 2016 to collect all entries registered from 2003 through 2015.

*The Pan-European 16p11.2 cohort and The Simons Variation in Individuals (Simons VIP) Collection of 16p11.2 families*

The Pan-European 16p11.2 cohort and The Simons VIP Collection participants were recruited based on direct or cascade diagnosis of 16p11.2 CNV carrier status in probands and their families. Together, they comprise 390 16p11.2 deletion carriers, 270 16p11.2 reciprocal duplication carriers, and their family members; they were recruited and assessed by clinical geneticists in Europe and North America, respectively. Enrollment details and detailed descriptions of these initiatives are provided in [4-6]. For the current study, available reports on pubertal timing and urogenital tract disorders were extracted from the medical history of participants. The latter was analyzed only in the Simons VIP paediatric cohort as information on urogenital and endocrinological diagnoses was not collected systematically for the

European 16p11.2 cohort. Since onset of puberty is known to be strongly heritable, one single CNV carrier per family was randomly selected for the analyses.

*CNV calling, QC and filtering in population cohorts*

We formatted previously produced Log R ratio (LRR) and B Allele Frequency (BAF) values for CNV calling with Hidden Markov Model-based software PennCNV v1.0.4 (from 27.12.2016) [7]. Adjacent CNVs were merged if the length of the interval between them was less than 20% of their combined length. To minimize the number of false positive findings, we excluded the samples with genotyping call-rate <98% or if an individual had more than 200 CNV calls. In addition, we applied the following criteria for sample exclusion: (i) samples with mismatch between their reported and genetic sex; (ii) one of each pair of second-degree or closer relatives. In total, 146,651 individuals of the first release UKBB cohort (discovery cohort), 297,977 individuals of the second release UKBB cohort (replication cohort) and 34,335 EGCUT samples (replication cohort) that passed the post-processing quality control parameters were retained. In this dataset, we extracted all syndromic CNVs (DECIPHER database, https://decipher.sanger.ac.uk/disorders#syndromes) if they had >70% overlap with the corresponding CNV region. We confirmed further all detected 16p11.2 CNV calls by an alternative CNV calling software iPsychCNV (http://www.biopsyk.dk/iPsychCNV; version from 30.05.2017). We included only the CNVs that had an overlap of >70% with the 16p11.2 BP4-BP5 region in the output of each software. The genotype of all EGCUT CNV carriers genotyped with OmniExpress, CNV370 and CoreExome arrays was subsequently confirmed by quantitative PCR as described in [8]. For both UKBB and EGCUT, individuals who had a CNV signal called in the BP4-BP5 region but which remained ambiguous in any of the confirmation steps were excluded from further analysis. Additionally, we excluded all BP4-BP5 CNV carriers (n=9) in the UKBB replication cohort who had second degree or closer relatives in the discovery cohort. We identified 42 deletion and 40 duplication carriers (population prevalence 0.03% for both) in the UKBB discovery cohort; 77 deletion (0.03%; 83 before samples QC and relatives filters) and 70 duplication (0.02%; 76 before samples QC and relatives filters) carriers in the second UKBB cohort; and 12 deletion and 14 duplication carriers (both 0.04%; 12 and 16, respectively, before QC and filtering for relatives) in EGCUT. In the UKBB first release cohort, a subset of 118,327 participants with self-reported "white

British" and genetically confirmed European origin, including 29 deletion and 34 duplication carriers, was used for genotype-phenotype association. In the second cohort, the British European subset was comprised of 246,628 individuals (59 deletion and 64 duplication carriers). All analyzed individuals in the EGCUT cohort were of European origin.

*Genotype-phenotype associations in the 16p11.2 CNV carriers*

All association analyses in population and 16p11.2 clinical cohorts were performed separately in male and female datasets. Women with missing or extreme outlier AaM values (<5 or >25 years) were removed from all phenotype analyses. To model the effect of the 16p11.2 deletion and duplication on the traits of interest, we applied linear regression models adjusted for appropriate covariates. If not otherwise indicated, the R Project for Statistical Computing environment (http://www.r-project.org) was used for statistical analyses and the difference between deletion carriers, duplication carriers and euploid controls was calculated. Depending on the data-type and distribution, Welch two-sample t-test, ANOVA or non-parametric Wilcoxon and Kruskal-Wallis tests were used for comparing the difference between the means of quantitative variables in each carrier status group. Fisher exact test was used to assess the association between binary variables (e.g. diagnosis). Odds ratios (OR), 95% confidence intervals (CI) and P-values were calculated; a threshold of $P \leq 0.05$ was set to indicate statistical significance for a single test. Multiple correction was applied where more than one independent test was used. See **Supplementary Table S2-S5** for the list of analyzed trait categories, phenotype assessment and determination of trait, and disease prevalence in the study cohorts.

*Phenotyping of urogenital organ function in 16p11.2 mouse models*

We used previously reported C57BL/6N mouse models harboring deletion ($16p11.2^{Del/+}$) or duplication ($16p11.2^{Dup/+}$) of the 7qF3 region syntenic to the human 16p11.2 BP4-BP5 rearrangements [9]. All mice were group-housed (3-5 animals per cage) under specific pathogen-free conditions in a room controlled to 21–22°C with a 12-hour light–dark cycle and *ad libitum* access to food and water. One $16p11.2^{Del/+}$ male and one $16p11.2^{Dup/+}$ male were each crossed with two C57BL/6N females (Charles River Laboratories, stock #027) to propagate the colony. To phenotype the female urogenital tract; determine onset of first ovulation; and model estrous cycle, we used 6 female 16p11.2$^{Del/+}$ together with 10 female wild-type (16p11.2$^{+/+}$) littermates, and 9 female 16p11.2$^{Dup/+}$ with 7 female 16p11.2$^{+/+}$ littermates. After weaning at postnatal day 21 (P21), female mice were evaluated visually on a daily basis for vaginal opening. From the day of vaginal opening to P60, vaginal smears were collected daily by inserting a tip of 10ul pipette filled with room temperature PBS into the vagina of the restrained mouse. The PBS was gently flushed two-three times. The final flush from the pipette was transferred to a dry glass and inspected by light microscopy to identify the phase of the estrous cycle, as described [10]. Daily collections of vaginal smears as described above were recapitulated from P95 until P130 to reconstruct the stabilized adult estrous cycle. The body weight of each mouse was measured every second day during both experimental phases. For gross morphological and histological evaluation of urogenital organs, animals were euthanized by $CO_2$ at first diestrus after P130.

To phenotype the male urogenital tract, we euthanized five 16p11.2$^{Del/+}$ and five 16p11.2$^{+/+}$ male littermates, and the same number of 16p11.2$^{Dup/+}$ and 16p11.2$^{+/+}$ male littermates at the age of 14 weeks. Anogenital distance was measured by a caliper positioning the two extremities in the center of the anus to the junction of the smooth perineal skin and in the rugated skin of the scrotum. To assess sperm concentration and motility, sperm were collected from the cauda epididymis of both testes. In short, the epididymis was scratched to allow sperm cell release in pre-warmed M2 media (#M7167, Sigma-Aldrich, St. Louis, MO). Cell suspension was then incubated at 37°C for 10 minutes in a low attachment petri dish to allow sperm swim-up. We diluted the cells to a concentration of <200 million cells/ml; loaded into a counting chamber (depth 100μm; Leja Products, Nieuw-Vennep, The Netherlands); and submitted to computer-assisted sperm analysis (CEROS II; Hamilton Thorne, Beverly, MA, USA). Automated sperm counting was performed at 37°C by recording ≥200 cells with at least four frames acquired at a frame rate of 60 Hz per mouse. We used the following kinematic parameters for sperm motility: smoothed path velocity (VAP; m/s); track velocity (VCL; m/s); straight line velocity (VSL; m/s); and amplitude of lateral head displacement (ALH; m). For histological evaluation, we fixed testes overnight at 4°C in 4% paraformaldehyde, dehydrated the tissue in ethanol series and embedded them in paraffin. Tissue sections (5 μm) were processed for hematoxylin - eosin staining and microscope images were taken in bright field at 10 and 40 times magnification. See **Supplementary Table S12** for the male and female reproductive phenotypes in 16p11.2 mouse model.

We analyzed the resulting data using R environment. Statistical tests to determine significance were selected according to data type: Breslow-Wilcoxon test (age at first estrus); one-way ANOVA (body weight); Student's t-test (all other experiments). Results are expressed as a mean of experiments with standard deviation. Differences were considered statistically significant when $P<0.05$. The number of animals per group was chosen according to the standard practice and requirements in design of animal experiments. The mouse experiments were conducted under protocols reviewed and authorized by the Canton Vaud Food Safety and Veterinary Office (license VD3127.d).

*Structural magnetic resonance imaging (MRI) of 16p11.2 mouse models*

To study mouse brain structure, we performed MRI on 13-week old $16p11.2^{Del/+}$ and $16p11.2^{Dup/+}$ mouse models [11] and control animals as described [12]. We used 10 females $16p11.2^{Del/+}$, 9 females $16p11.2^{Dup/+}$ and 11 females $16p11.2^{+/+}$ littermates maintained on a C57BL/6J background (Jackson Laboratory, Bar Harbor, ME). At 13 weeks of age, animals were anaesthetized with isoflurane and intracardially perfused (rate 1 ml/min) with 30 ml of 0.1 M phosphate-buffered saline (PBS) containing 1000 U/ml of heparin and 2 mM of the gadolinium contrast agent ProHance (Bracco, Milan, Italy) followed by 30 ml of 4% paraformaldehyde (PFA) diluted in PBS containing 1000 U/ml of heparin and 2 mM of ProHance. After perfusion, mice were decapitated and the skin, and lower jaw were removed from the skull containing the brain. Skulls were incubated in 4% PFA diluted in PBS containing 2 mM ProHance overnight at 4 °C, then transferred to PBS containing 2 mM ProHance and 0.02% sodium azide for at least 1 month at 4 °C. Briefly, a multi-channel 7.0 Tesla MRI scanner (Agilent Inc., Palo Alto, CA) was used to image each brain inside the skull. Sixteen custom-built solenoid coils were used to image the brains in parallel [13, 14]. We used the following parameters to detect volumetric changes, w: T2-weighted, 3-D fast spin-echo sequence, with a cylindrical acquisition of k-space, a TR of 350 ms, and TEs of 12 ms per echo for 6 echoes, field-of-view equaled to 20 x 20 x 25 mm3 and matrix size equaled to 504 x 504 x 630. Our parameters output an image with 0.040 mm isotropic voxels. The total imaging time was 14 hours [15]. To visualize and compare any variation within and among genotypes, the images are linearly (6 followed by 12 parameter) and non-linearly registered together. Registrations were performed with a combination of mni_autoreg tools [16] and ANTS (advanced normalization tools)[17, 18]. All scans are then resampled with the appropriate transform and averaged to create a population atlas representing the average anatomy of the study sample. The result of the registration is to have all images deformed into alignment with each other in an unbiased fashion [19, 20]. The Jacobian determinants of the deformation fields are then calculated as measures of volume at each voxel. Significant volume differences can then be calculated by warping a pre-existing classified MRI atlas onto the population atlas, which allows for the volume of 182 different segmented structures encompassing cortical lobes, large white matter structures (i.e. corpus callosum), ventricles, cerebellum, brain stem, and olfactory bulbs to be assessed [21-24]. Further, these measurements can be examined on a voxel-wise basis to localize the differences found within regions or across the brain. Multiple comparisons in this study were controlled for using the False Discovery Rate (FDR) [25].

*Brain structural MRI of human 16p11.2 BP4-BP5 CNV carriers*

For MRI analysis of humans, we selected post-pubertal females from the European and the Simons VIP 16p11.2 consortium with available MRI and AaM data to assess the effect of 16p11.2 dosage on the volume of hypothalamic region. The female-only sample included deletion (n=13, $\mu_{age}$=21.7±12 years, $\mu_{AAM}$=11.1±1 years), duplication carriers (n=22, $\mu_{age}$=36±11.4 years, $\mu_{AAM}$ =13.5±2 years), and intra-familial controls (n=13, $\mu_{age}$=37±9 years, $\mu_{AAM}$=12.6±1.8 years). For the additional analysis with male-only participants we included 16p11.2 deletion carriers (n=31, $\mu_{age}$=21.8±13 years), duplication carriers (n=30, $\mu_{age}$=32.4±14 years) and intra-familial controls (n=27, $\mu_{age}$=30.4±15 years). The local ethics committee approved the protocol. Informed consent was obtained from each participant prior to study inclusion.

Structural MRI data were acquired as described [26-28]. The multi-site data set included both single- and multi-echo images. All multi-echo images were averaged following a Root-Mean Square (RMS) averaging method. For each voxel, RMS calculates the mean of the intensities between the magnitude images of all echo times as follows:

$$RMS_{average} = \sqrt{I(TE1)^2 + I(TE2)^2 + I(TE3)^2 + I(TE4)^2}$$

Image pre-processing was performed using the Statistical Parametric Mapping software SPM12 (www.fil.ion.ucl.ac.uk/spm/software/spm12; Wellcome Trust Centre for Neuroimaging) running under Matlab 7.14 (Mathworks Inc., Sherborn, MA). The automated feature extraction followed the default settings including brain tissue classification in the "unified segmentation" Bayesian framework [29] using an enhanced set of brain tissue priors showing increased accuracy for subcortical structures [30]. For optimal anatomical accuracy the grey and white matter probability maps were spatially registered to a standardized Montreal Neurological Institute space using the diffeomorphic algorithm based on exponentiated lie algebra – DARTEL [31]. The spatially registered grey matter probability maps were scaled with the corresponding Jacobian determinants to preserve the initial total amount of signal intensity followed by spatial smoothing using an isotropic Gaussian kernel of 8 mm full-width-at-half-maximum.

For statistical analysis, a Voxel-Based Morphometry (VBM) ANOVA design was used including the non-CNV carrier participants and four subgroups of 16p11.2 CNV carriers (respectively, deletion and duplication carriers from the European and the Simons VIP cohort). Subject-specific AaM, BMI z-scores, age and its quadratic expansion were integrated as interaction terms; mean grey matter volume (GMV) and total intracranial volume (TIV) represented additional regressors to control for effects of these variables on brain anatomy [32]. In a supplementary analysis we included a male sub-sample consisting of non-carrier participants and four subgroups of 16p11.2 CNV and, besides AaM, use the same demographic and BMI variables as in female sub-sample. Significant results were reported at a statistical threshold of $P<0.05$ after family-wise error correction (FWE) for multiple comparisons and small volume correction as described in [33].

*Disease enrichment analysis of the 16p11.2 dosage-altered genes*

To find 16p11.2 dosage-relevant disease enrichments we exploited previously published transcriptome profiling datasets. These include lymphoblastoid cell lines

(LCL) of human 16p11.2 CNV carriers and brain cortex of the mouse models engineered to carry a deletion or a duplication of the 7qF3 region, respectively [34, 35]. We determined disease term enrichment in the context of altered 16p11.2 gene dosage in the human LCLs (n=1,215 transcripts, GEO Series accession number GSE57802 [34]; n=587 transcripts [35]), and a list of differentially expressed genes in the mouse cortex (n=1,079 [35]) with g:Profiler. We used gene annotations from the Human Phenotype Ontology (HPO); the Online Mendelian Inheritance in Man (OMIM) [36]; and Thomson Reuters *MetaCore*™ analysis suite with Disease Enrichment by Biomarker workflow. The human transcriptome datasets were composed of mostly pre-pubertal probands, hence AaM data were not available for module-trait correlative analyses.

*Mendelian randomization analysis*

We performed univariate and multivariate Mendelian randomization (MR) analysis to test for potential causal effects of 16p11.2 interval genes on regulation of AaM. We used expression quantitative trait loci (eQTLs) detected by the eQTLGen consortium in approximately 14,000 human whole-blood samples to maximize analysis power. To perform summary-level MR with the SMR tool [37], we integrated the eQTL data with SNP-based effect size and standard error estimates from genome-wide association study (GWAS) for AaM sample of 252,000 females [38]. The SMR tool performs a two-sample MR analysis for each gene, using the strongest eQTL of the gene as the only genetic instrument. We used the HEIDI test for heterogeneity [37] to detect whether a causal relationship suggested by SMR could arise due to LD between the instrument and two causal variants, one affecting gene expression variation and the other phenotype trait (e.g. AaM) variation, hampering discrimination between truly causal and pleiotropic models. We considered as potential genetic instruments only eQTLs that were available in GWAS data and had matching alleles. If alleles were switched between datasets (i.e. effect allele in one was alternate allele in other dataset), we flipped the sign of the effect size. SNP genotype data from the UK10K resource (n=3,781; https://www.ebi.ac.uk/ega/datasets/EGAD00001000776) was used to estimate effect allele frequencies in the eQTL and GWAS datasets, and to estimate linkage disequilibrium (LD) between SNPs. We removed from the analysis all SNPs that (i) were not measured in the UK10K data; (ii) had more than 5% missing values; (iii) value of estimated frequency was zero, or (iv) did not have matching alleles. We used a p-value threshold $10^{-5}$ for selecting eQTLs corresponding to F-statistic value of ~20. This selection guarantees inclusion of only strong genetic instruments, but different from more stringent genome-wide thresholds, did not compromise on power (**Supplementary Table S9**). 16 probes with the strongest eQTL passing the threshold were included in the analysis. The probes corresponded to 12 (*C16orf54*, *CORO1A*, *INO80E*, *KCTD13*, *MAPK3*, *MVP*, *PAGR1*, *PPP4C*, *QPRT*, *SPN*, *YPEL3*, *ZG16*) out of 28 unique BP4-BP5 interval genes.

We used equations [7] and [8] from [39] to calculate multivariate causal effect sizes and standard errors:

$$\widehat{\boldsymbol{b}} = \left(\hat{\Gamma}' C^{-1} \hat{\Gamma}\right)^{-1} \hat{\Gamma}' C^{-1} \hat{\gamma}$$

$$\mathrm{Var}(\widehat{\boldsymbol{b}}) = \hat{\sigma}^2 \frac{1}{n} \left(\hat{\Gamma}' C^{-1} \hat{\Gamma}\right)^{-1},$$

Where $\hat{\Gamma}$ is a matrix of standardized eQTL effect sizes, $C$ is an LD-matrix between SNPs, $\hat{\gamma}$ is a vector of standardized GWAS effect sizes, $n$ is GWAS sample size, residual variance $\hat{\sigma}^2 \approx 1$ due to risk factors (gene expression) explaining only a negligible portion of trait variance. If a gene was represented by multiple probes, we chose only one probe for the multivariate analysis. Testing for alternative probe combinations resulted in concordant results (**Supplementary Figure S3**). We used as genetic instruments the eQTLs (n=55) that had available association summary statistics for all genes and were identified by the approximate conditional analysis in GCTA [40] as independent strong eQTLs ($p < 10^{-5}$) for at least one gene.

We used the following equations for the standardized effect sizes and standard errors:

$$\hat{\beta}_{\mathrm{std}} = \frac{Z}{\sqrt{N + Z^2}}$$

$$\mathrm{Var}(\hat{\beta}_{std}) = \frac{1}{\sqrt{N + Z^2}},$$

where $Z$ is the Z-score and $N$ is the sample size of the gene expression or GWAS trait.

*mRNA overexpression and CRISPR/Cas9 genome editing in zebrafish embryos*

To overexpress human wild-type capped mRNA, we linearized pCS2+ constructs and transcribed mRNA using the mMessage mMachine SP6 Transcription Kit (Thermo Fisher Scientific, Waltham, MA) as described [41]. All mRNAs were injected into the yolk of the embryo at the 1- to 2-cell stage at 50, 25 or 12.5 pg doses (1 nl/injection). To deplete *asphd1*, we used CHOPCHOP [42] to identify a guide RNA (gRNA) 5'-GGCATGGGCAGAATTCACAA-3' targeting exon 3. gRNA was transcribed *in vitro* using the GeneArt precision gRNA synthesis kit (Thermo Fisher Scientific) according to the manufacturer's instructions; 1 nl of injection cocktail containing 100 pg/nl gRNA and 200 pg/nl Cas9 protein (PNA Bio, Thousand Oaks, CA) was injected into the cell of embryos at the 1-cell stage. To determine targeting efficiency in founder (F0) mutants, we extracted genomic DNA from 2 day post-fertilization (dpf) embryos and PCR amplified the region flanking the gRNA target site using primers exon3F: 5'-TCTCTTCTCCTAAATCATCGGC-3' and exon3R: 5'-TCCTGGCCTAAAGAAACAAAGA-3'. PCR products were denatured, reannealed slowly and separated on a 20% TBE 1.0-mm precast polyacrylamide gel (Thermo Fisher Scientific), which was then incubated in ethidium bromide and imaged on a ChemiDoc system (Bio-Rad, Hercules, CA) to visualize hetero- and homoduplexes. To estimate the percentage of mosaicism of *asphd1* F0 mutants (n=5/condition), PCR products were gel purified (Qiagen, Germantown, MD), and cloned into the pCR8/GW/TOPO-TA vector (Thermo Fisher Scientific). Plasmid was prepped from individual colonies (n=10–12 colonies/embryo) and Sanger sequenced according to standard procedures.

*Automated zebrafish imaging*

We obtained zebrafish embryos from natural matings of heterozygous Tg(*gnrh3*:*egfp*) transgenic adults [43] and maintained the larvae under standard conditions at 28.5°C until 5 dpf. Automatic imaging was conducted with an AxioScope.A1 microscope and Axiocam 503 monochromatic camera facilitated by Zen Pro software (Zeiss, Oberkochen, Germany), to capture dorsal images of GFP signal. Larval batches were positioned and imaged live using the Vertebrate Automated Screening Technology (VAST; software version 1.2.5.4; Union Biometrica) BioImager. We anesthetized larvae from each experimental condition with 0.2 mg/ml Tricaine prior to being loaded into the sample reservoir. We acquired dorsal and lateral image templates of uninjected controls and experimental larvae at a >70% minimum similarity for the patternrecognition algorithms. The larvae were rotated to 0° to acquire a dorsal image via a 10x Fluar objective (Zeiss) and fluorescent excitation at 470 nm to detect GFP. To perform automated measurement of VAST images, we saved each file corresponding to an experimental condition in CZI file format in Zen Pro. Images were batch processed by converting each CZI to individual TIFF files, which could be opened in ImageJ (NIH, Bethesda, MD). All images were stacked and converted to 8-bit format; threshold was adjusted to 60 (minimum) and 160 (maximum). Images were then measured quantitatively by the highlighted area of the fluorescent signal in each image with the ImageJ macro. All experimental conditions were normalized to uninjected controls and set to an arbitrary unit of 100. Statistical comparisons were performed using one-way ANOVA with Tukey's test (GraphPad Prism, San Diego, CA).

*ASPHD1 global gene expression similarity analyses*

Global gene expression similarity was analyzed using Multi Experiment Matrix (MEM) webtool [44]. MEM collects thousands of gene expression experiments from ArrayExpress repository (https://www.ebi.ac.uk/arrayexpress) and provides a robust rank aggregation method [45] to find Affymetrix probe pairs that compared to the query probe have the most similar expression patterns across hundreds of experiments. We queried for genes with expression profiles globally similar to that of *ASPHD1/Asphd1* using the probe set "1553997_a_at" for human and "1456837_at" for mouse. We used the largest Affymetrix microarray platforms available: HG U133 2.0 (>2,800 individual datasets) for human and Mouse 450 2.0 (>2,400 datasets) for mouse analyses.

Enrichment analysis was conducted using g:Profiler toolset [46] with default settings and an ordered query option. This performs incremental enrichment analysis with increasingly larger numbers of genes starting from the top of the list, and reports the minimal p-value that might lead for each of the enriched term's individual length of query genes taken into account. The latter feature is preferred in the analyses of ordered gene lists (e.g. differential expression analysis, gene lists ordered by global expression similarity). We used the default output of MEM tool with the 150 most similar Affymetrix probe sets to those of the *ASPHD1* as a g:Profiler query for both mouse and human. The mouse gene identifiers were converted to human genes for effortless comparison of the results from both species.

To assess similarity across human tissues, we used GTEx expression data from 38 tissues (https://gtexportal.org/home). The funcExplorer webtool [47] defines optimal clusters and re-groups genes to these clusters based on their tissue-specific expression profiles, potential gene regulators, Gene Ontology and pathway terms provided by the g:Profiler. We used default parameters with Pearson correlation for distance measure, and the "best annotation strategy" for cluster definition to determine the *ASPHD1* cluster membership.

#### Supplementary Results

*Association between BMI and 16p11.2 BP4-BP5 dosage in unselected population cohorts*

We first assessed whether unselected adult carriers of the 16p11.2 CNVs present the mirror effect on BMI reported previously in clinical cohorts [48]. In the first UKBB cohort, BMI values were associated significantly with the genomic dosage of the 16p11.2 interval (ANOVA $p=5.33e^{-25}$, all ethnic groups; ANOVA $p=2.48e^{-17}$, Europeans only). In the “combined ethnicities” set, compared to controls (males: $\mu_{BMI}=27.9$; females: $\mu_{BMI}=27.2$) both male (n=22, $\mu_{BMI}=34.7$, $\Delta=+6.8$; $p=2.48e^{-13}$, ANOVA) and female (n=20, $\mu_{BMI}=35.7$, $\Delta=+8.5$; $p=1.57e^{-13}$, ANOVA) deletion carriers had significantly higher BMI. We found that this signal was the strongest BMI association across all DECIPHER-listed recurrent CNVs that were present in the UKBB population sample (**Supplementary Figure S2A**). Although less pronounced, the BMI difference was opposite for duplication carriers (males: n=14, $\mu_{BMI}=26.7$, $\Delta=-1.2$, p=0.27; females: n=26, $\mu_{BMI}=25.3$, $\Delta=-1.9$, ANOVA p=0.049). We replicated the mirror effect on BMI in the second UKBB ($p=3.8e^{-44}$, all ethnic groups combined; controls: n=297,830, $\mu_{BMI}=27.4$; deletions: n=77, $\mu_{BMI}=34.9$; $\Delta=+7.5$; duplications: n=70, $\mu_{BMI}=25.9$, $\Delta=-1.5$) and EGCUT cohorts ($p=6.4e^{-08}$; controls: n=34,309, $\mu_{BMI}=26.3$; deletions: n=12, $\mu_{BMI}=32.9$, $\Delta=+6.6$; duplication: n=14, $\mu_{BMI}=22.8$, $\Delta=-3.5$). Detailed results by sex and ethnicity are listed in **Supplementary Figure S1A-C and Table S2**.

*Association between age at menarche and 16p11.2 BP4-BP5 dosage*

We next evaluated associations between AaM, the first menstrual cycle and a proxy for the onset of female puberty, and DECIPHER-listed CNVs in the first UKBB cohort. Seven recurrent reciprocal CNVs are sufficiently prevalent (overlap with recurrent CNV interval >70%, maximum length 1.2x the recurrent interval, ≥5 carriers for each copy-number state) in the UKBB dataset to allow formal testing of dosage-effect. Only the 16p11.2 BP4-BP5 CNVs showed significant dosage-dependent association with AaM after correction for the number of effective tests (ANOVA $p=1.93e^{-05}$, corrected for BMI, year of birth and four PCs; European females only; **Figure 1** and **Supplementary Figure 2B**). This trait has been associated previously with common SNPs in the proximal part of the BP4-BP5 interval [38, 49, 50], but, to our knowledge, not linked to a rare variant in the region such as the 16p11.2 CNV. Compared to controls (n=60,466; $\mu_{AAM}$=12.9 years, European females only) AaM was significantly decreased in deletion carriers (n=11; $\mu_{AAM}$=11.4 years, Δ=-1.5 years; p=0.00098, Wilcoxon) and, on the contrary, increased in duplication carriers (n=21; $\mu_{AAM}$=14.4 years, Δ=+1.5 years; p=0.002, Wilcoxon). Our results indicate that while the 16p11.2 mirror association with BMI in unselected populations was to a greater extent driven by the deletion, the duplication is largely responsible for the dosage-dependent AaM association (**Supplementary Table S3** and **Figure S1G-I**).

We replicated the association between 16p11.2 dosage and AaM in the second UKBB cohort ($p=6.5e^{-04}$, ANOVA, corrected for BMI, year of birth and PC 1-4, European females only). Compared to controls (n=129,943, $\mu_{AAM}$=12.95 years), deletion carriers had a significant decrease (n=21, $\mu_{AAM}$=11.86 years, Δ=-1.09 years; p=0.0016, Wilcoxon) and the duplication carriers a significant increase (n=27, $\mu_{AAM}$=14.0 years, Δ=+1.05 years, p = 0.0063, Wilcoxon) in AaM. The results obtained in the geographically distinct EGCUT cohort were consistent with those of UKBB ($p=2.4e^{-05}$, ANOVA, corrected for BMI, year of birth and PC 1-4). Compared to controls ($\mu_{AAM}$=13.5 years), the mean AaM was lower in deletion carriers (n=8; $\mu_{AoM}$=12.3 years, Δ=-1.2 years, Wilcoxon p=0.017, Wilcoxon) and higher in duplication carriers (n=10; $\mu_{AAM}$=15.6 years, Δ=+2.1 years; p=0.0014, Wilcoxon). Additionally, the set of UKBB female participants of non-European origin showed the same trend (controls: n=41,640, $\mu_{AAM}$=13.03 years; deletions: n=14, $\mu_{AAM}$=11.86 years, Δ=-1.17 years, p=0.0056, Wilcoxon; duplications: n=3, $\mu_{AAM}$=14.67

years, Δ=+1.64 years, p=0.093, Wilcoxon) despite the low numbers of CNV carriers with reported AaM (**Supplementary Table S3** and **Figure S1D**).

We confirmed further the dosage-dependent impact on AaM by comparing unrelated post-pubertal female deletion and duplication carriers in two independent 16p11.2 clinical cohorts. The mirror effect was significant in both the Simons VIP Collection (deletions: n=22, $\mu_{AAM}$=11.0 years; duplications: n=20, $\mu_{AAM}$=13.3 years; Δ=+2.3 years, p=7.7e$^{-05}$) and in the Pan-European cohort (deletions: n=14, $\mu_{AAM}$=11.6 years; duplications: n=11, $\mu_{AAM}$=13.5 years; Δ=+1.9 years, p=0.0034) (**Supplementary Figure S1EF**). Consistent with general population cohorts, ethnicity had no impact on the association signal and compared to CNV carrier status, the effect of BMI z-score on the pubertal timing was marginal (Simons VIP: beta=-0.09 years, p=0.77; Pan-European: beta=-0.001 years, p=1).

*16p11.2 BP4-BP5 dosage and male pubertal timing*

Pubertal onset in males is not as distinctively defined as in females, nor is there reliable quantitative data available for population biobank participants. We used indicative data collected for male participants in the UKBB cohort and in the Simons VIP 16p11.2 collection to evaluate whether the altered timing of puberty of BP4-BP5 CNV carriers is sex-specific. Although we found no statistically significant difference in the first UKBB cohort (all ethnicities combined), we observed, nonetheless, a consistent trend. Compared to controls (n=66,300; had facial hair at "younger than average age" 6.9%, "about average age" 80.3%, "older than average age" 12.8%) deletion carriers trended toward having first facial hair at "younger than average age" (n=21; "younger" 19.0%, "about average" 81.0%, "older" 0%; OR=2.73, p=0.08), and duplication carriers at "older than average age" (n=14; "younger" 7.1%, "about average" 64.3%, "older" 28.6%; OR=2.78, p=0.09). These trends replicated in the second UKBB cohort (controls: n=129,904; "younger" 6.75%, "about average" 80.51%, "older" 12.74%; deletions: n=38; "younger" 18.42%, "about average" 81.58%, "older" 0%, OR=2.69, p=0.025; duplications: n=36; "younger" 2.78%, "about average" 77.78%, "older" 19.44%, OR=1.58, p=0.32). We obtained similar results for the relative age at voice break (**Supplementary Table S4**).

Due to an unequal distribution of post-pubertal deletion and duplication carriers in pediatric (<18 years; deletion: n=21, duplication: n=6) and adult (>18 years; deletion: n=6, duplication: n=18) datasets of the Simons VIP collection, we analyzed these sub-cohorts separately. All five puberty-related traits (age at start of "growth spurt", "facial hair growth", "body hair growth", "skin changes" and "voice break") showed directionally consistent trends of advanced pubertal age in duplication carriers compared to deletions in both datasets (Δ ranging from +1.2 years to +2.5 years; all p-values <0.2; **Supplementary Table S4**). Four traits reached nominal significance levels at least in one sub-cohort (adult male cohort: "growth spurt" $\mu_{DEL}$=11.7 years, $\mu_{DUP}$=14.2 years, Δ=+2.5 years, p=0.01; "body hair growth" $\mu_{DEL}$=11.5 years, $\mu_{DUP}$=13.8 years, Δ=+2.3 years, p=0.04; pediatric male cohort: "facial hair growth" $\mu_{DEL}$=12.4 years, $\mu_{DUP}$=14.0 years, Δ=+1.6 years, p=0.04; "voice break" ($\mu_{DEL}$=11.9 years, $\mu_{DUP}$=13.5 years, Δ=+1.6 years, p=0.05). These results suggest that pubertal onset is affected by alterations of the 16p11.2 genomic dosage in both males and females.

*Disorders of reproductive health and sexual development in 16p11.2 CNV carriers*

We then assessed if perturbation of sexual maturity was correlated with impaired reproductive health. We tested the impact of 16p11.2 CNVs on sexual development by analyzing diagnoses related to disorders of reproductive tract and fertility. In the first UKBB release cohort, we found that compared to controls, female deletion carriers had a significant decrease in the number of live births (controls μ=1.81, deletions μ=0.91; P=0.013, Wilcoxon rank-sum test, Europeans only). We replicated this signal also in the second UKBB release (controls μ=1.81 deletions μ=0.78, P=2.9$e^{-05}$, Wilcoxon rank-sum test, Europeans only; **Supplementary Table S5**). However, this trait could be influenced by other features associated with the 16p11.2 deletion (e.g. cognitive, social and behavioral problems, congenital malformations). We also found that compared to controls, female duplication carriers in the UKBB first release had increased risk for miscarriages (controls: 20.5%; duplications: 39%, 9 out of 23; OR=2.5, P=0.037, Fisher's exact test, Europeans only). This did not replicate in the second UKBB release (controls: 20.5%; duplications: 29%, 8 out of 28; OR=1.55, P=0.35, Fisher's exact test, Europeans only; **Supplementary Table S5**). Maternal age at first and last live birth was not significantly different from general population in either of the CNV groups.

The associations between 16p11.2 CNVs and reproductive health was further confirmed by increased diagnoses of the female reproductive tract disorders in the EGCUT cohort, such as "absent, scanty and rare menstruation" (N91 according to the ICD-10 classification; OR=4.4, P=0.013, Fisher's Exact Test), "noninflammatory disorders of ovary, fallopian tube and broad ligament" (N83; OR=5.2, p=0.003, Fisher's Exact Test) and "inflammatory diseases of female pelvic organs" (N70-N77; OR=4.1, P=0.012, Fisher's Exact Test). Our data shows that altogether 50% and 64% of deletion and duplication EGCUT females were diagnosed with irregular/absent menstruation, hormonal disorders or (non-inflammatory) ovarian dysfunction (**Figure 1F** and **Supplementary Table S5**).

With respective live birth rates of 1.6-9.0% (reviewed in [51] and 0.2-0.4% [52] in Europe and Northern America, hypospadias and cryptorchidism are among the most common urogenital defects in boys. We observed that pediatric male duplication carriers in the Simons VIP cohort had an increase in undescended testicles (13.2%, n=5) and hypospadias (5.3%, n=2). By including diagnoses of other genital problems, 11 out of 38 (29.0%) boys with the 16p11.2 duplication were affected. Data about inborn genital defects was not available for adult male carriers in the population, nor in the clinical cohorts.

*Transcripts affected by the 16p11.2 dosage alterations are enriched for urogenital disease genes*

Next, we wondered if reproductive traits in 16p11.2 CNV carriers were paralleled by transcriptomic alterations. We found that the 1,188 16p11.2 dosage-dependent differentially expressed genes (DEGs) in lymphoblastoid cell lines (LCLs) [34] were highly enriched for genes associated with urogenital diseases (12 of the top-25 non-neoplastic terms; all with FDR <2.6e$^{-03}$). We replicated this result using LCLs of 16p11.2 CNV carriers from multiplex ASD families [35] (9 of the top-25 non-neoplastic terms; all with FDR <4.9e$^{-04}$). Similarly, urogenital dysfunction was the top dosage-sensitive disease enrichment when we profiled the cerebral cortex of mice carrying the syntenic 16p11.2 deletion or duplication [35] (11 of the top-25 non-neoplastic terms; all with FDR <4.9e$^{-09}$) (**Supplementary Table S6**).

*16p11.2 dosage and sexual development in mice*

The similarities observed between the genes differentially expressed in human 16p11.2 LCLs and mouse cortices motivated us to assess if 16p11.2 dosage was affecting sexual development in murine models. First, we determined age of sexual maturation in female mice carrying either the syntenic deletion or duplication and their respective controls. Consistent with previous reports the 16p11.2$^{Del/+}$ and 16p11.2$^{Dup/+}$ animals showed a mirror effect on body weight that is opposite to that observed in human 16p11.2 CNV carriers [9, 53]. The 16p11.2$^{Del/+}$ mice exhibited reduced body weight (female animals, at postnatal day 21 (P21): $\mu_{WT}$=10.45 g, $SD_{WT}$=1.15, $\mu_{DEL}$=8.2 g, $SD_{DEL}$=1.0, p=0.001; at first estrous: $\mu_{WT}$=15.79 g, $SD_{WT}$=1.38, $\mu_{DEL}$=14.28 g, $SD_{DEL}$=1.03, p=0.048; at age P97: $\mu_{WT}$=23.98 g, $SD_{WT}$=1.7, $\mu_{DEL}$=18.13 g, $SD_{DEL}$=1.13, p=6.3e$^{-06}$, Student's t-test) and the 16p11.2$^{Dup/+}$ mice showed a trend for increased body weight (female animals, at age P21: $\mu_{WT}$=9.26 g, $SD_{WT}$=1.26, $\mu_{DUP}$=9.01 g, $SD_{DUP}$=1.08, p=0.35; at first estrous: $\mu_{WT}$=16.3 g, $SD_{WT}$=0.94, $\mu_{DUP}$=16.52 g, $SD_{DUP}$=1.15, p=0.7; at age P97: $\mu_{WT}$=22.76 g, $SD_{WT}$=2.04, $\mu_{DUP}$=24.03 g, $SD_{DUP}$=2.86, p=0.83, Student's t-test; **Supplementary Table S12**). In line with previous publications [54, 55], we used the cumulative percentage of animals reaching their first ovulation (i.e. estrus phase of the first cycle) as a function of age to measure pubertal maturation in female mice. Although the influence of body weight cannot be completely excluded, we observed a mirror effect on sexual maturation, which similar to weight, was swapped when compared to humans. While 50% of control animals in our 16p11.2 deletion cohort exhibited their first estrus at P32, 50% of their 16p11.2$^{Del/+}$ littermates reached first ovulation five days later at P37 (p=4.8e$^{-06}$, Breslow-Wilcoxon test). Conversely, compared to their matched controls (50% reaching first estrus at P36), 50% of the 16p11.2$^{Dup/+}$ mice showed initial ovulation advanced by two days (P34; p=0.0021, Breslow-Wilcoxon test) (**Figure 2AB** and **Supplementary Table 12**). Following sexual maturation we assessed estrous cyclicity until P59 that corresponds to the end of adolescence and repeated the experiments at the age that corresponds to adult age with expected stable cyclicity (from P95 through P127). Whereas daily inspections of vaginal cytology showed normal estrous cyclicity, 16p11.2$^{Del/+}$ females showed perturbation of time spent in the estrus (wild type: $\mu_{days}$=12.4, SD=2.37; 16p11.2$^{Del/+}$: $\mu_{days}$=16.16, SD=2.85; p=0.034; Student's t-test) and diestrus (wild type: $\mu_{days}$=8.6, SD=2.24; 16p11.2$^{Del/+}$: $\mu_{days}$=6.33, SD=1.25; p=0.029; Student's t-test) phases of their cycle. The time spent in each phase and total number of cycles did not differ significantly in 16p11.2$^{Dup/+}$ mice compared to controls. (**Supplementary Table S12**).

Motivated by our human data on reproductive health, we assessed both female and male 16p11.2 mouse models to determine if altered sexual development was accompanied by gross morphological changes of the reproductive organs. We found that 16p11.2$^{Dup/+}$ females had significantly increased uterine size compared to estrous cycle harmonized littermate controls (controls: n=3, μ=36.0 mg; 16p11.2$^{Dup/+}$: n=3, μ=55.0 mg; p=0.044, Student's t-test, normalized for body weight). The size of ovaries remained unchanged in both 16p11.2$^{Dup/+}$ (littermate controls: n=3, μ=7.17 mg,: 16p11.2$^{Dup/+}$: n=3, μ=8.73 mg, p=0.61, Student's t-test, normalized for body weight) and 16p11.2$^{Del/+}$ (littermate controls: n=3, μ=10.7 mg, 16p11.2$^{Del/+}$: n=3, μ=9.1 mg, p=0.51, Student's t-test, normalized for body weight). See **Supplementary Table S12** for full results.
In males, the weight of urogenital tract organs, such as testes and kidney did not differ significantly after normalization for body weight in either of the mutant groups. However, we observed a significant reduction of ano-genital distance (AGD) in the 16p11.2$^{Dup/+}$ animals (n=5, $\mu_{AGD}$=1.52 cm) compared to controls (n=5, $\mu_{AGD}$=1.84 cm; p=0.004, Student's t-test, normalized for body weight). The AGD varies as a function of prenatal exposure to circulating androgen and is a commonly used endpoint for hormonally regulated disorders of sexual differentiation in rodents [56]. Normally, this distance is markedly longer in males than in females. Its reduction in males is indicative of abnormal reproductive tract masculinization, sexual dimorphism and has been associated with a higher risk for cryptorchidism and hypospadias. We did not observe any significant variation of AGD after normalization for body weight in 16p11.2$^{Del/+}$ males (n=5, $\mu_{AGD}$=1.54 cm) versus controls (n=5, $\mu_{AGD}$=1.73 cm, p=0.24, Student's t-test, normalized for body weight). See **Supplementary Table S12** for full results.

We further evaluated the histological architecture of testicular tubules and observed that two of five 16p11.2$^{Dup/+}$ mice showed tubular regions with abnormal accumulation of germ cells in the lumen and a third animal presenting with abnormally shaped vacuolar tubules (**Figure 2E**). Finally, we evaluated spermatogenesis and fertilizing capacity by investigating both sperm concentration (normalized to the testicular weight) and sperm motility. The sperm concentration of animals carrying either of the

16p11.2 CNVs was not statistically different from that of respective controls. These results indicate that the cellular phenotype probably does not impair the testicular structure or function to the point of infertility. However reduction of AGD suggests reduction of the levels of circulating androgens in the male 16p11.2$^{Dup/+}$ mice.

*Association between 16p11.2 and hypothalamic volume in humans and mice*

We hypothesized an involvement of the hypothalamus in the observed 16p11.2 CNV-dependent reproductive phenotypes, and used human MRI data from the Simons VIP and the European 16p11.2 clinical cohorts [26, 28] to assess possible gene dosage-dependent structural changes in the brain. We restricted the search volume to a 5 mm sphere centered on the hypothalamic region. We found a significant negative interaction ($p_{FWE}<0.05$) between the 16p11.2 CNV dosage and the volume of the hypothalamic region in female, but not in male, CNV carriers. Female deletion and duplication carriers have larger and smaller hypothalamic volume than controls, respectively. Whereas deletion carriers showed positive correlation between the hypothalamus volume and AaM, duplication carriers exhibited negative correlation (**Figure 2F and Supplementary Table S7**).

Correspondingly, abnormal hypothalamic morphology has been reported previously in 16p11.2$^{Del/+}$ mouse models [11, 57]. We observed alterations of the hypothalamic volume in both 16p11.2$^{Del/+}$ and 16p11.2$^{Dup/+}$ female mice (**Supplementary Table S8**). At 13 weeks both strains presented a decrease in the absolute volume of the hypothalamus compared to wild type (wt) littermates (wt, n=12, 11.14±0.27mm$^3$; 16p11.2$^{Del/+}$, n=10, 10.68±0.41mm$^3$; 16p11.2$^{Dup/+}$, n=9, 10.52±0.31mm$^3$), which resulted in a negative correlation between 16p11.2 dosage and hypothalamus volume relative to total brain volume. Compared to wt littermates (2.396±0.030), the 16p11.2$^{Del/+}$ and 16p11.2$^{Dup/+}$ mice have a respectively larger (2.505±0.031, p<0.0001, one-way ANOVA, post-hoc Tukey) and smaller (2.301±0.035, p<0.0001, one-way ANOVA, post-hoc Tukey) hypothalamus relative volume **Supplementary Table S8** and **Figure 2G**). Our results suggest that perturbation of neurons secreting GnRH, a hormone that is essential for initiating puberty and activating the hypothalamus–pituitary–gonadal axis, could contribute to the observed alterations of sexual development.

*Mendelian Randomization to identify genes with causal effect on age at menarche*

To identify which of the 28 genes in the 16p11.2 interval are causally related with pubertal timing, we first performed univariate SMR analysis using the most recent GWAS summary statistics for AaM [38] and an eQTL meta-analysis of 14,115 whole-blood samples [58]. We were able to test for a causal effect of 12 genes with strong genetic instruments, more than double the number of genes analyzed previously in the 16p11.2 region [38]. We identified three genes that showed evidence of causality ($P_{SMR} < 0.05/12$) while also passing the HEIDI test for heterogeneity ($P_{HEIDI} > 0.009$): *INO80E* (standardized causal effect size b=0.098, SE=0.015, p=1.3e$^{-10}$), *KCTD13* (b=-0.15, SE=0.034, p=4.5e$^{-6}$) and *MAPK3* (b=0.032, SE=0.0053, p=9.8e$^{-10}$) (**Figure 3B; Supplementary Table S9)**. As high gene density and physical overlap of genes in the 16p11.2 interval makes it difficult to identify valid genetic instruments for MR analysis and to account for pleiotropy, we performed in parallel a multivariate MR analysis using all 55 independent eQTLs of 16p11.2 genes as instruments. Consistent with the results of univariate SMR, we confirmed a causal effect of *INO80E* (b=0.071, SE=0.018, p=9.3e$^{-5}$) and *KCTD13* (b =-0.074, SE=0.023, p=9.7e$^{-4}$) (**Figure 3C; Supplementary Table S9**). *MAPK3* lost its significance in the multivariate analysis (p=0.98) possibly because the strongest genetic *MAPK3* variant was a strong eQTL for multiple other genes, including *INO80E* and *KCTD13*. However, the multivariate MR model measures the direct causal effect of each gene on AaM [59]. *MAPK3* might, alternatively, have an indirect effect mediated by *INO80E* or *KCTD13*. *YPEL3* was identified as having a positive effect on AaM in the multivariate MR analysis (b=0.071, SE=0.025, p=3.8e$^{-3}$), while not passing the HEIDI test for heterogeneity in the SMR analysis. The close proximity to and shared eQTLs of *YPEL3* with *KCTD13*, *INO80E* and *MAPK3* do not allow for conclusions about a possible causal role in AaM (**Supplementary Table S9**).

*ASPHD1 dosage influences GnRH neuron patterning*

One limitation of our MR analyses is that less than half (12 out of 28) of the 16p11.2 interval genes could be included in the MR analyses. Further, we recognize that whole blood might not be the most relevant tissue to reproductive phenotypes. Therefore, we modulated the dosage of all 16p11.2 genes in a *Tg(gnrh3:egfp)* transgenic zebrafish model [43, 60]. GnRH3 is expressed by neurons in the forebrain areas of olfactory bulb-terminal nerve, preoptic area, and hypothalamus [61]. We first evaluated the phenotypic consequences of human mRNA overexpression to model the 16p11.2 duplication condition. We injected each mRNA individually at a dosage of 50 pg, except *KIF22* and *PPP4C* which induced non-specific toxicity phenotypes and/or lethality; these were injected at a non-lethal dose of 12.5 pg, per embryo (n=30-50 larvae/batch, minimum three batch replicates). We performed automated live imaging of the dorsal aspect of larvae that were allowed to develop to 5 days post fertilization (dpf), measured the area of GFP-positive cells for each experimental condition, and compared relative GnRH3-GFP reporter neuron area across batches. We found that overexpression of a single transcript, *ASPHD1*, was sufficient to significantly reduce GFP signal in comparison with controls (19% of reduction; $p<0.0001$) (**Figure 4A-C** and **Supplementary Table S10**).

To model *ASPHD1* hemizygosity (i.e. deletion model), we used CRISPR/Cas9 genome editing to deplete endogenous *asphd1* transcripts (58% identity; 69% similarity for human ASPHD1 protein [GenBank ID: NP_859069.2] versus zebrafish [UniProt ID: E7F0P5]. We first defined guide RNA (gRNA) target sites in the *asphd1* locus, transcribed gRNAs *in vitro* and coinjected 100 pg with 200 pg of Cas9 protein into one-cell stage *gnrh3:egfp* embryos. Heteroduplex analysis followed by sequencing of cloned PCR fragments flanking each gRNA target site demonstrated induction of 96% mosaicism (n=5 embryos; n=10-12 colonies/embryo) (**Supplementary Figure S4**). We assessed the GnRH3:GFP neuron area of F0 (founder) larval batches at 5 dpf and observed a significant decrease of signal in *asphd1* F0 mutant larvae compared to controls (13%, p=0.0031). Notably, injection of 100 pg of the *asphd1* gRNA in absence of Cas9 produced no detectable changes in GnRH3 neuronal patterning (**Figure 4D** and **Supplementary Table S10**).

We have previously shown that genes within the 16p11.2 cytoband can interact to induce additive or epistatic phenotypes related to the neuroanatomical phenotype [41, 62]. To investigate whether the *ASPHD1* effect on reproductive axis could be exacerbated in the context of other 16p11.2 genes, we co-expressed it with other genes that showed causal evidence in MR analysis. To test for epistasis, we halved the dose of

*ASPHD1* to 25 pg, co-injected with 50 pg of *KCTD13*, *INO80E*, *MAPK3* or *YPEL3* and quantified the GnRH3:GFP area at 5 dpf. We found that the larvae co-injected with *ASPHD1* and *KCTD13* showed a significantly aggravated phenotype (compared to controls: 24% reduction, p<0.0001; compared to *ASPHD1* mono-injected batches: 14% reduction, p=0.003) suggesting genetic interaction between these two genes (**Figure 4E** and **Supplementary Table S11**). We observed no genetic interaction between *ASPHD1* and either *MAPK3* or *YPEL3* and only a modest interaction between *ASPHD1* and *INO80E*, wherein the GnRH3:GFP area was reduced compared to controls (12%, p=0.018), but remained unchanged compared to the batches injected with either mRNA alone (**Supplementary Table S11**).

*ASPHD1 expression profile is globally similar to genes involved in neuron projection and synaptic vesicle function*

As minimal information is available about the functionality of *ASPHD1*, we used global gene expression similarity analysis by MEM across hundreds of human and mouse gene expression data sets to understand its potential role in biological processes. We found that in human data sets, *ASPHD1* has globally the most similar expression patterns with the *SEZ6L2* in close physical proximity in the 16p11.2 (**Supplementary Figure S5A**; the results are accessible at **http://bit.ly/asphd1hsapiens**). In the mouse data, *SEZ6L2* is the second gene by similarity to *ASPHD1* after *CELF4*, a gene mapping to the long arm of human chromosome 18 (**Supplementary Figure S5B**; the results are accessible at **http://bit.ly/asphd1mmusculus**). *CELF4* is involved in neurodevelopmental regulation [63]. It was associated through GWAS with educational attainment [64], neuroticism and mood related alterations [65-67]. By selecting 50 genes with the most similar global expression to *ASPHD1* in human and mouse data for enrichment analysis, we found enrichment for GO terms related to synaptic vesicle function and neurotransmission in both mice and humans (**Supplementary Figure S6**). The detailed results are available at **https://biit.cs.ut.ee/gplink/l/RfK8rsZHRf**.

For similarity analysis across human tissues we used GTEx expression data from 38 tissues. *ASPHD1*, *SEZ6L2* and *CELF4* share expression patterns restricted to the brain and pituitary gland. *ASPHD1* and *CELF4* belong to the same cluster of 258 genes significantly annotated by three neuronal processes "neuron projection" (GO:0043005, p=2.16e$^{-09}$), "transmission across chemical synapses" (REAC:112315, p=1.04e$^{-10}$), "neurotransmitter receptor binding and downstream transmission in the postsynaptic cell" (REAC:112314, p=3.7e$^{-09}$) (**Figure 3D**; the results are accessible at **https://biit.cs.ut.ee/gplink/l/AOYgmspNQu**).

**Supplementary Figures and Tables**

Supplementary Figures and Tables appear at the end of this document.

**Supplementary Figures Legends**

**Supplementary Figure S1. Mirror associations of the 16p11.2 CNVs with BMI and AaM in adult population and clinical cohorts.** Association with BMI in (**A**) the UKBB first release, (**B**) the UKBB second release, and (**C**) the EGCUT cohort. Only individuals of European descent were included in the analysis.

Association with AaM in (**D**) the UKBB non-European sub-cohort, (**E**) the European 16p11.2 clinical cohort, and (**F**) the Simons VIP 16p11.2 clinical cohort. Variance in pubertal onset in female 16p11.2 CNV carriers in (**G**) the UKBB first release cohort, (**H**) the second release cohort and (**I**) the EGCUT cohort. Only European individuals are shown; yellow represents the 16p11.2 deletion carriers, blue duplication carriers, and grey non-CNV carrier controls.

**Supplementary Figure S2. Dosage-dependent associations between DECIPHER-listed recurrent CNVs, BMI and AaM.** Associations of DECIPHER-listed CNVs with (**A**) BMI and (**B**) AaM were evaluated in the first release UKBB cohort. Seven recurrent reciprocal CNVs were sufficiently prevalent (overlap with recurrent CNV interval >70%, maximum length 1.2x the recurrent interval, ≥5 carriers for each copy-number state) in the UKBB dataset to allow formal testing of dosage-effect. Only the 16p11.2 BP4-BP5 CNVs showed significant dosage-dependent association with BMI (corrected for year of birth, PC1-4, the number of effective tests) and AaM (corrected for BMI, year of birth, PC1-4 and the number of effective tests). Black dots represent -log10 adjusted p-value in each evaluated CNV loci; horizontal line represents the significance level =

0.05/(tests for BMI + tests for age of menarche) = 0.5/(7+7), i.e. number of effective tests.

**Supplementary Figure S3. Multivariate Mendelian randomization analysis using every possible combination of probes demonstrate similar results.** Standardized causal effect estimates for AaM with 95% confidence intervals (**A-L**). Genes with multiple probes are *MVP* (ILMN_1803277, ILMN_2344373), *MAPK3* (ILMN_2402341, ILMN_1667260, ILMN_1812747) and *C16orf54* (ILMN_1751061, ILMN_1681032). The null hypothesis of no causal effect is consistently rejected for *INO80E*, *KCTD13* and *YPEL3* under most combinations, indicating that these results do not emerge due to probe selection for analysis. Probes ILMN_1803277, ILMN_2402341 and ILMN_1751061 corresponding to (**A**) were used to report the analysis in the main text.

**Supplementary Figure S4**. Genome editing of *asphd1* using CRISPR/Cas9 to generate F0 zebrafish mutants. (**A**) Schematic of the asphd1 ortholog in zebrafish. The locus is shown with exons (black boxes); untranslated regions (white boxes); introns (dashed lines); guide (g)RNA target site and primers used to generate PCR products shown in panel B (red box and triangles, respectively). (**B**) Assessment of genome-editing efficiency using polyacrylamide gel electrophoresis (PAGE). Genomic DNA was extracted from single embryos at 2 dpf, and PCR amplified. PCR products were denatured, reannealed slowly and migrated on a 20% polyacrylamide gel. All twelve F0 embryos displayed heteroduplexes not present in two uninjected controls. Asterisks (*) indicate embryos assessed for percent mosaicism with pCR8/GW/TOPO-TA cloning and Sanger sequencing of individual clones. (**C**) Representative sequence alignments showing the most common targeting events for each embryo. To estimate percent mosaicism, one control and five F0 embryos were assessed (n=10-12 clones/embryo); a majority of F0 clones harbored deletions (green) and some clones harbored insertions (blue), suggesting ~96% mosaicism. gRNA target sequence (gray) and protospacer adjacent motif (PAM, red) are shown.

**Supplementary Figure S5. Global gene expression similarity of *ASPHD1* in human and mouse expression datasets.** The Multi Experiment Matrix web tool finds globally similarly expressed genes across thousands of (**A**) human (n=2,811) and (**B**) mouse (n=2,401) datasets. The top 100 datasets, in which *ASPHD1* (the input gene) has the highest standard deviation across samples are presented in the columns, while the top 150 genes with the expression pattern globally the most similar to *ASPHD1* are shown in the rows. The colored cells of the matrix illustrate the similarity of a given gene to *ASPHD1* in a specific dataset. Dark red denotes high similarity, white means no similarity and blue denotes opposite expression according to ranks calculated using Pearson correlation in each individual dataset. The matrix rows are sorted by the global gene similarity score calculated by Robust Rank Aggregation statistics, provided next to the colored matrix and gene name.

**Supplementary Figure S6 Comparative enrichment analyses in sets of *ASPHD1* globally similar genes in human and mouse.** Top-significant terms involved in synaptic function and location are highlighted and numbered in the Manhattan plot across eleven different data sources. The terms from different data sources are colored to ease the distinction between categories. The numbers next to terms indicate each term's position in both datasets and in the table below. The table includes selected term categories, identifiers, descriptions and adjusted p-values. The p-value cells are color-coded according to viridis color palette, highlighting smaller p-values with greener color. The results are available at **https://biit.cs.ut.ee/gplink/l/RfK8rsZHRf**

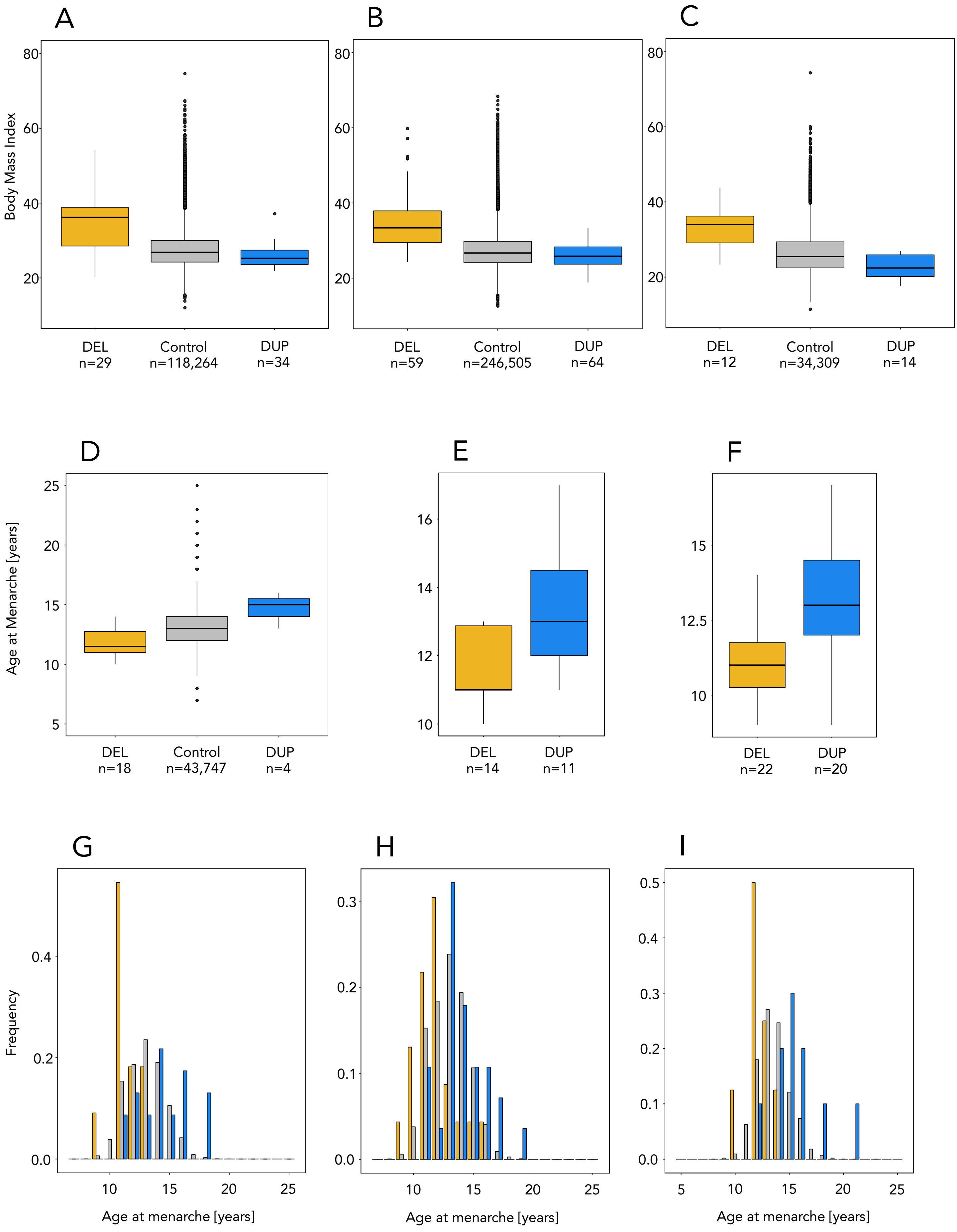
A
B
C
Body Mass Index
80
60
40
20
DEL
n=29
Control
n=118,264
DUP
n=34
DEL
n=59
Control
n=246,505
DUP
n=64
DEL
n=12
Control
n=34,309
DUP
n=14
D
E
F
Age at Menarche [years]
25
20
15
10
5
DEL
n=18
Control
n=43,747
DUP
n=4
16
14
12
10
DEL
n=14
DUP
n=11
15
12.5
10
DEL
n=22
DUP
n=20
G
H
I
Frequency
0.4
0.2
0.0
0.3
0.1
0.5
Age at menarche [years]
5
10
15
20
25

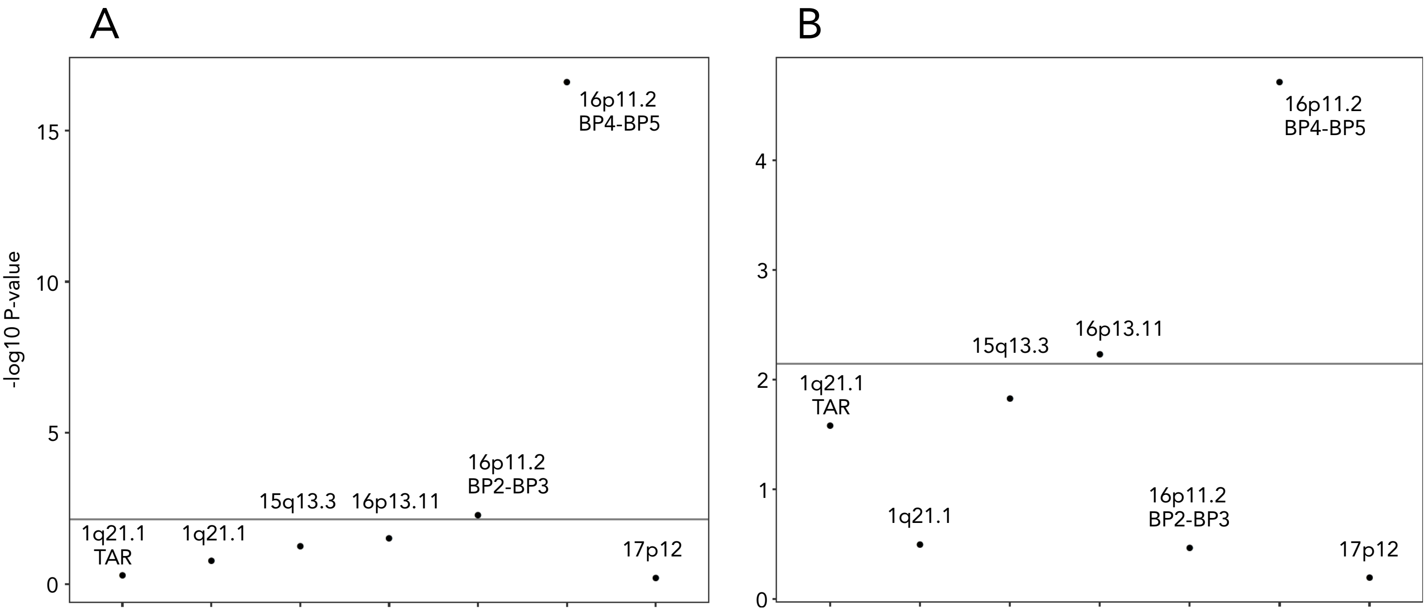
A
B
-log10 P-value
15
10
5
0
4
3
2
1
0
16p11.2
BP4-BP5
16p11.2
BP2-BP3
1q21.1
TAR
1q21.1
15q13.3
16p13.11
17p12
16p11.2
BP4-BP5
15q13.3
16p13.11
1q21.1
TAR
1q21.1
16p11.2
BP2-BP3
17p12

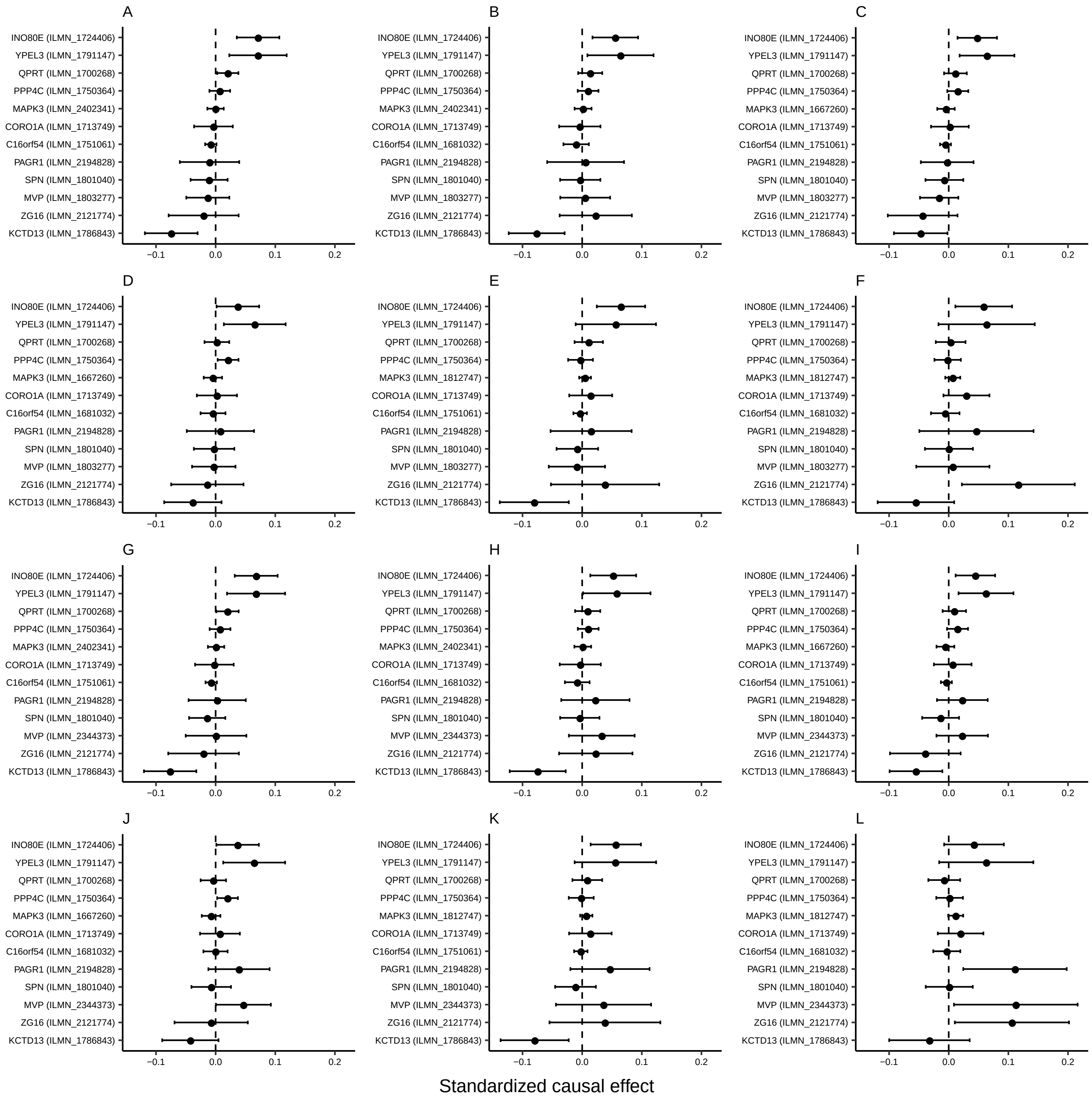
A
INO80E (ILMN_1724406)
YPEL3 (ILMN_1791147)
QPRT (ILMN_1700268)
PPP4C (ILMN_1750364)
MAPK3 (ILMN_2402341)
CORO1A (ILMN_1713749)
C16orf54 (ILMN_1751061)
PAGR1 (ILMN_2194828)
SPN (ILMN_1801040)
MVP (ILMN_1803277)
ZG16 (ILMN_2121774)
KCTD13 (ILMN_1786843)
−0.1 0.0 0.1 0.2
B
INO80E (ILMN_1724406)
YPEL3 (ILMN_1791147)
QPRT (ILMN_1700268)
PPP4C (ILMN_1750364)
MAPK3 (ILMN_2402341)
CORO1A (ILMN_1713749)
C16orf54 (ILMN_1681032)
PAGR1 (ILMN_2194828)
SPN (ILMN_1801040)
MVP (ILMN_1803277)
ZG16 (ILMN_2121774)
KCTD13 (ILMN_1786843)
−0.1 0.0 0.1 0.2
C
INO80E (ILMN_1724406)
YPEL3 (ILMN_1791147)
QPRT (ILMN_1700268)
PPP4C (ILMN_1750364)
MAPK3 (ILMN_1667260)
CORO1A (ILMN_1713749)
C16orf54 (ILMN_1751061)
PAGR1 (ILMN_2194828)
SPN (ILMN_1801040)
MVP (ILMN_1803277)
ZG16 (ILMN_2121774)
KCTD13 (ILMN_1786843)
−0.1 0.0 0.1 0.2
D
INO80E (ILMN_1724406)
YPEL3 (ILMN_1791147)
QPRT (ILMN_1700268)
PPP4C (ILMN_1750364)
MAPK3 (ILMN_1667260)
CORO1A (ILMN_1713749)
C16orf54 (ILMN_1681032)
PAGR1 (ILMN_2194828)
SPN (ILMN_1801040)
MVP (ILMN_1803277)
ZG16 (ILMN_2121774)
KCTD13 (ILMN_1786843)
−0.1 0.0 0.1 0.2
E
INO80E (ILMN_1724406)
YPEL3 (ILMN_1791147)
QPRT (ILMN_1700268)
PPP4C (ILMN_1750364)
MAPK3 (ILMN_1812747)
CORO1A (ILMN_1713749)
C16orf54 (ILMN_1751061)
PAGR1 (ILMN_2194828)
SPN (ILMN_1801040)
MVP (ILMN_1803277)
ZG16 (ILMN_2121774)
KCTD13 (ILMN_1786843)
−0.1 0.0 0.1 0.2
F
INO80E (ILMN_1724406)
YPEL3 (ILMN_1791147)
QPRT (ILMN_1700268)
PPP4C (ILMN_1750364)
MAPK3 (ILMN_1812747)
CORO1A (ILMN_1713749)
C16orf54 (ILMN_1681032)
PAGR1 (ILMN_2194828)
SPN (ILMN_1801040)
MVP (ILMN_1803277)
ZG16 (ILMN_2121774)
KCTD13 (ILMN_1786843)
−0.1 0.0 0.1 0.2
G
INO80E (ILMN_1724406)
YPEL3 (ILMN_1791147)
QPRT (ILMN_1700268)
PPP4C (ILMN_1750364)
MAPK3 (ILMN_2402341)
CORO1A (ILMN_1713749)
C16orf54 (ILMN_1751061)
PAGR1 (ILMN_2194828)
SPN (ILMN_1801040)
MVP (ILMN_2344373)
ZG16 (ILMN_2121774)
KCTD13 (ILMN_1786843)
−0.1 0.0 0.1 0.2
H
INO80E (ILMN_1724406)
YPEL3 (ILMN_1791147)
QPRT (ILMN_1700268)
PPP4C (ILMN_1750364)
MAPK3 (ILMN_2402341)
CORO1A (ILMN_1713749)
C16orf54 (ILMN_1681032)
PAGR1 (ILMN_2194828)
SPN (ILMN_1801040)
MVP (ILMN_2344373)
ZG16 (ILMN_2121774)
KCTD13 (ILMN_1786843)
−0.1 0.0 0.1 0.2
I
INO80E (ILMN_1724406)
YPEL3 (ILMN_1791147)
QPRT (ILMN_1700268)
PPP4C (ILMN_1750364)
MAPK3 (ILMN_1667260)
CORO1A (ILMN_1713749)
C16orf54 (ILMN_1751061)
PAGR1 (ILMN_2194828)
SPN (ILMN_1801040)
MVP (ILMN_2344373)
ZG16 (ILMN_2121774)
KCTD13 (ILMN_1786843)
−0.1 0.0 0.1 0.2
J
INO80E (ILMN_1724406)
YPEL3 (ILMN_1791147)
QPRT (ILMN_1700268)
PPP4C (ILMN_1750364)
MAPK3 (ILMN_1667260)
CORO1A (ILMN_1713749)
C16orf54 (ILMN_1681032)
PAGR1 (ILMN_2194828)
SPN (ILMN_1801040)
MVP (ILMN_2344373)
ZG16 (ILMN_2121774)
KCTD13 (ILMN_1786843)
−0.1 0.0 0.1 0.2
K
INO80E (ILMN_1724406)
YPEL3 (ILMN_1791147)
QPRT (ILMN_1700268)
PPP4C (ILMN_1750364)
MAPK3 (ILMN_1812747)
CORO1A (ILMN_1713749)
C16orf54 (ILMN_1751061)
PAGR1 (ILMN_2194828)
SPN (ILMN_1801040)
MVP (ILMN_2344373)
ZG16 (ILMN_2121774)
KCTD13 (ILMN_1786843)
−0.1 0.0 0.1 0.2
L
INO80E (ILMN_1724406)
YPEL3 (ILMN_1791147)
QPRT (ILMN_1700268)
PPP4C (ILMN_1750364)
MAPK3 (ILMN_1812747)
CORO1A (ILMN_1713749)
C16orf54 (ILMN_1681032)
PAGR1 (ILMN_2194828)
SPN (ILMN_1801040)
MVP (ILMN_2344373)
ZG16 (ILMN_2121774)
KCTD13 (ILMN_1786843)
−0.1 0.0 0.1 0.2
Standardized causal effect

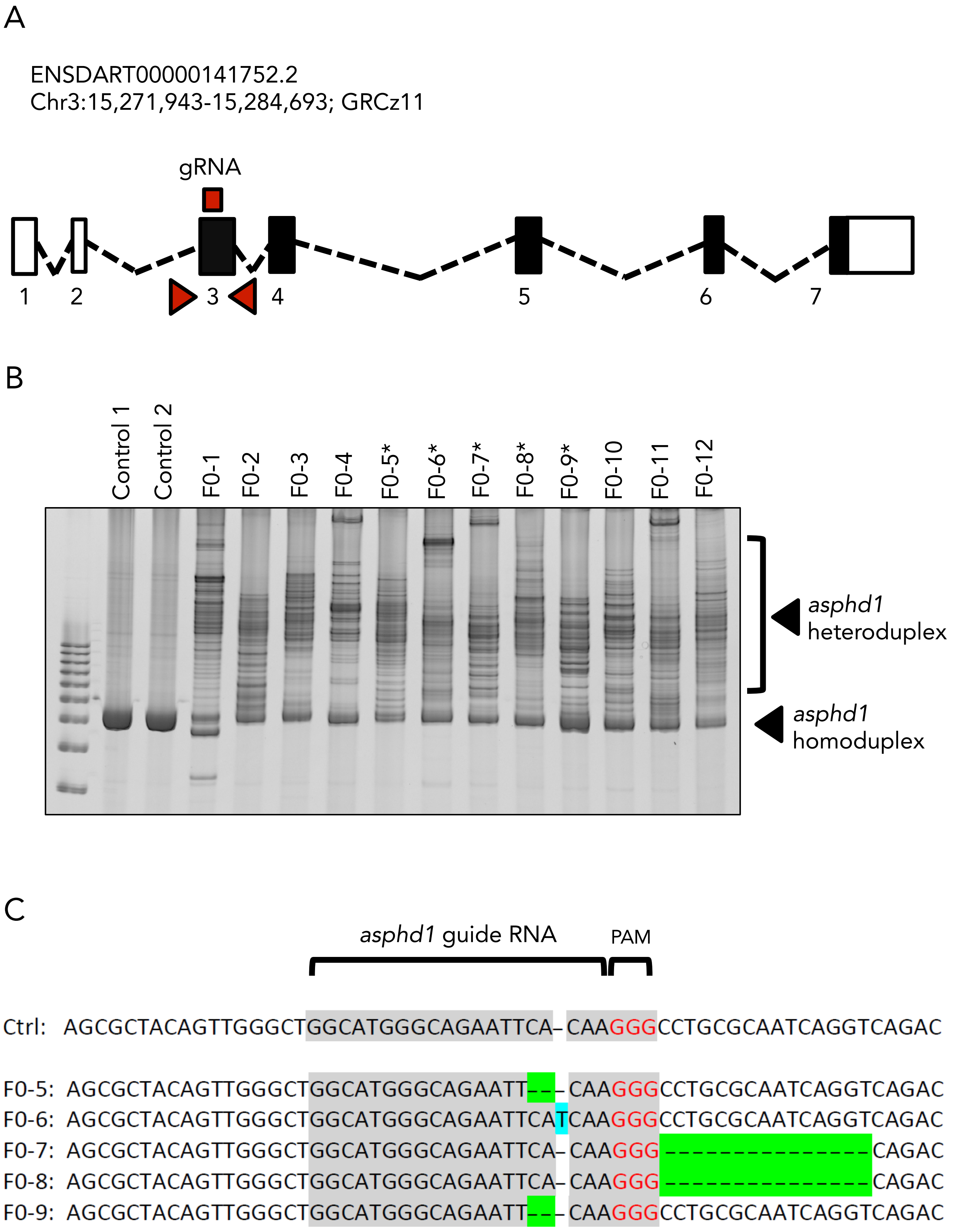
A
ENSDART00000141752.2
Chr3:15,271,943-15,284,693; GRCz11
gRNA
1
2
3
4
5
6
7
B
Control 1
Control 2
F0-1
F0-2
F0-3
F0-4
F0-5*
F0-6*
F0-7*
F0-8*
F0-9*
F0-10
F0-11
F0-12
asphd1 heteroduplex
asphd1 homoduplex
C
asphd1 guide RNA
PAM
Ctrl: AGCGCTACAGTTGGGCTGGCATGGGCAGAATTCA–CAAGGGCCTGCGCAATCAGGTCAGAC
F0-5: AGCGCTACAGTTGGGCTGGCATGGGCAGAATT–––CAAGGGCCTGCGCAATCAGGTCAGAC
F0-6: AGCGCTACAGTTGGGCTGGCATGGGCAGAATTCATCAAGGGCCTGCGCAATCAGGTCAGAC
F0-7: AGCGCTACAGTTGGGCTGGCATGGGCAGAATTCA–CAAGGG–––––––––––––––CAGAC
F0-8: AGCGCTACAGTTGGGCTGGCATGGGCAGAATTCA–CAAGGG–––––––––––––––CAGAC
F0-9: AGCGCTACAGTTGGGCTGGCATGGGCAGAATT–––CAAGGGCCTGCGCAATCAGGTCAGAC

A

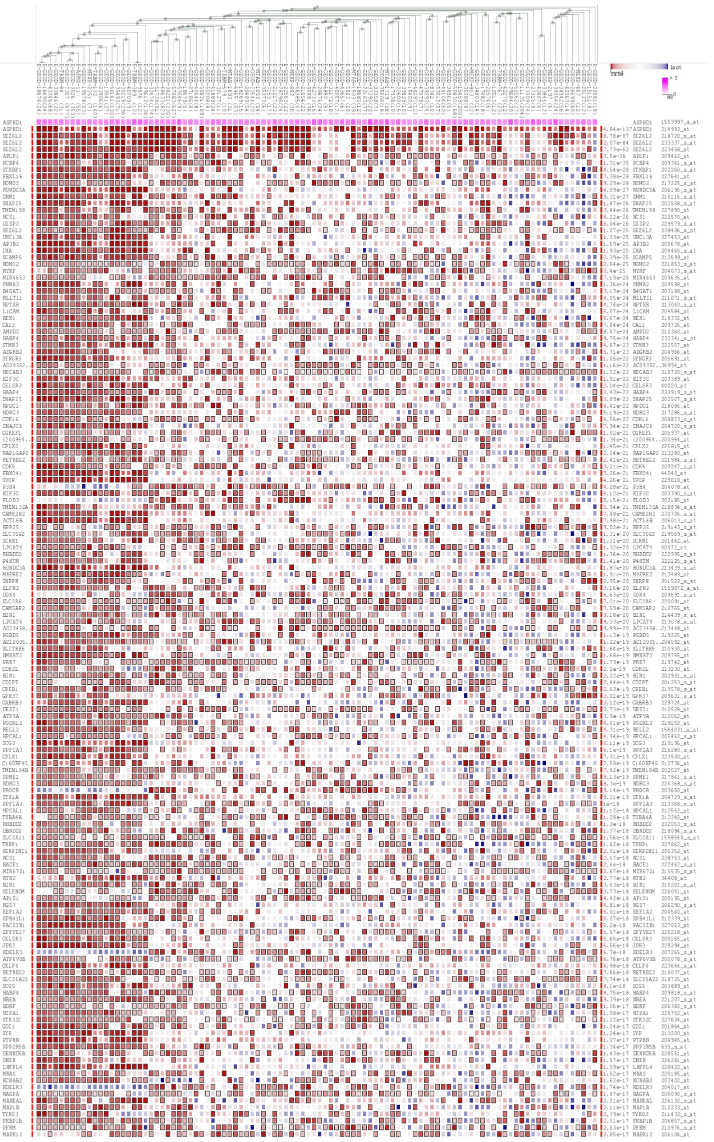

B

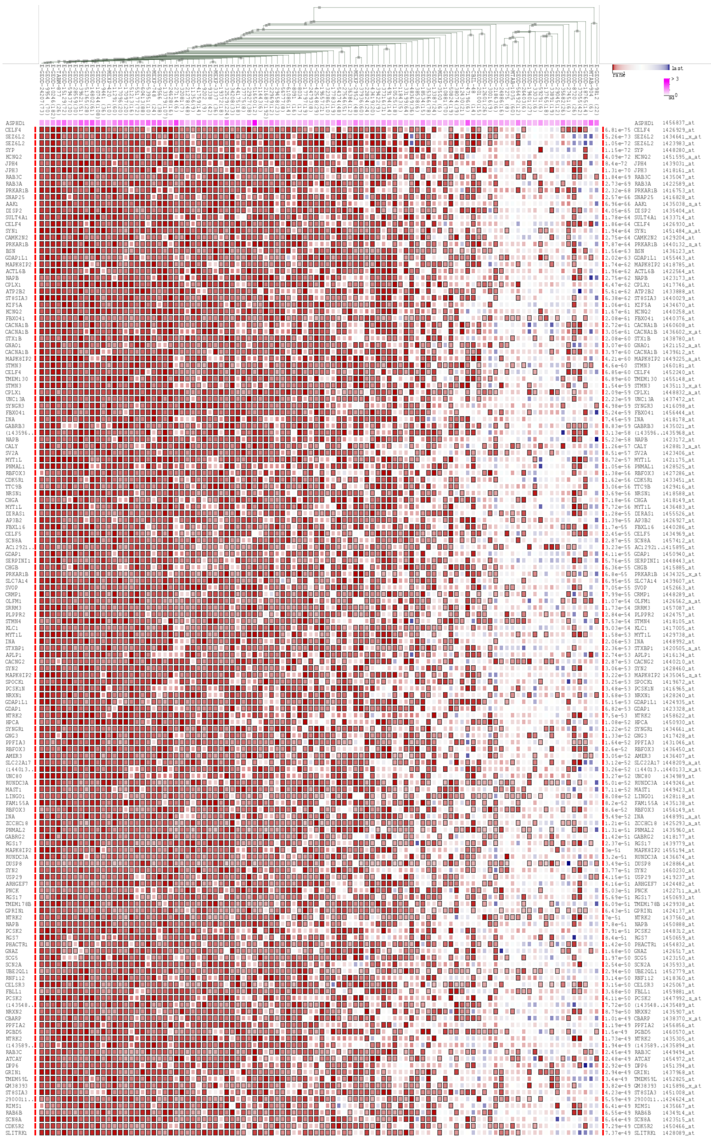

> human

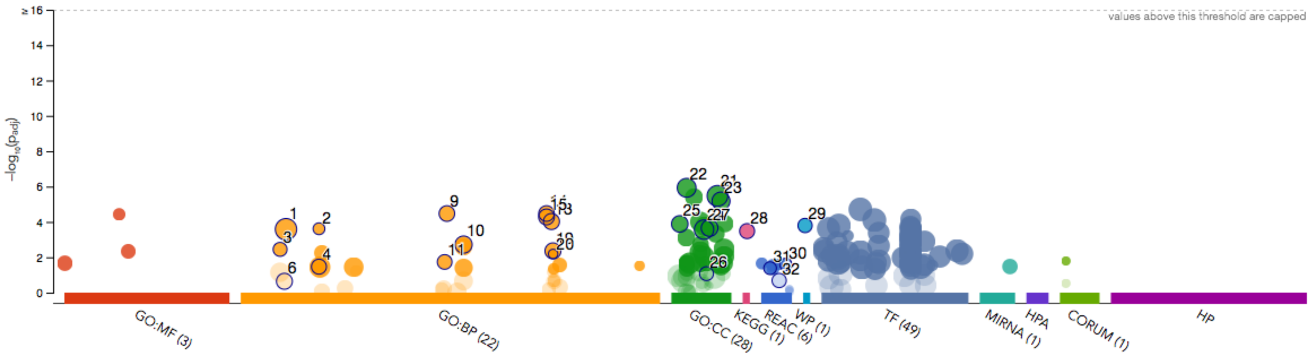


> mouse

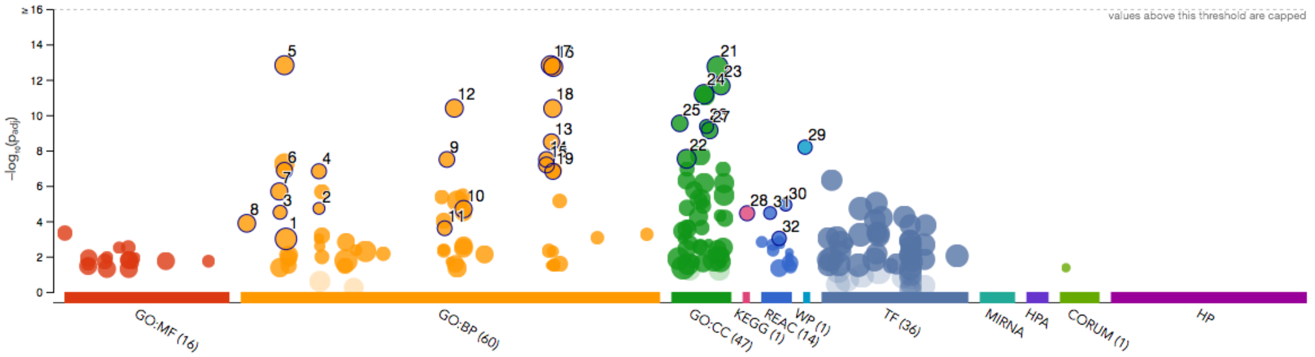


| id | source | term id | term name | $p_{adj}$ (human) | $p_{adj}$ (mouse) |
|---|---|---|---|---|---|
| 1 | GO:BP | GO:0007399 | nervous system development | $2.593\times10^{-4}$ | $9.955\times10^{-4}$ |
| 2 | GO:BP | GO:0016082 | synaptic vesicle priming | $2.434\times10^{-4}$ | $1.886\times10^{-5}$ |
| 3 | GO:BP | GO:0006904 | vesicle docking involved in exocytosis | $3.566\times10^{-3}$ | $3.118\times10^{-5}$ |
| 4 | GO:BP | GO:0016079 | synaptic vesicle exocytosis | $3.394\times10^{-2}$ | $1.490\times10^{-7}$ |
| 5 | GO:BP | GO:0007268 | chemical synaptic transmission | ~~1.000~~ | $1.498\times10^{-13}$ |
| 6 | GO:BP | GO:0007269 | neurotransmitter secretion | ~~$2.277\times10^{-1}$~~ | $1.337\times10^{-7}$ |
| 7 | GO:BP | GO:0006836 | neurotransmitter transport | ~~1.000~~ | $2.063\times10^{-6}$ |
| 8 | GO:BP | GO:0001505 | regulation of neurotransmitter levels | ~~1.000~~ | $1.310\times10^{-4}$ |
| 9 | GO:BP | GO:0048489 | synaptic vesicle transport | $3.353\times10^{-5}$ | $3.209\times10^{-8}$ |
| 10 | GO:BP | GO:0051650 | establishment of vesicle localization | $1.912\times10^{-3}$ | $2.054\times10^{-5}$ |
| 11 | GO:BP | GO:0048278 | vesicle docking | $1.836\times10^{-2}$ | $2.495\times10^{-4}$ |
| 12 | GO:BP | GO:0050804 | modulation of chemical synaptic transmission | ~~1.000~~ | $4.047\times10^{-11}$ |
| 13 | GO:BP | GO:0099003 | vesicle-mediated transport in synapse | $9.480\times10^{-5}$ | $3.259\times10^{-9}$ |
| 14 | GO:BP | GO:0097480 | establishment of synaptic vesicle localization | $3.353\times10^{-5}$ | $3.209\times10^{-8}$ |
| 15 | GO:BP | GO:0097479 | synaptic vesicle localization | $5.219\times10^{-5}$ | $6.544\times10^{-8}$ |
| 16 | GO:BP | GO:0099536 | synaptic signaling | ~~1.000~~ | $1.867\times10^{-13}$ |
| 17 | GO:BP | GO:0098916 | anterograde trans-synaptic signaling | ~~1.000~~ | $1.498\times10^{-13}$ |
| 18 | GO:BP | GO:0099177 | regulation of trans-synaptic signaling | ~~1.000~~ | $4.195\times10^{-11}$ |
| 19 | GO:BP | GO:0099504 | synaptic vesicle cycle | $4.351\times10^{-3}$ | $1.513\times10^{-7}$ |
| 20 | GO:BP | GO:0099525 | presynaptic dense core vesicle exocytosis | $6.483\times10^{-3}$ | ~~1.000~~ |
| 21 | GO:CC | GO:0097458 | neuron part | $3.338\times10^{-6}$ | $1.781\times10^{-13}$ |
| 22 | GO:CC | GO:0030424 | axon | $1.172\times10^{-6}$ | $2.949\times10^{-8}$ |
| 23 | GO:CC | GO:0098793 | presynapse | $6.589\times10^{-6}$ | $2.177\times10^{-12}$ |
| 24 | GO:CC | GO:0044456 | synapse part | $2.724\times10^{-4}$ | $6.561\times10^{-12}$ |
| 25 | GO:CC | GO:0008021 | synaptic vesicle | $1.298\times10^{-4}$ | $2.908\times10^{-10}$ |
| 26 | GO:CC | GO:0048786 | presynaptic active zone | ~~$8.427\times10^{-2}$~~ | $4.388\times10^{-10}$ |
| 27 | GO:CC | GO:0070382 | exocytic vesicle | $2.213\times10^{-4}$ | $7.243\times10^{-10}$ |
| 28 | KEGG | KEGG:04721 | Synaptic vesicle cycle | $3.302\times10^{-4}$ | $3.542\times10^{-5}$ |
| 29 | WP | WP:WP2267 | Synaptic Vesicle Pathway | $1.592\times10^{-4}$ | $6.486\times10^{-9}$ |
| 30 | REAC | REAC:R-HSA-1... | Serotonin Neurotransmitter Release Cycle | $2.508\times10^{-2}$ | $1.195\times10^{-5}$ |
| 31 | REAC | REAC:R-HSA-2... | Dopamine Neurotransmitter Release Cycle | $4.138\times10^{-2}$ | $3.435\times10^{-5}$ |
| 32 | REAC | REAC:R-HSA-1... | Neurotransmitter release cycle | ~~$2.060\times10^{-1}$~~ | $9.334\times10^{-4}$ |

| | |
|---|---|
| **version** | e95_eg42_p13_f6e58b9 |
| **date** | 5/24/2019, 1:41:37 PM |
| **organism** | hsapiens |

g:Profiler

### Supplementary Table S1. Summary characteristics in study cohorts

#### Adult general population cohorts

| | Controls | | | | Deletion carriers | | | | Duplication carriers | | | |
|---|---|---|---|---|---|---|---|---|---|---|---|---|
| | N | Mean age at recruitment | Mean BMI | Mean AaM | N | Mean age at recruitment | Mean BMI | Mean AaM | N | Mean age at recruitment | Mean BMI | Mean AaM |
| **UK Biobank 1st release Europeans** | | | | | | | | | | | | |
| Males+females | 118 264 | 56.9 | 27.5 | | 29 | 54.9 | 35.2 | | 34 | 60 | 25.9 | |
| Males | 56 047 | 57.3 | 28 | | 18 | 54.8 | 34.9 | | 11 | 59.1 | 26.3 | |
| Females | 62 217 | 56.6 | 27.2 | 12.9 | 11 | 55 | 35.8 | 11.4 | 23 | 60.5 | 25.7 | 14.4 |
| **UK Biobank 1st release all ethnicities** | | | | | | | | | | | | |
| Males+females | 146 569 | 56.6 | 27.5 | | 42 | 54 | 35.2 | | 40 | 60.1 | 25.7 | |
| Males | 68 993 | 57 | 27.9 | | 22 | 54.9 | 34.7 | | 14 | 60.4 | 26.7 | |
| Females | 77 576 | 56.3 | 27.2 | 13 | 20 | 53.2 | 35.7 | 11.6 | 26 | 59.9 | 25.3 | 14.5 |
| **UK Biobank 2nd release Europeans** | | | | | | | | | | | | |
| Males+females | 246 505 | 57 | 27.4 | | 59 | 53.2 | 35.1 | | 64 | 55.5 | 26 | |
| Males | 112 890 | 57.1 | 27.8 | | 36 | 53.8 | 34.7 | | 36 | 55.7 | 25.8 | |
| Females | 133 615 | 56.8 | 27 | 13 | 23 | 52.3 | 35.8 | 11.9 | 28 | 55.3 | 26.3 | 14 |
| **UK Biobank 2nd release all ethnicities** | | | | | | | | | | | | |
| Males+females | 297 830 | 56.5 | 27.4 | | 77 | 53.1 | 34.9 | | 70 | 55.5 | 25.9 | |
| Males | 135 820 | 56.7 | 27.8 | | 45 | 53.7 | 34 | | 41 | 55.9 | 25.7 | |
| Females | 162 010 | 56.4 | 27 | 13 | 32 | 52.2 | 36.2 | 11.9 | 29 | 54.9 | 26.1 | 14 |
| **UK Biobank full release non-Europeans** | | | | | | | | | | | | |
| Males+females | 79 617 | 54.8 | 27.4 | | 31 | 52.4 | 34.6 | | 12 | 57.7 | 24.6 | |
| Males | 35 870 | 54.9 | 27.8 | | 13 | 53.8 | 31.8 | | 8 | 60.1 | 26 | |
| Females | 43 747 | 54.7 | 27.2 | 13 | 18 | 51.4 | 36.4 | 11.9 | 4 | 52.8 | 22.1 | 14.7 |
| **EGCUT** | | | | | | | | | | | | |
| Males+females | 34 309 | 44.8 | 26.3 | | 12 | 36.5 | 32.9 | | 14 | 40.1 | 22.8 | |
| Males | 11 955 | 44.4 | 26.5 | | 4 | 35 | 30.2 | | 4 | 43.5 | 24.5 | |
| Females | 22 354 | 45.1 | 26.2 | 13.5 | 8 | 37.2 | 34.3 | 12.2 | 10 | 38.7 | 22.1 | 15.6 |

#### 16p11.2 clinical cohorts

| | Unrelated post-pubertal deletion carriers | | | | Unrelated post-pubertal duplication carriers | | | |
|---|---|---|---|---|---|---|---|---|
| | N | Mean age at recruitment | Mean BMI z-score | Mean AaM | N | Mean age at recruitment | Mean BMI z-score | Mean AaM |
| **Simons VIP cohort all ethnicities** | | | | | | | | |
| Pediatric males (<18 years) | 21 | 14.8 | 1.12 | | 6 | 15.6 | 0.04 | |
| Adult males (>18 years) | 6 | 41.6 | 2.5 | | 18 | 39.1 | 0.69 | |
| Females | 22 | 17.2 | 1.41 | 11 | 20 | 33.9 | 0.39 | 13.3 |
| **European 16p11.2 cohort all ethicities** | | | | | | | | |
| Females | 14 | 33.85 | 1.8 | 11.6 | 11 | 38 | 0.3 | 13.5 |

#### Supplementary Table S2. Association with BMI in adult population cohorts

| | Mean BMI | | | ANOVA P-values | | | T-test P-values | |
|---|---|---|---|---|---|---|---|---|
| | Controls | Deletions | Duplications | Three groups | DEL vs Ctrl | DUP vs Ctrl | DEL vs Ctrl | DUP vs Ctrl |
| **UK Biobank 1st release Europeans** | | | | | | | | |
| Males+females (adjusted for sex) | 27.54 | 35.23 | 25.86 | 2.48E-17 | 1.65E-17 | 0.047 | 4.43E-05 | 3.70E-03 |
| Males | 27.95 | 34.89 | 26.27 | 2.72E-11 | 7.54E-12 | 0.18 | 0.0013 | 0.13 |
| Females | 27.17 | 35.79 | 25.66 | 8.96E-08 | 3.89E-08 | 0.13 | 0.019 | 0.029 |
| **UK Biobank 1st release all ethnicities** | | | | | | | | |
| Males+females (adjusted for sex) | 27.54 | 35.17 | 25.73 | 5.33E-25 | 5.95E-25 | 0.021 | 6.99E-07 | 2.50E-03 |
| Males | 27.94 | 34.67 | 26.67 | 1.25E-12 | 2.48E-13 | 0.27 | 0.00022 | 0.24 |
| Females | 27.19 | 35.72 | 25.26 | 2.10E-13 | 1.57E-13 | 0.049 | 0.0011 | 0.007 |
| **UK Biobank 2nd release Europeans** | | | | | | | | |
| Males+females (adjusted for sex) | 27.35 | 35.11 | 26.01 | 5.20E-36 | 5.37E-36 | 0.018 | 2.56E-09 | 2.60E-03 |
| Males | 27.79 | 34.67 | 25.82 | 6.15E-23 | 2.44E-22 | 0.0053 | 6.23E-05 | 2.10E-04 |
| Females | 26.98 | 35.79 | 26.26 | 1.71E-16 | 2.06E-17 | 0.48 | 1.48E-05 | 3.60E-01 |
| **UK Biobank 2nd release all ethnicities** | | | | | | | | |
| Males+females (adjusted for sex) | 27.36 | 34.91 | 25.88 | 3.81E-44 | 7.20E-44 | 0.0083 | 2.88E-11 | 4.50E-04 |
| Males | 27.78 | 33.96 | 25.71 | 3.04E-23 | 3.07E-22 | 0.0019 | 3.23E-05 | 3.02E-05 |
| Females | 27 | 36.19 | 26.12 | 4.21E-24 | 4.64E-25 | 0.4 | 2.14E-07 | 2.50E-01 |
| **UK Biobank full release non-Europeans** | | | | | | | | |
| Males+females (adjusted for sex) | 27.45 | 34.58 | 24.62 | 2.07E-16 | 1.07E-16 | 0.066 | 6.66E-05 | 4.60E-02 |
| Males | 27.78 | 31.79 | 26.04 | 0.0034 | 1.30E-03 | 0.32 | 5.30E-02 | 2.80E-01 |
| Females | 27.18 | 36.43 | 22.13 | 1.90E-14 | 1.22E-14 | 0.055 | 5.10E-04 | 6.90E-02 |
| **EGCUT** | | | | | | | | |
| Males+females (adjusted for sex) | 26.32 | 32.9 | 22.75 | 6.35E-08 | 1.40E-07 | 0.02 | 4.40E-03 | 1.10E-03 |
| Males | 26.54 | 30.16 | 24.49 | 0.12 | 5.80E-02 | 0.41 | 2.30E-01 | 1.00E-01 |
| Females | 26.21 | 34.27 | 22.06 | 3.17E-07 | 4.79E-07 | 0.032 | 1.30E-02 | 4.10E-03 |

#### Supplementary Table S3. Association with AaM in adult population cohorts

| | Mean AaM | | | Wilcoxon P-values | | Kruskal-Wallis | ANOVA (BMI and YoB adjusted) | | |
|---|---|---|---|---|---|---|---|---|---|
| | Controls | Deletions | Duplications | Del vs Ctrl | Dup vs Ctrl | Three groups | Three groups | Del vs Ctrl | Dup vs Ctrl |
| **UK Biobank 1st release** | | | | | | | | | |
| Europeans | 12.9 | 11.4 | 14.4 | 0.00098 | 0.00203 | 3.74E-05 | 1.93E-05 | 0.013 | 7.84E-05 |
| All ethnicities | 13 | 11.6 | 14.5 | 0.00015 | 0.0004 | 1.42E-06 | 3.16E-06 | 0.01 | 1.55E-05 |
| **UK Biobank 2nd release** | | | | | | | | | |
| Europeans | 13 | 11.9 | 14 | 0.0016 | 0.0063 | 0.00017 | 0.00065 | 0.045 | 0.0011 |
| All ethnicities | 13 | 11.9 | 14 | 0.00024 | 0.008 | 3.45E-05 | 0.00083 | 0.037 | 0.0017 |
| **UK Biobank full release** | | | | | | | | | |
| Non-Europeans | 13 | 11.9 | 14.7 | 0.0056 | 0.093 | 0.0052 | 0.13 | 0.19 | 0.13 |
| **EGCUT** | | | | | | | | | |
| Europeans | 13.5 | 12.3 | 15.6 | 0.017 | 0.0014 | 0.00037 | 2.40E-05 | 0.19 | 9.84E-06 |

### Supplementary Table S4.Onset of pubertal traits in male CNV carriers

#### Adult general population cohorts

| | Controls | | | | Deletion carriers | | | | Duplication carriers | | | | Fisher's exact test P-values | |
|---|---|---|---|---|---|---|---|---|---|---|---|---|---|---|
| | "younger" | "average" | "older" | NAs | "younger" | "average" | "older" | NAs | "younger" | "average" | "older" | NAs | Del vs Ctrl "younger" | Dup vs Ctrl "older" |
| **UK Biobank 1st release Europeans** | | | | | | | | | | | | | | |
| Voice break | 2,231 (4.3%) | 46,700 (89.9%) | 2,992 (5.76%) | 4 124 | 0 | 15 (100%) | 0 | 3 | 0 | 9 (90%) | 1 (10%) | 1 | 1 | 0.46 |
| Facial hair | 3,640 (6.71%) | 43,652 (80.4%) | 6,985 (12.9%) | 1 770 | 2 (11.8%) | 15 (88.2%) | 0 | 1 | 1 (9.09%) | 6 (54.5%) | 4 (36.4%) | 0 | 0.38 | 0.038 |
| **UK Biobank 1st release all ethnicities** | | | | | | | | | | | | | | |
| Voice break | 2,815 (4.45%) | 56,845 (89.8%) | 3,631 (5.74%) | 5 702 | 2 (11.1%) | 16 (88.9%) | 0 | 4 | 0 | 11 (91.7%) | 1 (8.33%) | 2 | 0.21 | 0.52 |
| Facial hair | 4,592 (6.93%) | 53,210 (80.3%) | 8,498 (12.8%) | 2 693 | 4 (19.0%) | 17 (81.0%) | 0 | 1 | 1 (7.14%) | 9 (64.3%) | 4 (28.6%) | 0 | 0.08 | 0.09 |
| **UK Biobank 2nd release Europeans** | | | | | | | | | | | | | | |
| Voice break | 4,379 (4.19%) | 93,987 (89.9%) | 6,225 (7.93%) | 8 299 | 2 (6.90%) | 27 (93.1%) | 0 | 7 | 4 (13.8%) | 21 (72.4%) | 4 (13.8%) | 7 | 0.37 | 0.066 |
| Facial hair | 6,963 (6.37%) | 88,177 (80.7%) | 14,092 (12.9%) | 3 658 | 7 (22.6%) | 24 (77.4%) | 0 | 5 | 1 (3.03%) | 26 (78.8%) | 6 (18.2%) | 3 | 0.0062 | 0.44 |
| **UK Biobank 2nd release all ethnicities** | | | | | | | | | | | | | | |
| Voice break | 5,419 (4.37%) | 111,199 (89.7%) | 7,347 (5.93%) | 11 855 | 2 (5.56%) | 34 (94.4%) | 0 | 9 | 4 (12.5%) | 23 (71.9%) | 5 (15.6%) | 9 | 0.68 | 0.027 |
| Facial hair | 8,767 (6.75%) | 104,592 (80.5%) | 16,545 (12.7%) | 5 916 | 7 (18.4%) | 31 (81.6%) | 0 | 7 | 1 (2.78%) | 28 (77.8%) | 7 (19.4%) | 5 | 0.025 | 0.32 |
| **UK Biobank full release non-Europeans** | | | | | | | | | | | | | | |
| Voice break | 1,624 (5.28%) | 27,352 (89%) | 1,760 (5.73%) | 5 134 | 2 (20%) | 8 (80%) | 0 | 3 | 0 | 4 (80%) | 1 (20%) | 3 | 0.1 | 0.27 |
| Facial hair | 2,755 (8.43%) | 25,970 (79.4%) | 3,964 (12.1%) | 3 181 | 2 (18.2%) | 9 (81.8%) | 0 | 2 | 0 | 5 (83.3%) | 1 (16.7%) | 2 | 0.29 | 0.57 |

#### 16p11.2 clinical cohorts

| | Unadjusted | | | | | | |
|---|---|---|---|---|---|---|---|
| | Deletion | | Duplication | | | | |
| | N | Mean | N | Mean | Delta (Dup-Del) | P-value (genotype) | Observations deleted due to missingness |
| **Simons VIP pediatric males** | | | | | | | |
| Facial hair | 16 | 12.4 | 5 | 14 | +1.6 | 0.04 | 6 |
| Growth spurt | 17 | 11.2 | 4 | 12.5 | +1.3 | 0.16 | 6 |
| Body hair | 18 | 11.2 | 5 | 12.6 | +1.4 | 0.09 | 4 |
| Skin changes | 13 | 11.8 | 5 | 13 | +1.2 | 0.15 | 9 |
| Voice change | 16 | 11.9 | 4 | 13.5 | +1.6 | 0.05 | 7 |
| **Simons VIP adult males** | | | | | | | |
| Facial hair | 6 | 14.5 | 18 | 16.6 | +2.1 | 0.11 | 0 |
| Growth spurt | 6 | 11.7 | 18 | 14.2 | +2.5 | 0.01 | 0 |
| Body hair | 6 | 11.5 | 18 | 13.8 | +2.3 | 0.04 | 0 |
| Skin changes | 5 | 13.2 | 18 | 14.5 | +1.3 | 0.17 | 1 |
| Voice change | 6 | 13.2 | 18 | 15.1 | +1.9 | 0.09 | 0 |

| | Adjusted for BMI | | | | | | | | | | |
|---|---|---|---|---|---|---|---|---|---|---|---|
| | Deletion | | | Duplication | | | | | | | |
| | N | Mean (puberty) | Mean (bmi z-score) | N | Mean (puberty) | Mean (bmi z-score) | Delta (puberty) | P-value (genotype) | Delta (bmi z-score) | P-value (BMI) | Observations deleted due to missingness |
| **Simons VIP pediatric males** | | | | | | | | | | | |
| Facial hair | 11 | 12.6 | 1.17 | 4 | 13.5 | 0.04 | +0.9 | 0.43 | -1.13 | 0.91 | 12 |
| Growth spurt | 13 | 11.5 | 1.22 | 4 | 12.5 | 0.04 | +1.0 | 0.29 | -1.18 | 0.91 | 10 |
| Body hair | 13 | 11.3 | 1.06 | 4 | 12 | 0.04 | +0.7 | 0.55 | -1.02 | 0.87 | 10 |
| Skin changes | 9 | 12 | 0.65 | 4 | 13.8 | 0.04 | +1.8 | 0.07 | -0.61 | 0.94 | 14 |
| Voice change | 11 | 12.1 | 1.06 | 4 | 13.5 | 0.04 | +1.4 | 0.23 | -1.02 | 0.6 | 12 |
| **Simons VIP adult males** | | | | | | | | | | | |
| Facial hair | 4 | 13.8 | 2.51 | 17 | 16.7 | 0.69 | +2.9 | 0.28 | -1.82 | 0.52 | 3 |
| Growth spurt | 4 | 11.8 | 2.51 | 17 | 14.2 | 0.69 | +2.4 | 0.48 | -1.82 | 0.14 | 3 |
| Body hair | 4 | 10.8 | 2.51 | 17 | 13.8 | 0.69 | +3.0 | 0.28 | -1.82 | 0.19 | 3 |
| Skin changes | 4 | 13.3 | 2.51 | 17 | 14.5 | 0.69 | +1.2 | 0.48 | -1.82 | 0.76 | 3 |
| Voice change | 4 | 13 | 2.51 | 17 | 15.1 | 0.69 | +2.1 | 0.25 | -1.82 | 0.98 | 3 |

### Supplementary Table S5. Reproductive traits and diagnoses in female CNV carriers

| | N (with phenotype value available) | | | Mean (or frequency) | | | | | |
|---|---|---|---|---|---|---|---|---|---|
| | Control | Deletion | Duplication | Control | Deletion | Duplication | P control vs deletion | P control vs duplication | Test |
| **UK Biobank 1st release Europeans** | | | | | | | | | |
| Has had miscarriage | 62 148 | 11 | 23 | 0.2046 | 0.091 (1/11) | 0.39 (9/23) | 0.71 | 0.037 | Fisher's exact test |
| Number of live births | 62 174 | 11 | 23 | 1.81 | 0.91 | 2.09 | 0.013 | 0.49 | Wilcoxon rank-sum test |
| Age at first birth | 50 618 | 4 | 19 | 25.73 | 26 | 24.53 | 0.91 | 0.2 | Wilcoxon rank-sum test |
| Age at last birth | 50 562 | 4 | 16 | 29.76 | 33 | 30.75 | 0.23 | 0.5 | Wilcoxon rank-sum test |
| Age at natural menopause | 23 551 | 3 | 10 | 49.97 | 48.33 | 49.3 | 0.64 | 0.31 | Wilcoxon rank-sum test |
| **UK Biobank 2nd release Europeans** | | | | | | | | | |
| Has had miscarriage | 133 431 | 23 | 28 | 0.2047 | 0.17 (4/23) | 0.29 (8/28) | 1 | 0.35 | Fisher's exact test |
| Number of live births | 133 551 | 23 | 28 | 1.81 | 0.78 | 1.71 | 2.89E-05 | 0.54 | Wilcoxon rank-sum test |
| Age at first birth | 109 239 | 10 | 20 | 26.01 | 24.6 | 22.65 | 0.48 | 0.00034 | Wilcoxon rank-sum test |
| Age at last birth | 109 109 | 10 | 18 | 30.01 | 27.5 | 28.83 | 0.094 | 0.28 | Wilcoxon rank-sum test |
| Age at menopause | 51 077 | 4 | 7 | 50.15 | 49.5 | 47 | 0.73 | 0.098 | Wilcoxon rank-sum test |
| **UK Biobank full release non-Europeans** | | | | | | | | | |
| Has had miscarriage | 43 617 | 18 | 4 | 0.2166 | 0.056 (1/18) | 0.5 (2/4) | 0.15 | 0.21 | Fisher's exact test |
| Number of live births | 43 568 | 18 | 4 | 1.83 | 1.28 | 2.25 | 0.087 | 0.14 | Wilcoxon rank-sum test |
| Age at first birth | 33 849 | 11 | 4 | 26.23 | 23.82 | 25.75 | 0.14 | 0.93 | Wilcoxon rank-sum test |
| Age at last birth | 33 714 | 11 | 3 | 30.91 | 28.36 | 31 | 0.29 | 0.95 | Wilcoxon rank-sum test |
| Age at menopause | 15 629 | 3 | 0 | 49.81 | 47.33 | - | 0.32 | - | Wilcoxon rank-sum test |

| Diagnosis name | Diagnosis code (ICD-10) | N (with phenotype data available) | | | | Frequency | | | | Fisher's exact test | | | | | |
|---|---|---|---|---|---|---|---|---|---|---|---|---|---|---|---|
| | | Control | Deletion | Duplication | Both CNVs combined | Control | Deletion | Duplication | CNV | P-value (deletion) | OR (95%CI) | P-value (duplication) | OR (95%CI) | p-value (CNV) | OR (95%CI) |
| **EGCUT female Europeans** | | | | | | | | | | | | | | | |
| Absent, scanty and rare menstruation | N91 | 34,054 | 6 | 11 | 17 | 0.087 | 0.333 (2/6) | 0.273 (3/11) | 0.294 (5/17) | 0.09 | 5.2 (0.5; 36.6) | 0.06 | 3.9 (0.7; 16.4) | 0.013 | 4.4 (1.2; 13.3) |
| Ovarian dysfunction | E28 | 34,054 | 6 | 11 | 17 | 0.103 | 0.167 (1/6) | 0.273 (3/11) | 0.235 (4/17) | - | - | 0.1 | 3.3 (0.6; 13.6) | 0.09 | 2.7 (0.6; 8.7) |
| Female infertility | N97 | 34,054 | 6 | 11 | 17 | 0.055 | 0.167 (1/6) | 0.091 (1/11) | 0.118 (2/17) | - | - | | | 0.23 | 2.3 (0.3; 9.9) |
| Cystitis | N30 | 34,054 | 6 | 11 | 17 | 0.413 | 0.167 (1/6) | 0.727 (8/11) | 0.529 (9/17) | - | - | 0.06 | 3.8 (0.9; 22.1) | 0.34 | 1.6 (0.5; 4.8) |
| Inflammatory diseases of female pelvic organs | N70-N77 | 34,054 | 6 | 11 | 17 | 0.444 | 0.667 (4/6) | 0.818 (9/11) | 0.765 (13/17) | 0.42 | 2.5 (0.4; 27.7) | 0.015 | 5.6 (1.2; 53.6) | 0.012 | 4.1 (1.3; 17.1) |
| Endometriosis | N80 | 34,054 | 6 | 11 | 17 | 0.042 | - | 0.182 (2/11) | 0.118 (2/17) | - | - | 0.07 | 5.1 (0.5; 24.6) | 0.16 | 3.1 (0.3; 13.2) |
| Noninlammatory disorders of ovary, fallopian tube and broad ligament | N83 | 34,054 | 6 | 11 | 17 | 0.094 | 0.333 (2/6) | 0.364 (4/11) | 0.353 (6/17) | 0.1 | 4.8 (0.4; 33.6) | 0.015 | 5.6 (1.2; 21.6) | 0.003 | 5.2 (1.6; 15.5) |

#### Individual female 16p11.2 CNV carriers in EGCUT cohort

| Sample ID | CNV | Gender | Age at recruitment | Major reproductive diagnoses (ICD-10) | Other diagnoses related to reproduction and the genitourinary system (ICD-10) |
|---|---|---|---|---|---|
| EGCUT_1 | DEL | F | 38 | N92.6, irrerular menstruation; N95.1, menopausal state | N72, inflammatory disease of cervix uteri; N81.2, incomplete uterovaginal prolapse; N83.2, unspecified ovarian cysts; N84.0, polyp of corpus uteri; N85.0, endometrial glandular hyperplasia; N85.9 unspecified noninflammatory disorder of uterus; N87.9, dysplasia of cervix uteri; N92.4, excessive bleeding in the premenopausal period |
| EGCUT_2 | DEL | F | 40 | 0 | N72, inflammatory disease of cervix uteri; N87.1, moderate cervical dysplasia |
| EGCUT_3 | DEL | F | 35 | E28.2, polycystic ovarian syndrome; N91.3, primary oligomenorhhoea; N97.8, female infertility; Q51.3, bicornate uterus; Q52.1, doubling of vagina | A56.0, chlamydial infection of lower genitourinary tract; N76.1, subacute and chronic vaginitis; N83.2, unspecified ovarian cysts; O02.0, blighted ovum and nonhydatidiform mole; O10, pre-existing hypertension complicating pregnancy; O14.0, mild to moderate pre-eclampsia; O82.0, delivery by elective caesarean section; Z31.2, in vitro fertilization |
| EGCUT_4 | DEL | F | 50 | 0 | N30, cystitis |
| EGCUT_5 | DEL | F | 24 | 0 | N94.3, premenstrual tension syndrome |
| EGCUT_6 | DEL | F | 33 | N91.1, secondary amenorhhoea; N91.4, secondary oligomenorhhoea | N11, chronic tubulo-interstitial nephritis; N28.8, other specified disorders of kidney and ureter; N70.0, acute salpingitis and oophoritis; N71.0, acute inflammatory disease of uterus; N72, inflammatory disease of cervix uteri; N76, inflammation of vagina and vulva; N86, erosion and ectropion of cervix uteri; O16, unspecified maternal hypertension; O26.0, excessive weight gain in pregnancy; O28.8, other abnormal findings on antenatal screening of mother |
| EGCUT_7 | DUP | F | 48 | N80.8, endometriosis | N23, unspecified renal colic; N30, cystitis; N39, other disorders of urinary system; N81.6, rectocele; N83.2, unspecified ovarian cysts; O08.0, genital tract and pelvic infection following abortion, ectopic and molar pregnancy |
| EGCUT_8 | DUP | F | 34 | 0 | N10, acute tubulo-interstitial nephritis; N11, chronic tubulo-interstitial nephritis; N30, cystitis |

| | | | | | |
|---|---|---|---|---|---|
| EGCUT_9 | DUP | F | 30 | E28, ovarian dysfunction | N72, inflammatory disease of cervix uteri; O01.0, classical hydatidiform mole; O02.0, blighted ovum and nonhydatidiform mole; O03.4, spontaneous abortion: incomplete, without complication; O20.0, threatened abortion; O30.0, twin pregnancy; O34.2, maternal care due to uterine scar from previous surgery; O36.6, maternal care for excessive fetal growth; O47.1, false labour at or after 37 weeks of gestation; O66.2, obstructed labour due to unusually large fetus; O69.1, labour and delivery complicated by cord around neck, with compression; O70.0, first degree perineal laceration during delivery; O71.4, obstetric high vaginal laceration; O80.0, spontaneous vertex delivery; O99.6, diseases of the digestive system complicating pregnancy |
| EGCUT_10 | DUP | F | 61 | 0 | N03.9, chronic nephritis syndrome; N11.9, chronic tubulo-interstitial nephritis; N30, cystitis; N72, inflammatory disease of cervix uteri; N95.2, postmenopausal atrophic vaginitis |
| EGCUT_11 | DUP | F | 27 | N92, excessive, frequent and irregular menstruation | N30.0, acute cystitis; N61, inflammatory disorders of breast; N64.4, mastodynia; N70.9, salpingitis and oophoritis; N83.2, unspecified ovarian cysts |
| EGCUT_12 | DUP | F | 70 | 0 | A59.0, urogenital trichomoniasis; N30.0, acute cystitis; N70.1, chronic salpingitis and oophoritis; N95.2, postmenopausal atrophic vaginitis |
| EGCUT_13 | DUP | F | 20 | N91.2, unspecified amenorhhoea; Q51.1, doubling of uterus, cervix and vagina; | A63.8, other predominatly sexually transmitted diseases; N30, cystitis; N70.1, chronic salpingitis and oophoritis; N72, inflammatory disease of cervix uteri; N73.3, female acute pelvic peritonitis; N87.9, dysplasia of cervix uteri; O03, spontaneous abortion; O12.0, gestational oedema; O14.1, severe pre-eclampsia; O20.0, threatened abortion; O24.9, diabetes mellitus in pregnancy; O45.9, unspecified premature separation of placenta; O60, preterm labour and delivery; O82.1, delivery by emergency caesarean section; O90.8, other complications of the puerperium; O99.0, anaemia complicating pregnancy |
| EGCUT_14 | DUP | F | 23 | N91.3, primary oligomenorhhoea; Q50.5, embryonic cyst of broad ligament; Q62.5, duplication of ureter | A56.0, chlamydial infection of lower genitourinary tract; N30.2, chronic cystitis; N71.1, chronic inflammatory disease of uterus; N76.0, acute vaginitis; N83.2, unspecified ovarian cysts; N85.4, malposition of uterus; O47.1, false labour at or after 37 weeks of gestation; O71.4, obstetric high vaginal laceration; O71.7, obstetric haematoma of pelvis; O86.0, infection of obstetric surgical wound; O99.0, anaemia complicating pregnancy; O99.3, mental disorders and diseases of the nervous system complicating pregnancy |
| EGCUT_15 | DUP | F | 37 | E28.2, polycystic ovarian syndrome; N80, endometriosis; N92.1, excessive and frequent menstruation with irregular cycle; N97.1, female infertility | A56, chlamydial infection of lower genitourinary tract; N10, acute tubulo-interstitial nephritis; N11, chronic tubulo-interstitial nephritis; N30, cystitis; N70, salpingitis and oophoritis; N71, inflammatory disease of uterus; N73.6, female pelvic peritoneal adhesions; N76, inflammation of vagina and vulva; N93.8, other specified abnormal uterine and vaginal bleeding; N94, mittelschmerz; O82, single delivery by caesarean section; O99.0, anaemia complicating pregnancy |
| EGCUT_16 | DUP | F | 43 | E28.0, ovarian dysfunction: estrogen excess; N91.1, secondary amenorhhoea | A63.8, other predominatly sexually transmitted diseases; N76.0, acute vaginitis |
| EGCUT_17 | DUP | F | 42 | 0 | N39.9, unspecified disorder of urinary system; N76.1, subacute and chronic vaginitis; N83.0, follicular cyst of ovary; N94.5, secondary dysmenorrhoea |

**Supplementary Table S6. Enrichment analysis reports for gene lists differentially expressed in 16p11.2 human patients and mouse models**

Enrichment by Diseases (by Biomarkers). The top-50 most significant terms, both neoplastic and non-neoplastic, are represented.

**Mouse cortex (Blumenthal *et al*, Am J Hum Genet 2014); doi: 10.1016/j.ajhg.2014.05.004.**

| # | Diseases | Total | pValue | Min FDR | p-value | FDR | Network Objects In Data |
|---|---|---|---|---|---|---|---|
| 1 | Lung Neoplasms | 18496 | 1.015E-23 | 1.434E-20 | 1.015E-23 | 1.434E-20 | 914 |
| 2 | Thoracic Neoplasms | 18510 | 1.507E-23 | 1.434E-20 | 1.507E-23 | 1.434E-20 | 914 |
| 3 | Respiratory Tract Neoplasms | 18598 | 1.755E-22 | 1.113E-19 | 1.755E-22 | 1.113E-19 | 914 |
| 4 | Prostatic Neoplasms | 10758 | 1.135E-21 | 4.791E-19 | 1.135E-21 | 4.791E-19 | 606 |
| 5 | Prostatic Diseases | 10762 | 1.259E-21 | 4.791E-19 | 1.259E-21 | 4.791E-19 | 606 |
| 6 | Genital Neoplasms, Male | 10777 | 1.854E-21 | 5.882E-19 | 1.854E-21 | 5.882E-19 | 606 |
| 7 | Genital Diseases, Male | 10886 | 2.775E-21 | 7.543E-19 | 2.775E-21 | 7.543E-19 | 610 |
| 8 | Lung Diseases | 19035 | 4.124E-21 | 9.811E-19 | 4.124E-21 | 9.811E-19 | 925 |
| 9 | Respiratory Tract Diseases | 19170 | 7.086E-20 | 1.498E-17 | 7.086E-20 | 1.498E-17 | 926 |
| 10 | Urinary Bladder Neoplasms | 5147 | 4.622E-19 | 8.796E-17 | 4.622E-19 | 8.796E-17 | 338 |
| 11 | Urinary Bladder Diseases | 5173 | 5.543E-19 | 9.589E-17 | 5.543E-19 | 9.589E-17 | 339 |
| 12 | Uterine Diseases | 12673 | 5.194E-18 | 8.236E-16 | 5.194E-18 | 8.236E-16 | 672 |
| 13 | Uterine Neoplasms | 12666 | 7.694E-18 | 1.126E-15 | 7.694E-18 | 1.126E-15 | 671 |
| 14 | Urogenital Neoplasms | 18766 | 2.459E-17 | 3.283E-15 | 2.459E-17 | 3.283E-15 | 904 |
| 15 | Male Urogenital Diseases | 16294 | 2.588E-17 | 3.283E-15 | 2.588E-17 | 3.283E-15 | 813 |
| 16 | Endometrial Neoplasms | 11447 | 6.615E-17 | 7.868E-15 | 6.615E-17 | 7.868E-15 | 616 |
| 17 | Female Urogenital Diseases | 17493 | 8.805E-17 | 9.856E-15 | 8.805E-17 | 9.856E-15 | 856 |
| 18 | Neoplasms by Site | 22644 | 1.866E-16 | 1.972E-14 | 1.866E-16 | 1.972E-14 | 1026 |
| 19 | Female Urogenital Diseases and Pregnancy Complications | 17568 | 2.638E-16 | 2.642E-14 | 2.638E-16 | 2.642E-14 | 857 |
| 20 | Neoplasms | 23164 | 1.611E-15 | 1.532E-13 | 1.611E-15 | 1.532E-13 | 1038 |
| 21 | Genital Neoplasms, Female | 13760 | 2.408E-15 | 2.182E-13 | 2.408E-15 | 2.182E-13 | 705 |
| 22 | Urologic Neoplasms | 12487 | 3.426E-15 | 2.963E-13 | 3.426E-15 | 2.963E-13 | 652 |
| 23 | Rectal Neoplasms | 7089 | 4.064E-15 | 3.362E-13 | 4.064E-15 | 3.362E-13 | 415 |
| 24 | Genital Diseases, Female | 13893 | 8.181E-15 | 6.487E-13 | 8.181E-15 | 6.487E-13 | 708 |
| 25 | Urologic Diseases | 12869 | 2.301E-14 | 1.752E-12 | 2.301E-14 | 1.752E-12 | 664 |
| 26 | Colorectal Neoplasms | 10101 | 4.932E-14 | 3.610E-12 | 4.932E-14 | 3.610E-12 | 545 |
| 27 | Intestinal Neoplasms | 10114 | 6.452E-14 | 4.548E-12 | 6.452E-14 | 4.548E-12 | 545 |
| 28 | Rectal Diseases | 9766 | 9.510E-14 | 6.464E-12 | 9.510E-14 | 6.464E-12 | 529 |
| 29 | Uterine Cervical Neoplasms | 3917 | 5.972E-13 | 3.919E-11 | 5.972E-13 | 3.919E-11 | 253 |
| 30 | Uterine Cervical Diseases | 3920 | 6.500E-13 | 4.123E-11 | 6.500E-13 | 4.123E-11 | 253 |
| 31 | Gastrointestinal Neoplasms | 11479 | 2.144E-12 | 1.316E-10 | 2.144E-12 | 1.316E-10 | 596 |
| 32 | Intestinal Diseases | 10524 | 2.267E-12 | 1.348E-10 | 2.267E-12 | 1.348E-10 | 555 |
| 33 | Head and Neck Neoplasms | 7493 | 4.895E-12 | 2.823E-10 | 4.895E-12 | 2.823E-10 | 419 |
| 34 | Digestive System Diseases | 13247 | 5.166E-12 | 2.891E-10 | 5.166E-12 | 2.891E-10 | 668 |
| 35 | Digestive System Neoplasms | 12748 | 9.264E-12 | 5.037E-10 | 9.264E-12 | 5.037E-10 | 646 |
| 36 | Stomatognathic Diseases | 6090 | 1.364E-11 | 7.208E-10 | 1.364E-11 | 7.208E-10 | 352 |
| 37 | Gastrointestinal Diseases | 11755 | 1.655E-11 | 8.510E-10 | 1.655E-11 | 8.510E-10 | 603 |
| 38 | Breast Neoplasms | 9824 | 2.600E-11 | 1.302E-09 | 2.600E-11 | 1.302E-09 | 519 |
| 39 | Breast Diseases | 9826 | 2.698E-11 | 1.316E-09 | 2.698E-11 | 1.316E-09 | 519 |
| 40 | Skin Diseases | 12280 | 4.042E-11 | 1.923E-09 | 4.042E-11 | 1.923E-09 | 623 |
| 41 | Kidney Neoplasms | 10840 | 8.883E-11 | 4.123E-09 | 8.883E-11 | 4.123E-09 | 560 |
| 42 | Kidney Diseases | 11296 | 1.089E-10 | 4.932E-09 | 1.089E-10 | 4.932E-09 | 579 |
| 43 | Neoplasms by Histologic Type | 12700 | 1.174E-10 | 5.194E-09 | 1.174E-10 | 5.194E-09 | 638 |
| 44 | Mood Disorders | 1073 | 1.993E-10 | 8.618E-09 | 1.993E-10 | 8.618E-09 | 91 |
| 45 | Neuroectodermal Tumors, Primitive | 2944 | 2.217E-10 | 9.374E-09 | 2.217E-10 | 9.374E-09 | 193 |
| 46 | Behavior | 196 | 2.348E-10 | 9.715E-09 | 2.348E-10 | 9.715E-09 | 31 |
| 47 | Depressive Disorder, Major | 1054 | 3.977E-10 | 1.610E-08 | 3.977E-10 | 1.610E-08 | 89 |
| 48 | Self-Injurious Behavior | 156 | 4.299E-10 | 1.670E-08 | 4.299E-10 | 1.670E-08 | 27 |
| 49 | Suicide | 156 | 4.299E-10 | 1.670E-08 | 4.299E-10 | 1.670E-08 | 27 |
| 50 | Depressive Disorder | 1066 | 7.011E-10 | 2.668E-08 | 7.011E-10 | 2.668E-08 | 89 |

#### Human LCLs (Migliavacca *et al*, Am J Hum Genet 2015); doi: 10.1016/j.ajhg.2015.04.002.

| # | Diseases | Total | pValue | Min FDR | p-value | FDR | Network Objects In Data |
|---|---|---|---|---|---|---|---|
| 1 | Neoplasms by Site | 22644 | 3.614E-19 | 5.967E-16 | 3.614E-19 | 5.967E-16 | 1216 |
| 2 | Lung Neoplasms | 18496 | 1.486E-18 | 1.168E-15 | 1.486E-18 | 1.168E-15 | 1054 |
| 3 | Thoracic Neoplasms | 18510 | 2.182E-18 | 1.168E-15 | 2.182E-18 | 1.168E-15 | 1054 |
| 4 | Urologic Neoplasms | 12487 | 2.830E-18 | 1.168E-15 | 2.830E-18 | 1.168E-15 | 775 |
| 5 | Female Urogenital Diseases | 17493 | 1.159E-17 | 3.469E-15 | 1.159E-17 | 3.469E-15 | 1007 |
| 6 | Respiratory Tract Neoplasms | 18598 | 1.261E-17 | 3.469E-15 | 1.261E-17 | 3.469E-15 | 1055 |
| 7 | Female Urogenital Diseases and Pregnancy Complications | 17568 | 4.470E-17 | 1.054E-14 | 4.470E-17 | 1.054E-14 | 1008 |
| 8 | Lung Diseases | 19035 | 2.306E-16 | 4.759E-14 | 2.306E-16 | 4.759E-14 | 1069 |
| 9 | Urogenital Neoplasms | 18766 | 5.680E-16 | 1.042E-13 | 5.680E-16 | 1.042E-13 | 1056 |
| 10 | Urologic Diseases | 12869 | 9.495E-16 | 1.452E-13 | 9.495E-16 | 1.452E-13 | 782 |
| 11 | Neoplasms | 23164 | 9.671E-16 | 1.452E-13 | 9.671E-16 | 1.452E-13 | 1226 |
| 12 | Respiratory Tract Diseases | 19170 | 2.282E-15 | 3.139E-13 | 2.282E-15 | 3.139E-13 | 1071 |
| 13 | Kidney Neoplasms | 10840 | 3.141E-15 | 3.989E-13 | 3.141E-15 | 3.989E-13 | 678 |
| 14 | Male Urogenital Diseases | 16294 | 3.024E-14 | 3.566E-12 | 3.024E-14 | 3.566E-12 | 938 |
| 15 | Kidney Diseases | 11296 | 9.718E-13 | 1.070E-10 | 9.718E-13 | 1.070E-10 | 688 |
| 16 | Genital Neoplasms, Female | 13760 | 3.576E-12 | 3.690E-10 | 3.576E-12 | 3.690E-10 | 807 |
| 17 | Uterine Neoplasms | 12666 | 1.362E-11 | 1.323E-09 | 1.362E-11 | 1.323E-09 | 750 |
| 18 | Uterine Diseases | 12673 | 1.563E-11 | 1.433E-09 | 1.563E-11 | 1.433E-09 | 750 |
| 19 | Endometrial Neoplasms | 11447 | 2.096E-11 | 1.821E-09 | 2.096E-11 | 1.821E-09 | 688 |
| 20 | Genital Diseases, Female | 13893 | 2.207E-11 | 1.822E-09 | 2.207E-11 | 1.822E-09 | 809 |
| 21 | Urinary Bladder Neoplasms | 5147 | 1.615E-08 | 1.270E-06 | 1.615E-08 | 1.270E-06 | 338 |
| 22 | Urinary Bladder Diseases | 5173 | 1.910E-08 | 1.433E-06 | 1.910E-08 | 1.433E-06 | 339 |
| 23 | Rectal Neoplasms | 7089 | 1.068E-07 | 7.668E-06 | 1.068E-07 | 7.668E-06 | 438 |
| 24 | Digestive System Diseases | 13247 | 1.025E-06 | 7.052E-05 | 1.025E-06 | 7.052E-05 | 746 |
| 25 | Medulloblastoma | 2554 | 1.282E-06 | 8.464E-05 | 1.282E-06 | 8.464E-05 | 180 |
| 26 | Digestive System Neoplasms | 12748 | 3.364E-06 | 2.136E-04 | 3.364E-06 | 2.136E-04 | 717 |
| 27 | Breast Diseases | 9826 | 4.917E-06 | 3.007E-04 | 4.917E-06 | 3.007E-04 | 568 |
| 28 | Breast Neoplasms | 9824 | 6.260E-06 | 3.573E-04 | 6.260E-06 | 3.573E-04 | 567 |
| 29 | Papilloma, Intraductal | 4 | 6.276E-06 | 3.573E-04 | 6.276E-06 | 3.573E-04 | 4 |
| 30 | Neuroectodermal Tumors, Primitive | 2944 | 7.766E-06 | 4.274E-04 | 7.766E-06 | 4.274E-04 | 198 |
| 31 | Uterine Cervical Neoplasms | 3917 | 9.863E-06 | 5.253E-04 | 9.863E-06 | 5.253E-04 | 252 |
| 32 | Uterine Cervical Diseases | 3920 | 1.043E-05 | 5.380E-04 | 1.043E-05 | 5.380E-04 | 252 |
| 33 | Glioma | 5034 | 1.461E-05 | 7.308E-04 | 1.461E-05 | 7.308E-04 | 312 |
| 34 | Neoplasms, Hormone-Dependent | 5 | 3.013E-05 | 1.463E-03 | 3.013E-05 | 1.463E-03 | 4 |
| 35 | Virus Diseases | 1667 | 3.614E-05 | 1.705E-03 | 3.614E-05 | 1.705E-03 | 120 |
| 36 | RNA Virus Infections | 1159 | 4.096E-05 | 1.878E-03 | 4.096E-05 | 1.878E-03 | 89 |
| 37 | Neoplasms, Neuroepithelial | 5288 | 4.327E-05 | 1.931E-03 | 4.327E-05 | 1.931E-03 | 322 |
| 38 | Retroviridae Infections | 860 | 4.711E-05 | 2.047E-03 | 4.711E-05 | 2.047E-03 | 70 |
| 39 | Immunologic Deficiency Syndromes | 1007 | 5.480E-05 | 2.247E-03 | 5.480E-05 | 2.247E-03 | 79 |
| 40 | Sexually Transmitted Diseases | 849 | 5.581E-05 | 2.247E-03 | 5.581E-05 | 2.247E-03 | 69 |
| 41 | Sexually Transmitted Diseases, Viral | 849 | 5.581E-05 | 2.247E-03 | 5.581E-05 | 2.247E-03 | 69 |
| 42 | HIV Infections | 851 | 5.997E-05 | 2.297E-03 | 5.997E-05 | 2.297E-03 | 69 |
| 43 | Lentivirus Infections | 851 | 5.997E-05 | 2.297E-03 | 5.997E-05 | 2.297E-03 | 69 |
| 44 | Gastrointestinal Neoplasms | 11479 | 6.122E-05 | 2.297E-03 | 6.122E-05 | 2.297E-03 | 642 |
| 45 | Prostatic Neoplasms | 10758 | 6.966E-05 | 2.556E-03 | 6.966E-05 | 2.556E-03 | 605 |
| 46 | Prostatic Diseases | 10762 | 7.305E-05 | 2.622E-03 | 7.305E-05 | 2.622E-03 | 605 |
| 47 | Endocarditis | 6 | 8.678E-05 | 2.999E-03 | 8.678E-05 | 2.999E-03 | 4 |
| 48 | Genital Neoplasms, Male | 10777 | 8.720E-05 | 2.999E-03 | 8.720E-05 | 2.999E-03 | 605 |
| 49 | Gastrointestinal Diseases | 11755 | 9.516E-05 | 3.206E-03 | 9.516E-05 | 3.206E-03 | 654 |
| 50 | Smooth Muscle Tumor | 11 | 1.124E-04 | 3.710E-03 | 1.124E-04 | 3.710E-03 | 5 |

#### Human LCLs (Blumenthal *et al*, Am J Hum Genet 2014); doi: 10.1016/j.ajhg.2014.05.004.

| # | Diseases | Total | pValue | Min FDR | p-value | FDR | Network Objects In Data |
|---|---|---|---|---|---|---|---|
| 1 | Respiratory Tract Neoplasms | 18598 | 2.627E-12 | 1.859E-09 | 2.627E-12 | 1.859E-09 | 480 |
| 2 | Lung Neoplasms | 18496 | 2.961E-12 | 1.859E-09 | 2.961E-12 | 1.859E-09 | 478 |
| 3 | Thoracic Neoplasms | 18510 | 3.612E-12 | 1.859E-09 | 3.612E-12 | 1.859E-09 | 478 |
| 4 | Lung Diseases | 19035 | 1.141E-10 | 4.403E-08 | 1.141E-10 | 4.403E-08 | 483 |
| 5 | Respiratory Tract Diseases | 19170 | 1.588E-10 | 4.905E-08 | 1.588E-10 | 4.905E-08 | 485 |
| 6 | Uterine Neoplasms | 12666 | 7.424E-10 | 1.778E-07 | 7.424E-10 | 1.778E-07 | 352 |
| 7 | Uterine Diseases | 12673 | 8.059E-10 | 1.778E-07 | 8.059E-10 | 1.778E-07 | 352 |
| 8 | Stomatognathic Diseases | 6090 | 3.619E-09 | 6.984E-07 | 3.619E-09 | 6.984E-07 | 196 |
| 9 | Endometrial Neoplasms | 11447 | 1.081E-08 | 1.717E-06 | 1.081E-08 | 1.717E-06 | 320 |
| 10 | Neoplasms by Site | 22644 | 1.112E-08 | 1.717E-06 | 1.112E-08 | 1.717E-06 | 538 |
| 11 | Skin Diseases | 12280 | 1.332E-08 | 1.734E-06 | 1.332E-08 | 1.734E-06 | 338 |
| 12 | Genital Neoplasms, Female | 13760 | 1.348E-08 | 1.734E-06 | 1.348E-08 | 1.734E-06 | 370 |
| 13 | Mouth Neoplasms | 4328 | 1.777E-08 | 2.111E-06 | 1.777E-08 | 2.111E-06 | 148 |
| 14 | Breast Neoplasms | 9824 | 2.013E-08 | 2.118E-06 | 2.013E-08 | 2.118E-06 | 282 |
| 15 | Breast Diseases | 9826 | 2.058E-08 | 2.118E-06 | 2.058E-08 | 2.118E-06 | 282 |
| 16 | Skin and Connective Tissue Diseases | 12945 | 2.796E-08 | 2.699E-06 | 2.796E-08 | 2.699E-06 | 351 |
| 17 | Genital Diseases, Female | 13893 | 3.377E-08 | 3.067E-06 | 3.377E-08 | 3.067E-06 | 371 |
| 18 | Mouth Diseases | 4970 | 4.518E-08 | 3.875E-06 | 4.518E-08 | 3.875E-06 | 163 |
| 19 | Urogenital Neoplasms | 18766 | 4.794E-08 | 3.896E-06 | 4.794E-08 | 3.896E-06 | 469 |
| 20 | Male Urogenital Diseases | 16294 | 9.413E-08 | 7.267E-06 | 9.413E-08 | 7.267E-06 | 419 |
| 21 | Rectal Neoplasms | 7089 | 1.086E-07 | 7.987E-06 | 1.086E-07 | 7.987E-06 | 214 |
| 22 | Head and Neck Neoplasms | 7493 | 2.470E-07 | 1.734E-05 | 2.470E-07 | 1.734E-05 | 222 |
| 23 | Neoplasms | 23164 | 5.822E-07 | 3.908E-05 | 5.822E-07 | 3.908E-05 | 542 |
| 24 | Genital Neoplasms, Male | 10777 | 6.855E-07 | 4.410E-05 | 6.855E-07 | 4.410E-05 | 296 |
| 25 | Genital Diseases, Male | 10886 | 8.342E-07 | 5.152E-05 | 8.342E-07 | 5.152E-05 | 298 |
| 26 | Laryngeal Diseases | 2430 | 9.001E-07 | 5.203E-05 | 9.001E-07 | 5.203E-05 | 90 |
| 27 | Prostatic Diseases | 10762 | 9.098E-07 | 5.203E-05 | 9.098E-07 | 5.203E-05 | 295 |
| 28 | Prostatic Neoplasms | 10758 | 1.336E-06 | 7.369E-05 | 1.336E-06 | 7.369E-05 | 294 |
| 29 | Colorectal Neoplasms | 10101 | 1.388E-06 | 7.387E-05 | 1.388E-06 | 7.387E-05 | 279 |
| 30 | Intestinal Neoplasms | 10114 | 1.571E-06 | 8.085E-05 | 1.571E-06 | 8.085E-05 | 279 |
| 31 | Urinary Bladder Neoplasms | 5147 | 2.016E-06 | 1.004E-04 | 2.016E-06 | 1.004E-04 | 160 |
| 32 | Digestive System Neoplasms | 12748 | 2.234E-06 | 1.078E-04 | 2.234E-06 | 1.078E-04 | 337 |
| 33 | Rectal Diseases | 9766 | 2.419E-06 | 1.132E-04 | 2.419E-06 | 1.132E-04 | 270 |
| 34 | Urinary Bladder Diseases | 5173 | 2.736E-06 | 1.243E-04 | 2.736E-06 | 1.243E-04 | 160 |
| 35 | Pulmonary Eosinophilia | 6 | 3.485E-06 | 1.537E-04 | 3.485E-06 | 1.537E-04 | 4 |
| 36 | Urologic Neoplasms | 12487 | 3.762E-06 | 1.614E-04 | 3.762E-06 | 1.614E-04 | 330 |
| 37 | Laryngeal Neoplasms | 2412 | 4.092E-06 | 1.708E-04 | 4.092E-06 | 1.708E-04 | 87 |
| 38 | Female Urogenital Diseases and Pregnancy Complications | 17568 | 4.267E-06 | 1.734E-04 | 4.267E-06 | 1.734E-04 | 437 |
| 39 | Female Urogenital Diseases | 17493 | 5.335E-06 | 2.112E-04 | 5.335E-06 | 2.112E-04 | 435 |
| 40 | Skin Neoplasms | 4797 | 5.635E-06 | 2.175E-04 | 5.635E-06 | 2.175E-04 | 149 |
| 41 | Intestinal Diseases | 10524 | 6.030E-06 | 2.271E-04 | 6.030E-06 | 2.271E-04 | 285 |
| 42 | Hypopharyngeal Neoplasms | 1187 | 1.212E-05 | 4.432E-04 | 1.212E-05 | 4.432E-04 | 50 |
| 43 | Lymphoproliferative Disorders | 4193 | 1.234E-05 | 4.432E-04 | 1.234E-05 | 4.432E-04 | 132 |
| 44 | Urologic Diseases | 12869 | 1.420E-05 | 4.906E-04 | 1.420E-05 | 4.906E-04 | 335 |
| 45 | Immunoproliferative Disorders | 4127 | 1.430E-05 | 4.906E-04 | 1.430E-05 | 4.906E-04 | 130 |
| 46 | Carcinoma, Adenoid Cystic | 37 | 1.482E-05 | 4.974E-04 | 1.482E-05 | 4.974E-04 | 7 |
| 47 | Digestive System Diseases | 13247 | 1.567E-05 | 5.147E-04 | 1.567E-05 | 5.147E-04 | 343 |
| 48 | Pharyngeal Neoplasms | 1930 | 1.871E-05 | 6.019E-04 | 1.871E-05 | 6.019E-04 | 71 |
| 49 | Lymphatic Diseases | 2997 | 1.941E-05 | 6.116E-04 | 1.941E-05 | 6.116E-04 | 100 |
| 50 | Pharyngeal Diseases | 1941 | 2.251E-05 | 6.952E-04 | 2.251E-05 | 6.952E-04 | 71 |

#### Supplementary Table S7. Human MRI cohort data

| Anatomical location | Coordinates [mm] | | | Statistical results | |
|---|---|---|---|---|---|
| | *x* | *y* | *z* | *p-value* | *Z-score* |
| Hypothalamus | 2 | 2 | -8 | <.001 | 3.49 |
| Caudate left | -14 | -8 | 21 | 0.001 | 4.27 |
| Caudate right | 17 | 3 | 26 | 0.001 | 4.37 |

[1]Coordinates according to Montreal Neurological Institute reference space with p-values after family-wise error correction (following small-volume correction procedure) and corresponding Z-scores for the obtained findings (see Worsley *et al*, *Hum Brain Mapp* 1996).

#### Supplementary Table S8. Description of mouse MRI data and analyses

##### Relative volume of the hypothalamus

| Cohort | $16p11.2^{Del/+}$ | WT | $16p11.2^{Dup/+}$ |
|---|---|---|---|
| Animal_1 | 2.451 | 2.429 | 2.251 |
| Animal_2 | 2.512 | 2.399 | 2.346 |
| Animal_3 | 2.523 | 2.415 | 2.303 |
| Animal_4 | 2.561 | 2.342 | 2.324 |
| Animal_5 | 2.498 | 2.364 | 2.298 |
| Animal_6 | 2.512 | 2.395 | 2.246 |
| Animal_7 | 2.528 | 2.438 | 2.333 |
| Animal_8 | 2.505 | 2.379 | 2.325 |
| Animal_9 | 2.483 | 2.438 | 2.286 |
| Animal_10 | 2.475 | 2.391 | NA |
| Animal_11 | NA | 2.385 | NA |
| Animal_12 | NA | 2.377 | NA |
| Mean | 2.505 | 2.396 | 2.301 |
| SD | 0.031 | 0.030 | 0.035 |

| Tukey's multiple comparisons test | Mean Diff. | 95.00% CI of diff. | Adjusted P-value |
|---|---|---|---|
| DEL vs WT | 0.1106 | 0.076 to 0.145 | <0.0001 |
| DEL vs DUP | 0.2035 | 0.167 to 0.240 | <0.0001 |
| WT vs DUP | 0.09292 | 0.057 to 0.129 | <0.0001 |

| Test contrast | Mean 1 | Mean 2 | Mean Diff. | SE of diff. | N1 | N2 | q | DF |
|---|---|---|---|---|---|---|---|---|
| DEL vs WT | 2.505 | 2.394 | 0.1106 | 0.01401 | 10 | 11 | 11.17 | 27 |
| DEL vs DUP | 2.505 | 2.301 | 0.2035 | 0.01473 | 10 | 9 | 19.54 | 27 |
| WT vs DUP | 2.394 | 2.301 | 0.09292 | 0.01441 | 11 | 9 | 9.122 | 27 |

| ANOVA test summary | F | P value | R square |
|---|---|---|---|
| | 96.19 | <0.0001 | 0.8769 |

| ANOVA table | SS | DF | MS | F (DFn, DFd) | P value |
|---|---|---|---|---|---|
| Treatment (between columns) | 0.1976 | 2 | 0.09882 | F (2, 27) = 96.19 | P<0.0001 |
| Residual (within columns) | 0.02774 | 27 | 0.001027 | | |
| Total | 0.2254 | 29 | | | |

#### Absolute volume of the hypothalamus

| Cohort | $16p11.2^{Del/+}$ | WT | $16p11.2^{Dup/+}$ |
|---|---|---|---|
| Animal_1 | 10.52174 | 11.219155 | 10.37925 |
| Animal_2 | 10.943981 | 11.106334 | 10.37361 |
| Animal_3 | 10.954544 | 11.426926 | 10.968065 |
| Animal_4 | 11.162942 | 10.941198 | 10.568838 |
| Animal_5 | 10.722183 | 11.175008 | 11.035989 |
| Animal_6 | 10.811293 | 11.290945 | 10.297445 |
| Animal_7 | 10.952579 | 11.681481 | 10.60832 |
| Animal_8 | 10.088052 | 10.594931 | 10.196688 |
| Animal_9 | 10.791144 | 11.185131 | 10.227525 |
| Animal_10 | 9.890261 | 11.098188 | NA |
| Animal_11 | NA | 11.045756 | NA |
| Animal_12 | NA | 10.942418 | NA |
| Mean | 10.6838719 | 11.14228925 | 10.51730333 |
| SD | 0.405921358 | 0.26887007 | 0.307808913 |

| Tukey's multiple comparisons test | Mean Diff. | 95.00% CI of diff. | Adjusted P-value |
|---|---|---|---|
| DEL vs WT | -38.56 | -51.36 to -25.76 | <0.0001 |
| DEL vs DUP | -30.61 | -44.35 to -16.87 | <0.0001 |
| WT vs DUP | 7.951 | -5.233 to 21.14 | 0.3099 |

| Test contrast | Mean 1 | Mean 2 | Mean Diff. | SE of diff. | N1 | N2 | q | DF |
|---|---|---|---|---|---|---|---|---|
| DEL vs WT | 426.5 | 465 | -38.56 | 5.174 | 10 | 12 | 10.54 | 28 |
| DEL vs DUP | 426.5 | 457.1 | -30.61 | 5.552 | 10 | 9 | 7.797 | 28 |
| WT vs DUP | 465 | 457.1 | 7.951 | 5.328 | 12 | 9 | 2.11 | 28 |

| ANOVA test summary | F | p-value | R square |
|---|---|---|---|
| | 29.78 | <0.0001 | 0.6802 |

| ANOVA table | SS | DF | MS | F (DFn, DFd) | p-value |
|---|---|---|---|---|---|
| Treatment (between columns) | 8696 | 2 | 4348 | F (2, 28) = 29.78 | P<0.0001 |
| Residual (within columns) | 4088 | 28 | 146 | | |
| Total | 12784 | 30 | | | |

### Supplementary Table S9. Mendelian randomization test and data summary

##### MR summary of main results

| Probe_ID | Gene | Univariate analysis | | | | Multivariate analysis | | |
|---|---|---|---|---|---|---|---|---|
| | | Beta_SMR | SE_SMR | P_SMR | P_HEIDI | Beta_MMR | SE_MMR | P_MMR |
| ILMN_1724406 | INO80E | 0.0978 | 0.0152 | 1.3e-10 | 1.3e-02 | 0.07114 | 0.0182 | 9.3e-5 |
| ILMN_1786843 | KCTD13 | -0.1537 | 0.0335 | 4.5e-06 | 1.1e-02 | -0.07443 | 0.0226 | 9.7e-4 |
| ILMN_1791147 | YPEL3 | -0.2295 | 0.0544 | 2.4e-05 | 3.0e-03 | 0.07102 | 0.0246 | 3.8e-3 |
| ILMN_1700268 | QPRT | -0.0268 | 0.0133 | 4.5e-02 | 5.6e-01 | 0.02047 | 0.0090 | 2.4e-2 |
| ILMN_1751061 | C16orf54 | -0.0107 | 0.0047 | 2.4e-02 | 2.8e-11 | -0.00808 | 0.0049 | 9.8e-2 |
| ILMN_1750364 | PPP4C | 0.0507 | 0.0090 | 1.6e-08 | 7.3e-04 | 0.00696 | 0.0089 | 4.3e-1 |
| ILMN_1803277 | MVP | -0.1417 | 0.0325 | 1.3e-05 | 7.7e-06 | -0.01309 | 0.0184 | 4.8e-1 |
| ILMN_1801040 | SPN | 0.0303 | 0.0194 | 1.2e-01 | 2.4e-01 | -0.01094 | 0.0158 | 4.9e-1 |
| ILMN_2121774 | ZG16 | 0.0181 | 0.0211 | 3.9e-01 | 4.1e-01 | -0.02012 | 0.0299 | 5.0e-1 |
| ILMN_2194828 | PAGR1 | -0.0042 | 0.0342 | 9.0e-01 | 1.4e-03 | -0.01024 | 0.0254 | 6.9e-1 |
| ILMN_1713749 | CORO1A | 0.0940 | 0.0207 | 5.4e-06 | 2.3e-04 | -0.00367 | 0.0166 | 8.2e-1 |
| ILMN_2402341 | MAPK3 | 0.0322 | 0.0053 | 9.8e-10 | 1.1e-02 | -0.00016 | 0.0070 | 9.8e-1 |

Significance levels: P_SMR < 0.05/12; P_HEIDI > 0.009

##### Full results of MR analysis with the SMR tool

| Chr | Probe_ID | Probe_bp | Gene | SNP | SNP_bp | A1 | A2 | A1_freq_UK10K | Beta_gwas | SE_gwas | P_gwas | Beta_eQTL | SE_eQTL | Z_eQTL | Beta_SMR | SE_SMR | P_SMR | P_HEIDI | Std_Beta_SMR | Std_SE_SMR |
|---|---|---|---|---|---|---|---|---|---|---|---|---|---|---|---|---|---|---|---|---|
| 16 | ILMN_1751061 | 29754963 | C16orf54 | rs75422760 | 29748442 | T | C | 0.1747 | -0.0160 | 0.0071 | 2.4e-2 | 0.785 | 0.015 | 52.81 | -0.0204 | 0.0090 | 2.4e-02 | 2.8e-11 | -0.0107 | 0.0047 |
| 16 | ILMN_1681032 | 29754963 | C16orf54 | rs7203712 | 29716569 | T | G | 0.1989 | -0.0110 | 0.0061 | 7.3e-2 | 0.434 | 0.015 | 29.62 | -0.0253 | 0.0141 | 7.3e-02 | 7.0e-10 | -0.0146 | 0.0081 |
| 16 | ILMN_1713749 | 30199790 | CORO1A | rs7199462 | 30158757 | T | G | 0.4985 | -0.0210 | 0.0043 | 1.0e-6 | -0.147 | 0.012 | -12.40 | 0.1432 | 0.0315 | 5.4e-06 | 2.3e-04 | 0.0940 | 0.0207 |
| 16 | ILMN_1724406 | 30016716 | INO80E | rs4788204 | 29995218 | A | G | 0.4564 | -0.0300 | 0.0043 | 3.8e-12 | -0.201 | 0.012 | -17.01 | 0.1493 | 0.0232 | 1.3e-10 | 1.3e-02 | 0.0978 | 0.0152 |
| 16 | ILMN_1786843 | 29917858 | KCTD13 | rs4548895 | 29923510 | G | A | 0.3815 | 0.0240 | 0.0044 | 4.2e-8 | -0.103 | 0.012 | -8.38 | -0.2320 | 0.0506 | 4.5e-06 | 1.1e-02 | -0.1537 | 0.0335 |
| 16 | ILMN_2402341 | 30125607 | MAPK3 | rs55732507 | 30141985 | C | T | 0.4035 | 0.0280 | 0.0045 | 6.7e-10 | 0.550 | 0.012 | 44.03 | 0.0509 | 0.0083 | 9.8e-10 | 1.1e-02 | 0.0322 | 0.0053 |
| 16 | ILMN_1812747 | 30125540 | MAPK3 | rs28529403 | 30134656 | C | T | 0.4017 | 0.0280 | 0.0045 | 6.7e-10 | 0.687 | 0.087 | 7.93 | 0.0407 | 0.0084 | 1.1e-06 | 3.8e-01 | 0.0258 | 0.0053 |
| 16 | ILMN_1667260 | 30125472 | MAPK3 | rs55732507 | 30141985 | C | T | 0.4035 | 0.0280 | 0.0045 | 6.7e-10 | 0.495 | 0.013 | 39.06 | 0.0565 | 0.0093 | 1.1e-09 | 1.5e-02 | 0.0358 | 0.0059 |
| 16 | ILMN_2344373 | 29857568 | MVP | rs34504737 | 29879215 | T | C | 0.0058 | -0.0270 | 0.0295 | 3.6e-1 | -0.659 | 0.116 | -5.67 | 0.0410 | 0.0453 | 3.7e-01 | 8.5e-04 | 0.0257 | 0.0285 |
| 16 | ILMN_1803277 | 29859121 | MVP | rs71389428 | 29842632 | A | G | 0.1359 | -0.0350 | 0.0067 | 1.7e-7 | 0.151 | 0.019 | 7.93 | -0.2310 | 0.0529 | 1.3e-05 | 7.7e-06 | -0.1417 | 0.0325 |
| 16 | ILMN_2194828 | 29831201 | PAGR1 | rs12933548 | 29790709 | T | C | 0.1108 | -0.0010 | 0.0080 | 9.0e-1 | 0.131 | 0.019 | 6.79 | -0.0076 | 0.0612 | 9.0e-01 | 1.4e-03 | -0.0042 | 0.0342 |
| 16 | ILMN_1750364 | 30096501 | PPP4C | rs66467443 | 30050494 | A | G | 0.4712 | -0.0250 | 0.0043 | 5.3e-9 | -0.325 | 0.014 | -22.79 | 0.0770 | 0.0136 | 1.6e-08 | 7.3e-04 | 0.0507 | 0.0090 |
| 16 | ILMN_1700268 | 29708897 | QPRT | rs11574552 | 29674033 | C | T | 0.0739 | 0.0170 | 0.0084 | 4.3e-2 | -0.406 | 0.023 | -17.70 | -0.0419 | 0.0209 | 4.5e-02 | 5.6e-01 | -0.0268 | 0.0133 |
| 16 | ILMN_1801040 | 29681405 | SPN | rs12928754 | 29679989 | A | G | 0.3779 | -0.0076 | 0.0048 | 1.1e-1 | -0.151 | 0.012 | -12.22 | 0.0502 | 0.0320 | 1.2e-01 | 2.4e-01 | 0.0303 | 0.0194 |
| 16 | ILMN_1791147 | 30103799 | YPEL3 | rs4788212 | 30034469 | G | T | 0.4717 | -0.0250 | 0.0043 | 4.3e-9 | 0.072 | 0.012 | 6.07 | -0.3464 | 0.0821 | 2.4e-05 | 3.0e-03 | -0.2295 | 0.0544 |
| 16 | ILMN_2121774 | 29791795 | ZG16 | rs188630014 | 28909924 | A | G | 0.0036 | -0.0360 | 0.0411 | 3.8e-1 | -1.141 | 0.245 | -4.66 | 0.0316 | 0.0366 | 3.9e-01 | 4.1e-01 | 0.0181 | 0.0211 |

##### Full results of multivariate MR analysis with different probe and tool combinations

| Probe_ID | Gene | McDaid et al. (eQTL threshold 5e-08) | | | Burgess et al. (eQTL threshold 5e-08) | | | McDaid et al. (eQTL threshold 1e-05) | | |
|---|---|---|---|---|---|---|---|---|---|---|
| | | Beta | SE | P | Beta | SE | P | Beta | SE | P |
| ILMN_1751061 | C16orf54 | -0.0087 | 0.0058 | 1.4e-1 | -0.0102 | 0.0091 | 2.6e-1 | -0.00808 | 0.0049 | 9.8e-2 |
| ILMN_1713749 | CORO1A | 0.0327 | 0.0226 | 1.5e-1 | 0.0262 | 0.0265 | 3.2e-1 | -0.00367 | 0.0166 | 8.2e-1 |
| ILMN_1724406 | INO80E | 0.0781 | 0.0215 | 2.8e-4 | 0.0719 | 0.0294 | 1.5e-2 | 0.07114 | 0.0182 | 9.3e-5 |
| ILMN_1786843 | KCTD13 | -0.0678 | 0.0274 | 1.3e-2 | -0.0832 | 0.0373 | 2.6e-2 | -0.07443 | 0.0226 | 9.7e-4 |
| ILMN_2402341 | MAPK3 | -0.0059 | 0.0090 | 5.1e-1 | -0.0052 | 0.0108 | 6.3e-1 | -0.00016 | 0.0070 | 9.8e-1 |
| ILMN_1803277 | MVP | -0.0408 | 0.0227 | 7.2e-2 | -0.0588 | 0.0257 | 2.2e-2 | -0.01309 | 0.0184 | 4.8e-1 |
| ILMN_2194828 | PAGR1 | -0.0616 | 0.0341 | 7.1e-2 | -0.0824 | 0.0384 | 3.2e-2 | -0.01024 | 0.0254 | 6.9e-1 |
| ILMN_1750364 | PPP4C | 0.0044 | 0.0117 | 7.1e-1 | -0.0103 | 0.0168 | 5.4e-1 | 0.00696 | 0.0089 | 4.3e-1 |
| ILMN_1700268 | QPRT | 0.0112 | 0.0127 | 3.8e-1 | 0.0076 | 0.0168 | 6.5e-1 | 0.02047 | 0.0090 | 2.4e-2 |
| ILMN_1801040 | SPN | -0.0084 | 0.0170 | 6.2e-1 | 0.0024 | 0.0197 | 9.1e-1 | -0.01094 | 0.0158 | 4.9e-1 |
| ILMN_1791147 | YPEL3 | 0.0558 | 0.0278 | 4.5e-2 | 0.0547 | 0.0330 | 9.8e-2 | 0.07102 | 0.0246 | 3.8e-3 |
| ILMN_2121774 | ZG16 | | | | | | | -0.02012 | 0.0299 | 5.0e-1 |

Summary statistics of multivariate MR analysis are availabe upon request.

### Supplementary Table S10. GnRH3 overexpression and downregulation

#### 16p11.2 BP4-BP5 single-gene overexpression experiments

| Injection | Measurement | | | | Adjusted p-value (Tukey post-hoc) |
|---|---|---|---|---|---|
| | N | Mean | SD | SEM | vs Control |
| Controls | 1807 | 100.000 | 30.966 | 0.728 | - |
| 50pg *BOLA2B* RNA | 201 | 92.474 | 24.457 | 1.725 | 0.2974 |
| 50pg *SLX1B* RNA | 203 | 92.427 | 37.236 | 2.613 | 0.2753 |
| 50pg *SPN* RNA | 135 | 93.760 | 26.406 | 2.273 | 0.9519 |
| 50pg *QPRT* RNA | 120 | 96.623 | 32.303 | 2.949 | >0.9999 |
| 50pg *C16orf54* RNA | 133 | 96.297 | 28.718 | 2.490 | >0.9999 |
| 50pg *ZG16* RNA | 188 | 94.964 | 28.218 | 2.058 | 0.9782 |
| 12.5pg *KIF22* RNA | 106 | 96.643 | 41.247 | 4.006 | >0.9999 |
| 50pg *MAZ* RNA | 239 | 103.763 | 38.023 | 2.460 | 0.9988 |
| 50pg *PRRT2* RNA | 169 | 98.396 | 34.587 | 2.661 | >0.9999 |
| 50pg *C16orf53* RNA | 153 | 99.215 | 24.682 | 1.995 | >0.9999 |
| 50pg *MVP* RNA | 183 | 98.592 | 33.525 | 2.478 | >0.9999 |
| 50pg *CDIPT* RNA | 154 | 99.714 | 34.140 | 2.751 | >0.9999 |
| 50pg SEZ6L2 RNA | 93 | 104.605 | 30.609 | 3.174 | >0.9999 |
| 50pg *ASPHD1* RNA | 174 | 81.307 | 29.932 | 2.269 | <0.0001 |
| 50pg *KCTD13* RNA | 180 | 96.073 | 31.616 | 2.356 | 0.9997 |
| 50pg *TMEM219* RNA | 163 | 100.661 | 32.575 | 2.552 | >0.9999 |
| 50pg *TAOK2* RNA | 170 | 99.585 | 41.096 | 3.152 | >0.9999 |
| 50pg *HIRIP3* RNA | 194 | 93.947 | 28.131 | 2.020 | 0.8084 |
| 50pg *INO80E* RNA | 177 | 90.899 | 32.6120 | 2.452 | 0.0889 |
| 50pg *DOC2A* RNA | 78 | 90.706 | 36.905 | 4.179 | 0.8032 |
| 50pg *C16orf92* RNA | 180 | 95.618 | 28.061 | 2.092 | 0.9981 |
| 50pg *FAM57B* RNA | 139 | 102.349 | 36.594 | 3.104 | >0.9999 |
| 50pg *ALDOA* RNA | 192 | 100.951 | 33.176 | 2.394 | >0.9999 |
| 12.5pg *PPP4C* RNA | 160 | 107.543 | 29.986 | 2.371 | 0.5331 |
| 50pg *TBX6* RNA | 123 | 99.218 | 34.747 | 3.133 | >0.9999 |
| 50pg *YPEL3* RNA | 140 | 99.085 | 33.946 | 2.869 | >0.9999 |
| 50pg *GDPD3* RNA | 210 | 96.660 | 32.070 | 2.213 | >0.9999 |
| 50pg *MAPK3* RNA | 146 | 100.624 | 31.294 | 2.590 | >0.9999 |
| 50pg *CORO1A* RNA | 157 | 95.304 | 26.845 | 2.143 | 0.9979 |
| 50pg *SULT1A3* RNA | 265 | 100.446 | 33.932 | 2.084 | >0.9999 |

#### *ASPHD1* gene editing

| Injection | Measurement | | | | Adjusted p-value (Tukey post-hoc) | |
|---|---|---|---|---|---|---|
| | N | Mean | SD | SEM | vs Control | vs gRNA alone |
| Controls | 153 | 100.000 | 34.108 | 2.758 | - | - |
| 100pg gRNA alone | 111 | 100.597 | 32.981 | 3.130 | 0.9893 | - |
| 100pg gRNA + 200pg Cas9 | 138 | 86.718 | 35.645 | 3.034 | 0.0031 | 0.0047 |

### Supplementary Table S11. *ASPHD1* epistasis

| Injection | Measurement | | | | Adjusted p-value (Tukey post-hoc) | |
|---|---|---|---|---|---|---|
| | N | Mean | SD | SEM | vs Control | vs. *ASPHD1* RNA |
| Controls | 170 | 100.000 | 24.160 | 1.853 | - | - |
| 25pg *ASPHD1* RNA | 108 | 90.784 | 22.589 | 2.174 | 0.0519 | - |
| 50pg *KCTD13* RNA | 67 | 96.635 | 30.660 | 3.745 | 0.9931 | 0.8541 |
| 25pg *ASPHD1* RNA + 50pg *KCTD13* RNA | 70 | 76.307 | 23.412 | 2.798 | <0.0001 | 0.003 |
| 50pg *INO80E* RNA | 53 | 97.924 | 22.262 | 3.058 | >0.9999 | 0.7375 |
| 25pg *ASPHD1* RNA + 50pg *INO80E* RNA | 61 | 87.646 | 25.875 | 3.313 | 0.018 | 0.9982 |
| 50pg *MAPK3* RNA | 57 | 92.595 | 18.270 | 2.420 | 0.5697 | >0.9999 |
| 25pg *ASPHD1* RNA + 50pg *MAPK3* RNA | 63 | 92.178 | 21.360 | 2.691 | 0.4316 | >0.9999 |
| 50pg *YPEL3* RNA | 65 | 89.603 | 22.781 | 2.826 | 0.0802 | >0.9999 |
| 25pg *ASPHD1* RNA + 50pg *YPEL3* RNA | 76 | 90.375 | 22.758 | 2.611 | 0.0957 | >0.9999 |

### Supplementary Table S12. Reproductive phenotypes and estrous cyclicity in 16p11.2 mouse model

#### Reproductive traits in female 16p11.2Del/+, 16p11.2Dup/+ and 16p11.2+/+ mice (C57BL/6N background; Arbogast et al, PLoS Genet 2016)

| Mouse ID | Genotype | Cohort | DoB | Age (weeks) | Estrous stage | Body weight (g) | Left Ovary weight (mg) | Right Ovary weight (mg) | Ovaries weight (mg) | Ovaries, weight corrected | Uterus weight (mg) | Uterus, weight corrected |
|---|---|---|---|---|---|---|---|---|---|---|---|---|
| **Deletion cohort** | | | | | | | | | | | | |
| WT_1 | +/+ | 16pDel | 13.06.17 | 18 | D | 29.5 | 5.9 | 4.3 | 10.2 | 0.00035 | 49.7 | 0.00168 |
| WT_2 | +/+ | 16pDel | 13.06.17 | 18 | D | 24.9 | 5.6 | 5.8 | 11.4 | 0.00046 | 48.8 | 0.00196 |
| WT_3 | +/+ | 16pDel | 15.06.17 | 18 | D | 27.3 | 6.4 | 4 | 10.4 | 0.00038 | 50.2 | 0.00184 |
| DEL_1 | Del/+ | 16pDel | 13.06.17 | 18 | D | 18.2 | 5.4 | 6 | 11.4 | 0.00063 | 48.4 | 0.00266 |
| DEL_2 | Del/+ | 16pDel | 15.06.17 | 18 | DP | 20.2 | 6.7 | 4 | 10.7 | 0.00053 | 88.2 | 0.00437 |
| DEL_3 | Del/+ | 16pDel | 13.06.17 | 19 | D | 19.2 | 2.3 | 2.9 | 5.2 | 0.00027 | 41 | 0.00214 |
| **Duplication cohort** | | | | | | | | | | | | |
| DUP_1 | Dp/+ | 16pDup | 13.06.17 | 18 | D | 29.5 | 6.1 | 3.6 | 9.7 | 0.00033 | 58.3 | 0.00198 |
| DUP_2 | Dp/+ | 16pDup | 13.06.17 | 18 | D | 19.4 | 3.8 | 2.5 | 6.3 | 0.00032 | 50.1 | 0.00258 |
| DUP_3 | Dp/+ | 16pDup | 15.06.17 | 18 | D | 25.9 | 5 | 5.2 | 10.2 | 0.00039 | 56.7 | 0.00219 |
| WT_4 | +/+ | 16pDup | 13.06.17 | 19 | D | 22.4 | 4.6 | 4.5 | 9.1 | 0.00041 | 36.4 | 0.00163 |
| WT_5 | +/+ | 16pDup | 13.06.17 | 19 | D | 29.2 | 1.4 | 2.1 | 3.5 | 0.00012 | 30.8 | 0.00105 |
| WT_6 | +/+ | 16pDup | 15.06.17 | 19 | D | 24.2 | 4.4 | 4.5 | 8.9 | 0.00037 | 41 | 0.00169 |

| | Deletion cohort | | | | | Duplication cohort | | | | | Test |
|---|---|---|---|---|---|---|---|---|---|---|---|
| | Mean_WT | SD_WT | Mean_DEL | SD_DEL | P-value | Mean_WT | SD_WT | Mean_DUP | SD_DUP | P-value | |
| Body weight | 27.23 | 1.879 | 19.20 | 0.817 | 0.005 | 25.27 | 2.877 | 24.93 | 4.18 | 0.93 | T-test |
| Ovary weight (left) | 5.97 | 0.33 | 4.80 | 1.846 | 0.43 | 3.47 | 1.464 | 4.97 | 0.939 | 0.29 | T-test |
| Ovary weight (right) | 4.70 | 0.787 | 4.30 | 1.283 | 0.73 | 3.70 | 1.131 | 3.77 | 1.11 | 0.96 | T-test |
| Ovaries weight | 10.67 | 0.525 | 9.10 | 2.773 | 0.48 | 7.17 | 2.594 | 8.73 | 1.733 | 0.52 | T-test |
| Ovaries weight, corrected | 0.00039 | 0.00005 | 0.00048 | 0.00015 | 0.51 | 0.00030 | 0.00013 | 0.00035 | 0.00003 | 0.61 | |
| | | | | | | | | | | | T-test |
| Uterus weight | 49.57 | 0.579 | 59.20 | 20.727 | 0.55 | 36.07 | 4.171 | 55.03 | 3.549 | 0.008 | T-test |
| Uterus weight, corrected | 0.00183 | 0.00011 | 0.00305 | 0.00095 | 0.15 | 0.00146 | 0.00029 | 0.00225 | 0.00025 | 0.04 | T-test |

#### Reproductive traits in male 16p11.2Del/+, 16p11.2Dup/+ and 16p11.2+/+ mice (C57BL/6N background; Arbogast et al, PLoS Genet 2016)

| Mouse ID | Body weight (g) | AGD (cm) | Seminal vesicle (g) | Left Testis (g) | Right Testis (g) | Right Kidney (g) | Testes, weight corrected (g/g) | Testes, kidney corrected (g/g) | Kidney, weight corrected (g/g) | Seminal vesicle, weight corrected (g/g) | Seminal vesicle, kidney corrected (g/g) | Ano-genital distance, weight corrected (cm/g) | Ano-genital distance, kidney corrected (cm/g) | Sperm count | Sperm motility | Progressive | Sperm/Testis | Sperm/Kidney |
|---|---|---|---|---|---|---|---|---|---|---|---|---|---|---|---|---|---|---|
| **Deletion cohort** | | | | | | | | | | | | | | | | | | |
| DUP_1 | 34.62 | 1.45 | 0.200 | 0.076 | 0.076 | 0.248 | 0.0044 | 0.6129 | 0.0072 | 0.0058 | 0.81 | 0.000 | 5.85 | 128.85 | 0.2482 | 0.1450 | 847.70 | 519.56 |
| DUP_2 | 33.10 | 1.64 | 0.272 | 0.082 | 0.083 | 0.210 | 0.0050 | 0.7857 | 0.0063 | 0.0082 | 1.30 | 0.050 | 7.81 | 113.17 | 0.2640 | 0.1174 | 685.86 | 538.89 |
| DUP_3 | 30.77 | 1.51 | 0.294 | 0.085 | 0.078 | 0.163 | 0.0053 | 1.0000 | 0.0053 | 0.0096 | 1.80 | 0.049 | 9.26 | 95.60 | 0.2593 | 0.2227 | 586.48 | 586.48 |
| DUP_4 | 33.08 | 1.51 | 0.315 | 0.096 | 0.094 | 0.200 | 0.0057 | 0.9500 | 0.0060 | 0.0095 | 1.58 | 0.046 | 7.55 | 94.96 | 0.1863 | 0.0433 | 499.79 | 474.80 |
| DUP_5 | 38.25 | 1.49 | 0.361 | 0.100 | 0.093 | 0.217 | 0.0050 | 0.8894 | 0.0057 | 0.0094 | 1.66 | 0.039 | 6.87 | 102.50 | 0.2153 | 0.1747 | 531.09 | 472.35 |
| WT_1 | 33.24 | 1.98 | 0.253 | 0.078 | 0.073 | 0.228 | 0.0045 | 0.6623 | 0.0069 | 0.0076 | 1.11 | 0.060 | 8.68 | 77.23 | 0.2268 | 0.1598 | 511.46 | 338.73 |
| WT_2 | 30.86 | 1.81 | 0.289 | 0.089 | 0.085 | 0.193 | 0.0056 | 0.9016 | 0.0063 | 0.0094 | 1.50 | 0.059 | 9.38 | 87.81 | 0.1290 | 0.0591 | 504.66 | 454.97 |
| WT_3 | 35.50 | 1.89 | 0.369 | 0.100 | 0.100 | 0.197 | 0.0056 | 1.0152 | 0.0055 | 0.0104 | 1.87 | 0.053 | 9.59 | 124.24 | 0.2243 | 0.1753 | 621.22 | 630.68 |
| WT_4 | 31.00 | 1.74 | 0.289 | 0.090 | 0.090 | 0.219 | 0.0058 | 0.8219 | 0.0071 | 0.0093 | 1.32 | 0.056 | 7.95 | 97.56 | 0.1860 | 0.0943 | 542.02 | 445.49 |
| WT_5 | 34.67 | 1.75 | 0.273 | 0.098 | 0.100 | 0.198 | 0.0057 | 1.0000 | 0.0057 | 0.0079 | 1.38 | 0.050 | 8.84 | 102.73 | 0.1097 | 0.1007 | 518.81 | 518.81 |
| **Duplication cohort** | | | | | | | | | | | | | | | | | | |
| DEL_1 | 23.60 | 1.35 | 0.263 | 0.070 | 0.070 | 0.160 | 0.0059 | 0.8750 | 0.0068 | 0.0111 | 1.64 | 0.057 | 8.44 | 135.67 | 15.0667 | 5.1667 | 969.05 | 847.92 |
| DEL_2 | 27.15 | 1.76 | 0.386 | 0.081 | 0.078 | 0.180 | 0.0059 | 0.8833 | 0.0066 | 0.0142 | 2.14 | 0.065 | 9.78 | 107.00 | 17.1667 | 4.1000 | 672.96 | 594.44 |
| DEL_3 | 25.68 | 1.48 | 0.230 | 0.073 | 0.074 | 0.151 | 0.0057 | 0.9735 | 0.0059 | 0.0090 | 1.52 | 0.058 | 9.80 | 75.24 | 16.4667 | 10.9333 | 511.84 | 498.28 |
| DEL_4 | 22.00 | 1.53 | 0.233 | 0.063 | 0.060 | 0.156 | 0.0056 | 0.7885 | 0.0071 | 0.0106 | 1.49 | 0.070 | 9.81 | 65.43 | 33.5333 | 22.1333 | 531.95 | 419.42 |
| DEL_5 | 23.00 | 1.56 | 0.307 | 0.076 | 0.081 | 0.168 | 0.0068 | 0.9345 | 0.0073 | 0.0133 | 1.83 | 0.068 | 9.29 | 97.00 | 35.4000 | 20.0000 | 617.83 | 577.38 |
| WT_6 | 30.94 | 1.79 | 0.329 | 0.079 | 0.086 | 0.193 | 0.0053 | 0.8549 | 0.0062 | 0.0106 | 1.70 | 0.058 | 9.27 | 114.00 | 14.6000 | 10.7333 | 690.91 | 590.67 |
| WT_7 | 33.00 | 1.89 | 0.275 | 0.090 | 0.090 | 0.165 | 0.0055 | 1.0909 | 0.0050 | 0.0083 | 1.67 | 0.057 | 11.45 | 139.67 | 15.6667 | 4.4667 | 775.93 | 846.46 |
| WT_8 | 26.84 | 1.56 | 0.335 | 0.090 | 0.090 | 0.177 | 0.0067 | 1.0169 | 0.0066 | 0.0125 | 1.89 | 0.058 | 8.81 | 68.33 | 31.8000 | 22.6333 | 379.63 | 386.06 |
| WT_9 | 29.76 | 1.72 | 0.335 | 0.090 | 0.090 | 0.189 | 0.0060 | 0.9524 | 0.0064 | 0.0113 | 1.77 | 0.058 | 9.10 | 145.00 | 19.6000 | 10.7333 | 805.56 | 767.20 |
| WT_10 | 25.42 | 1.69 | 0.279 | 0.091 | 0.088 | 0.150 | 0.0070 | 1.1933 | 0.0059 | 0.0110 | 1.86 | 0.066 | 11.27 | 135.67 | 23.4667 | 8.3333 | 757.91 | 904.44 |

| | Deletion cohort | | | | | Duplication cohort | | | | | Test |
|---|---|---|---|---|---|---|---|---|---|---|---|
| | Mean_WT | SD_WT | Mean_DEL | SD_DEL | P-value | Mean_WT | SD_WT | Mean_DUP | SD_DUP | P-value | |
| Body weight | 29.19 | 3.068 | 24.29 | 2.091 | 0.018 | 33.05 | 2.101 | 33.96 | 2.763 | 0.574 | T-test |
| Ano-genital distance | 1.73 | 0.122 | 1.536 | 0.149 | 0.054 | 1.834 | 0.101 | 1.52 | 0.071 | 0.0005 | T-test |
| Seminal vesicle | 0.3106 | 0.031 | 0.2838 | 0.065 | 0.429 | 0.2946 | 0.044 | 0.2884 | 0.059 | 0.856 | T-test |
| Testis (left) | 0.0888 | 0.002 | 0.0726 | 0.008 | 0.003 | 0.091 | 0.009 | 0.0878 | 0.010 | 0.604 | T-test |
| Testis (right) | 0.088 | 0.005 | 0.0726 | 0.007 | 0.003 | 0.0896 | 0.011 | 0.0848 | 0.008 | 0.467 | T-test |
| Kidney (right) | 0.1748 | 0.018 | 0.163 | 0.011 | 0.244 | 0.0055 | 0.001 | 0.0051 | 0.000 | 0.277 | T-test |
| Testis, weight corrected | 0.0061 | 0.001 | 0.0060 | 0.000 | 0.752 | 0.8802 | 0.145 | 0.8476 | 0.154 | 0.739 | T-test |
| Testis, kidney corrected | 1.0217 | 0.129 | 0.8910 | 0.070 | 0.082 | 0.2070 | 0.016 | 0.2076 | 0.031 | 0.970 | T-test |
| Kidney, weight corrected | 0.0060 | 0.001 | 0.0067 | 0.001 | 0.087 | 0.0063 | 0.001 | 0.0061 | 0.001 | 0.687 | T-test |
| Seminal vesicle, weight corrected | 0.0107 | 0.002 | 0.0117 | 0.002 | 0.456 | 0.0089 | 0.001 | 0.0085 | 0.002 | 0.656 | T-test |
| Seminal vesicle, kidney corrected | 1.779 | 0.097 | 1.726 | 0.268 | 0.689 | 1.436 | 0.282 | 1.429 | 0.394 | 0.975 | T-test |
| Anogenital distance, weight corrected | 0.0595 | 0.004 | 0.0634 | 0.006 | 0.244 | 0.0556 | 0.004 | 0.0450 | 0.005 | 0.004 | T-test |
| Anogenital distance, kidney corrected | 9.982 | 1.271 | 9.422 | 0.593 | 0.398 | 8.888 | 0.646 | 7.467 | 1.259 | 0.055 | T-test |
| Sperm count | 120.533 | 31.469 | 96.067 | 27.663 | 0.228 | 97.914 | 17.652 | 107.015 | 14.236 | 0.396 | T-test |
| Sperm motility | 21.027 | 6.966 | 23.527 | 10.037 | 0.659 | 0.175 | 0.054 | 0.235 | 0.033 | 0.068 | T-test |
| Progressive | 11.380 | 6.792 | 12.467 | 8.304 | 0.827 | 0.118 | 0.048 | 0.141 | 0.067 | 0.555 | T-test |
| Sperm/testis | 681.987 | 174.183 | 660.725 | 184.257 | 0.856 | 539.632 | 47.729 | 630.183 | 140.663 | 0.210 | T-test |
| Sperm/kidney | 698.968 | 211.109 | 587.489 | 161.383 | 0.376 | 477.737 | 107.190 | 518.416 | 47.639 | 0.460 | T-test |

#### Estrous cyclicity in female 16p11.2Del/+, 16p11.2Dup/+ and 16p11.2+/+ mice (C57BL/6N background; Arbogast et al, PLoS Genet 2016)

| Mouse ID | Genotype | Cohort | DoB | Body weight at P21 (g) | Age at first estrous (days) | Body weight at first estrous (g) | Body weight at P97 (g) | Sum of days in Proestrous | Sum of days in Estrous | Sum of days in Metestrous | Sum of days in Diestrous | Number of cycles |
|---|---|---|---|---|---|---|---|---|---|---|---|---|
| **Deletion cohort** | | | | | | | | | | | | |
| WT_1 | +/+ | 16pDel | 13.06.17 | 9.9 | 28 | 15 | 24.3 | 6 | 12 | 7 | 7 | 6.5 |
| WT_2 | +/+ | 16pDel | 13.06.17 | 12 | 34 | 16.9 | 24 | 6 | 15 | 5 | 6 | 7 |
| WT_3 | +/+ | 16pDel | 13.06.17 | 10.2 | 33 | 17.9 | 27.8 | 7 | 12 | 4 | 9 | 6.5 |
| WT_4 | +/+ | 16pDel | 13.06.17 | 11.1 | 32 | 16 | 23.4 | 6 | 10 | 4 | 12 | 6.5 |
| WT_5 | +/+ | 16pDel | 13.06.17 | 10.2 | 32 | 16.8 | 21.9 | 7 | 10 | 5 | 10 | 5.5 |
| WT_6 | +/+ | 16pDel | 13.06.17 | 10 | 33 | 16.5 | 25.2 | 6 | 16 | 3 | 7 | 5.5 |
| WT_7 | +/+ | 16pDel | 13.06.17 | 7.9 | 34 | 16.2 | 21.9 | 5 | 8 | 6 | 13 | 6 |
| WT_8 | +/+ | 16pDel | 13.06.17 | 10.3 | 33 | 15.7 | 22.2 | 7 | 14 | 4 | 7 | 6.5 |
| WT_9 | +/+ | 16pDel | 15.06.17 | 10.7 | 25 | 13.4 | 24.7 | 6 | 13 | 5 | 8 | 7.5 |
| WT_10 | +/+ | 16pDel | 15.06.17 | 12.2 | 25 | 13.5 | 24.4 | 7 | 14 | 4 | 7 | 6.5 |
| DEL_1 | Del/+ | 16pDel | 13.06.17 | 6.8 | 41 | 12.5 | 17.5 | 5 | 13 | 6 | 8 | 7.5 |
| DEL_2 | Del/+ | 16pDel | 13.06.17 | 7.4 | 37 | 15 | 17 | 3 | 21 | 3 | 5 | 6 |
| DEL_3 | Del/+ | 16pDel | 13.06.17 | 7.7 | 40 | 13.7 | 17.4 | 7 | 13 | 4 | 8 | 6.5 |
| DEL_4 | Del/+ | 16pDel | 13.06.17 | 8.7 | 36 | 14.3 | 19.9 | 6 | 17 | 3 | 6 | 6.5 |
| DEL_5 | Del/+ | 16pDel | 13.06.17 | 8.8 | 38 | 14.4 | 17.5 | 6 | 18 | 3 | 5 | 5.5 |
| DEL_6 | Del/+ | 16pDel | 15.06.17 | 9.8 | 33 | 15.8 | 19.5 | 6 | 15 | 5 | 6 | 6 |
| **Duplication cohort** | | | | | | | | | | | | |
| WT_1 | +/+ | 16pDup | 13.06.17 | 8.8 | 40 | 15.8 | 19.6 | 5 | 10 | 9 | 8 | 6.5 |
| WT_2 | +/+ | 16pDup | 13.06.17 | 7.3 | 42 | 14.4 | 26.7 | 6 | 13 | 7 | 6 | 6 |
| WT_3 | +/+ | 16pDup | 13.06.17 | 11.5 | 31 | 16.2 | 19.7 | 13 | 9 | 2 | 8 | 3 |
| WT_4 | +/+ | 16pDup | 13.06.17 | 10.5 | 36 | 17 | 23.6 | 5 | 11 | 7 | 9 | 6 |
| WT_5 | +/+ | 16pDup | 13.06.17 | 8.6 | 36 | 17.5 | 22.7 | 6 | 8 | 6 | 12 | 6.5 |
| WT_6 | +/+ | 16pDup | 15.06.17 | 9 | 33 | 16.3 | 26.8 | 7 | 11 | 6 | 8 | 6.5 |
| WT_7 | +/+ | 16pDup | 15.06.17 | 9.1 | 38 | 16.9 | 24 | 7 | 14 | 5 | 6 | 5.5 |
| DUP_1 | Dp/+ | 16pDup | 13.06.17 | 7 | 39 | 15.2 | 28.2 | 6 | 13 | 7 | 6 | 6 |
| DUP_2 | Dp/+ | 16pDup | 13.06.17 | 7.9 | 34 | 16.1 | 25 | 7 | 15 | 3 | 7 | 6 |
| DUP_3 | Dp/+ | 16pDup | 13.06.17 | 7.9 | 39 | 14.8 | 20.7 | 4 | 14 | 5 | 9 | 6 |
| DUP_4 | Dp/+ | 16pDup | 13.06.17 | 9.9 | 35 | 15.9 | 20.1 | 4 | 10 | 5 | 13 | 5.5 |
| DUP_5 | Dp/+ | 16pDup | 13.06.17 | 9.2 | 33 | 16.6 | 26.8 | 12 | 6 | 5 | 9 | 4 |
| DUP_6 | Dp/+ | 16pDup | 15.06.17 | 9.9 | 32 | 16.7 | 23.8 | 7 | 16 | 3 | 6 | 6 |
| DUP_7 | Dp/+ | 16pDup | 15.06.17 | 10.4 | 34 | 17.5 | 22.2 | 5 | 15 | 7 | 5 | 6 |
| DUP_8 | Dp/+ | 16pDup | 15.06.17 | 9.3 | 32 | 17.1 | 23 | 10 | 10 | 4 | 8 | 6 |
| DUP_9 | Dp/+ | 16pDup | 15.06.17 | 9.6 | 38 | 18.8 | 22.7 | 7 | 8 | 4 | 13 | 5 |

| | **Deletion cohort** | | | | | **Duplication cohort** | | | | | Test |
|---|---|---|---|---|---|---|---|---|---|---|---|
| | Mean_WT | SD_WT | Mean_DEL | SD_DEL | P-value | Mean_WT | SD_WT | Mean_DUP | SD_DUP | P-value | |
| Body weight at P21 | 10.45 | 1.145 | 8.20 | 1.002 | 0.001 | 9.257 | 1.261 | 9.011 | 1.080 | 0.350 | T-test |
| Body weight at first estrous | 15.790 | 1.379 | 14.283 | 1.029 | 0.048 | 16.300 | 0.938 | 16.522 | 1.145 | 0.703 | T-test |
| Body weight at P97 | 23.980 | 1.704 | 18.133 | 1.126 | 6.2708E-06 | 22.757 | 2.039 | 24.033 | 2.857 | 0.828 | T-test |
| | **Deletion cohort** | | | | | **Duplication cohort** | | | | | Test |
| | Mean_WT | SD_WT | Mean_DEL | SD_DEL | P-value | Mean_WT | SD_WT | Mean_DUP | SD_DUP | P-value | |
| Days in Proestrous | 6.30 | 0.640 | 5.5 | 1.258 | 0.229 | 7 | 2.563 | 6.9 | 2.514 | 0.937 | T-test |
| Days in Estrous | 12.4 | 2.375 | 16.2 | 2.853 | 0.034 | 10.9 | 1.959 | 11.9 | 3.315 | 0.480 | T-test |
| Days in Metestrous | 4.7 | 1.100 | 4.0 | 1.155 | 0.295 | 6.0 | 2.000 | 4.8 | 1.397 | 0.229 | T-test |
| Days in Diestrous | 8.6 | 2.245 | 6.3 | 1.247 | 0.029 | 8.1 | 1.884 | 8.4 | 2.753 | 0.811 | T-test |
| Total number of cycles | 6.4 | 0.583 | 6.3 | 0.624 | 0.849 | 5.7 | 1.161 | 5.6 | 0.657 | 0.849 | T-test |
